# Temperature alters gene expression in mosquitoes during arbovirus infection

**DOI:** 10.1101/2020.07.31.230425

**Authors:** BMC Randika Wimalasiri-Yapa, Roberto A. Barrero, Liesel Stassen, Louise M. Hafner, Elizabeth A. McGraw, Alyssa T. Pyke, Cassie C. Jansen, Andreas Suhrbier, Laith Yakob, Wenbiao Hu, Gregor J. Devine, Francesca D. Frentiu

## Abstract

Arthropod-borne viruses (arboviruses) such as dengue, Zika and chikungunya constitute a significant proportion of the global disease burden. The principal vector of these pathogens is the mosquito *Aedes* (*Ae*.) *aegypti*, and its ability to transmit virus to a human host is influenced by environmental factors such as temperature. However, exactly how ambient temperature influences virus replication within mosquitoes remains poorly elucidated, particularly at the molecular level. Here, we use chikungunya virus (CHIKV) as a model to understand how the host mosquito transcriptome responds to arbovirus infection under different ambient temperatures. We exposed CHIKV-infected mosquitoes to 18 °C, 28 °C and 32 °C, and found higher temperature correlated with higher virus replication levels, particularly at early time points post-infection. Lower ambient temperatures resulted in reduced virus replication levels. Using RNAseq, we found that temperature significantly altered gene expression levels in mosquitoes, particularly components of the immune response. The highest number of significantly differentially expressed genes in response to CHIKV was observed at 28 °C, with a markedly more muted effect observed at either lower (18 °C) or higher (32 °C) temperatures. At the higher temperature, the expression of many classical immune genes, including *Dicer-2* in the RNAi pathway, was not substantially altered in response to CHIKV. Upregulation of Toll, IMD and JAK-STAT pathways was only observed at 28 °C. Time post infection also led to substantially different gene expression profiles, and this effect varied depending upon the which temperature mosquitoes were exposed to. Taken together, our data indicate temperature significantly modulates mosquito gene expression in response to infection, potentially leading to impairment of immune defences at higher ambient temperatures.

## INTRODUCTION

Arthropod-borne diseases constitute a significant proportion of the global infectious disease burden, with yearly estimates of ~ 1 billion infections and 1 million deaths [1]. Dengue viruses (DENVs 1-4), Zika virus (ZIKV) and chikungunya (CHIKV) are some of the most common pathogens causing epidemics of arthropod-borne virus (arboviruses) disease in recent decades [2]. These viruses are principally vectored to humans by the mosquitoes *Aedes (Ae.) aegypti* and *Ae. albopictus* [3]. Because mosquitoes are poikilothermic, almost all their biological activities are influenced by ambient environmental conditions [4], such as temperature. Understanding how mosquitoes respond to changes in this key parameter, over the short and long term, are necessary to improve predictions of arbovirus futures.

Recent projections have suggested that climate change may increase the risk of arbovirus transmission, as higher average temperatures are projected to expand the geographic distributions and lengthen active seasons of arthropod vectors [5–9] Temperature is also known to alter the ability of *Aedes* spp. mosquitoes to transmit viruses, with higher temperatures following infection leading to increased viral replication and earlier transmission potential [10, 11]. Conversely, lower ambient temperatures lead to decreased viral replication and delayed transmission by mosquitoes [12, 13]. Exactly how ambient temperature influences the physiological, molecular and genetic interactions between virus and mosquito during infection remains poorly elucidated.

Mosquitoes possess physical and physiological barriers against pathogen infection. Insect protection relies on the humoral and cellular immune responses, which comprise the innate immune system [14]. Mosquito immune response can be divided into four components: pathogen recognition, activation of immune signalling, immune effector mechanisms [15] and immune modulation by the regulation of mosquito homeostasis. Apart from known (termed ‘classical’) immune genes within these four categories, recent studies have indicated the involvement of additional gene families during arbovirus infection in *Ae. aegypti*. Additional genes included cytoskeleton and cellular trafficking, heat shock response, cytochrome P450, cell proliferation, chitin and small RNAs [15]. Long noncoding RNAs (lncRNAs) are also involved in *Ae. aegypti*-virus interactions, particularly in DENV and ZIKV infections [16, 17]. However, it is not well understood how gene expression is modulated in mosquitoes exposed to different ambient temperatures. Even less well understood is how the mosquito immune response may change in response to temperature during prolonged virus infection, and how this may impact on the insect’s ability to vector pathogens.

CHIKV is an emerging arbovirus, responsible for several recent large outbreaks globally with an estimated 10 million cases [18–20]. Rarely fatal, CHIKV typically causes an acute febrile syndrome and severe, debilitating rheumatic disorders in humans that may persist for months [21]. The main vector of CHIKV is *Ae. aegypti* [22], with *Ae. albopictus* also playing an important role in more recent outbreaks [23]. CHIKV cases were reported from Africa, Asia, Europe, islands of Indian and Pacific oceans until 2013, when the first autochthonous cases were reported in the Americas on islands in the Caribbean. By the end of 2017, more than 2.6 million suspected cases of CHIKV had been reported from the Caribbean and the Americas [24]. Since then, the virus has continued to circulate and cause sporadic disease and periodic outbreaks with very high attack rates in many areas of the world [24]. CHIKV is a single-stranded, positive sense RNA virus from the *Togaviridae* family, genus *Alphavirus*. To date, the interactions between emerging alphaviruses, such as CHIKV, and their mosquito hosts under different temperatures have been less characterised than those of flaviviruses, namely DENV and ZIKV. In addition, although mosquitoes remain infected with arboviruses for life [25], the immune responses underlying long term persistence of these pathogens is not well understood.

Here, we report how exposure to 3 ambient temperature regimes (18 °C, 28 °C and 32 °C) alter gene expression in mosquitoes infected with CHIKV, sampled at two time points post infection. We found that temperature alters the transcriptome, with the highest number of upregulated genes observed at 28 °C, while lower temperatures were associated with more downregulated genes. Importantly, we observed the absence of *Dicer-2* and low levels of immune gene expression at 32 °C, suggesting heat may impair mosquito immunity and the ability to mount an adequate RNAi response. We also show that mosquito gene expression is not constant over the course of arbovirus infection, with distinct gene expression profiles observed for each temperature and time point sampled.

## RESULTS

### CHIKV replication varies with ambient temperature and age of mosquitoes

To determine how ambient temperature modulates replication of CHIKV, and how this may vary over the course of infection, *Ae. aegypti* mosquitoes were orally infected with a CHIKV strain from the Asian genotype (GenBank accession MF773560) and held at 18, 28 or 32 °C for either 3- or 7-days post infection (dpi) (n=18-20 mosquitoes per combination of T °C and dpi). Using qRT-PCR (as per [26]), we found that at 3 dpi, the number of virus genome copies in mosquito bodies (including heads but without wings and legs) held at 32 °C was significantly higher than that from mosquitoes held at the other temperatures (**Fig. 1 A**). By contrast, at 7 dpi, the highest amount of virus was present in mosquitoes held at 28 °C, with a decline in CHIKV copy number observed at 32 °C (**Fig. 1 B**). Next, we performed RNAseq on a subset of six mosquitoes for each temperature and timepoint (n=72 total). Of the 2,661,279,246 Illumina paired end reads generated in this study, 10,155,660 (0.38%) of reads mapped onto the CHIKV genome (MF773560 strain). CHIKV reads obtained from the RNAseq data were largely congruent with qRT-PCR results (**Fig. 1 C**), with the highest read count (normalized reads per million) observed in mosquitoes held at 32 °C and sampled at 3 dpi. Normalized viral read counts were significantly correlated with copy number from qRT-PCR (r^2^=0.8916, p<0.001). Two samples, from a total of 72 submitted for RNAseq, were removed from downstream analyses of differential gene expression. These samples were identified as outliers in the correlation between virus titer and normalised read counts (**Fig. 1 C**).

**Fig. 1.**
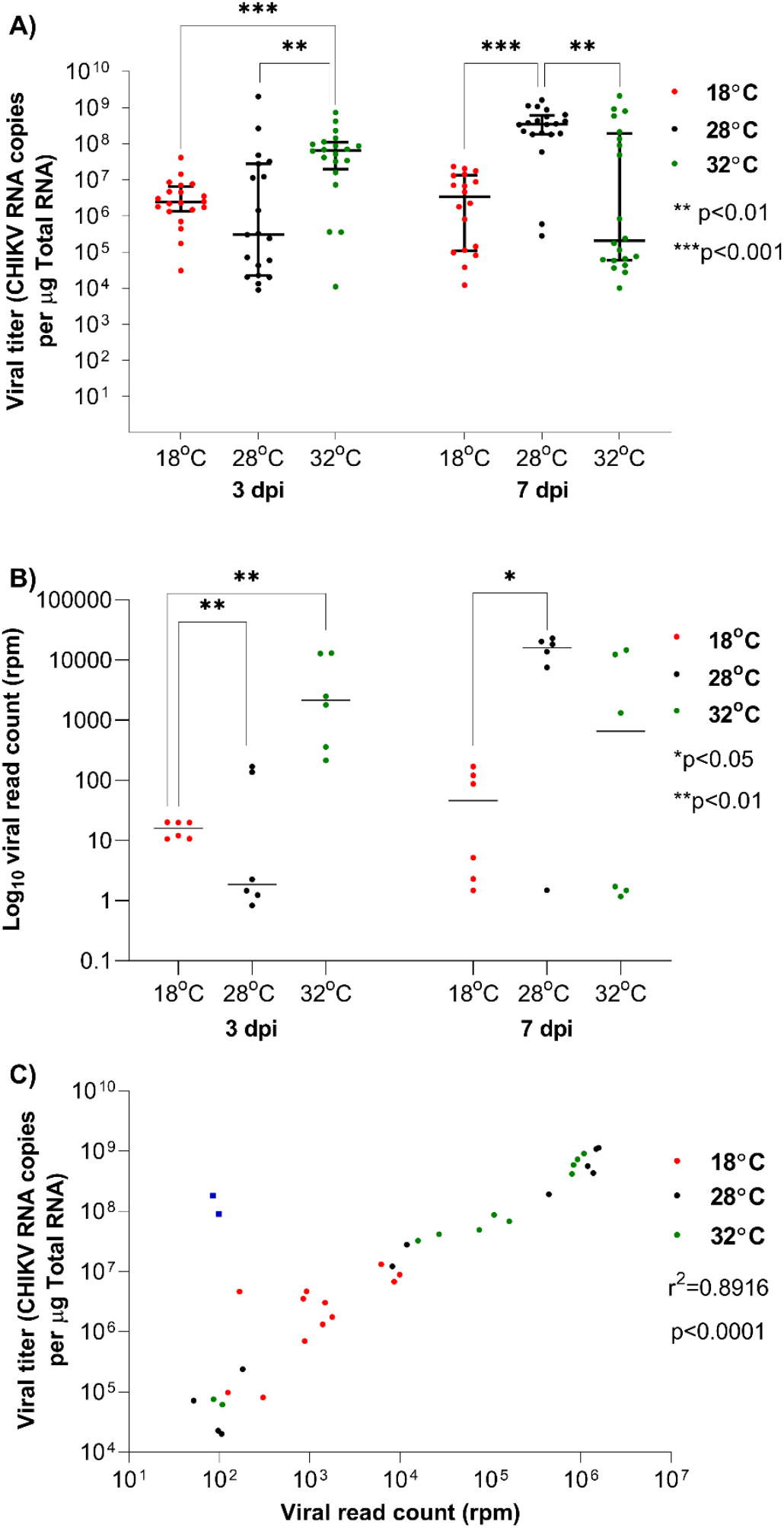
Effect of temperature and day post infection (dpi) on CHIKV replication in *Ae. aegypti* mosquitoes. **A**) CHIKV RNA copy numbers detected using qRT-PCR in whole mosquito bodies, 18 °C (n=20 and n=18), 28 °C (n=19 and n=20) and 32 °C (n=20 in each timepoint), sampled at 3 and 7 dpi. Statistical significance was assessed using Mann Whitney tests. **B**) CHIKV read counts obtained from RNAseq of *Ae. aegypti* samples (6 mosquito bodies per each temperature/dpi combination). Log10 normalised reads per million (rpm) counts shown. **C**) Pearson correlation between body CHIKV titer from qRT-PCR and virus read counts, across all temperatures and both time points. Each point on the plots represents an individual mosquito. The dark blue squares in C) indicate two outlier samples that were removed from downstream analyses of the mosquito transcriptome.

### Temperature alters differential gene expression during CHIKV infection

A total of 2,511,065,774 (94.36%) reads from 70 samples were mapped to the reference genome of *Ae. aegypti*, version AaegL5.2 (**Supplementary Table 1**). For each temperature and time regime, differentially expressed genes (DEGs) in response to CHIKV infection were identified using DEseq2 by comparison to uninfected control mosquitoes. DEGs were considered statistically significant if the adjusted p-value <0.05 and absolute fold change (FC) > ± 1.5. At 3 dpi, we detected 715 DEGs across all temperature regimes. We observed a large number of upregulated genes (n=374) in mosquitoes held at 28 °C, but a dramatically lower number at 32 °C, with classical immune gene expression generally following the same pattern (**Fig 2 A; Supplementary Table 2** for a list of genes). At 7 dpi, we found a similar number of DEGs (n=726) in response to CHIKV infection to that observed at 3 dpi (**Fig 2 B**). The highest number of DEGs was observed in mosquitoes held at 28 °C, including classical immune response genes, a pattern similar to 3 dpi (**Supplementary Table 3**). By contrast, at 7 dpi we observed a substantial decrease in DEGs at 18 °C but a marked increase in upregulated DEGs at 32 °C.

**Fig 2.**
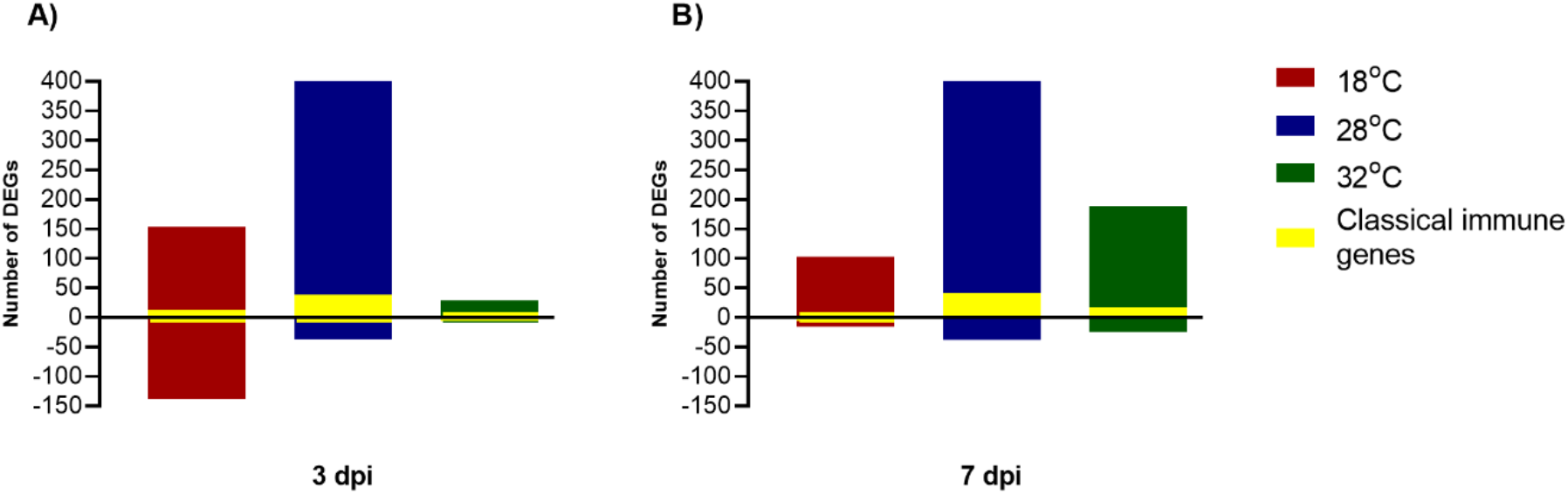
Differentially expressed genes (DEGs) in response to CHIKV infection in *Ae. aegypti*. Mosquitoes were held at 3 ambient temperatures and sampled at **A**) 3 dpi and **B**) 7 dpi. DEGs were identified using DESeq2, with a Fold Change (FC) > ±1.5 and adjusted p-value < 0.05. The number of DEGs involved in the classical immune response is shown in yellow.

Across all temperatures and time points, the top 10 upregulated (in terms of fold-change) DEGs were predominantly genes known to be involved in thermoregulatory responses, such as heat shock proteins (particularly *hsp70*) and lethal (2) essential for life protein (*l2efl*) (**Table 1**). By contrast, the top 10 downregulated DEGs were more heterogeneous across all temperatures and both time points. At 3 dpi, proteolysis, intracellular signal transduction, protein-binding and oxidation-reduction process related genes were among the top 10 downregulated genes at 18 °C. At 28 °C, downregulated genes were predominantly involved in oxidation-reduction, cell division, zinc ion binding, nucleic acid binding and integral component of membrane (**Table 1**). Four genes downregulated at 32 °C were related to oxidation-reduction and protein binding. At 7 dpi, RNA binding, pigment binding, lipid binding and transport, proteolysis and integral component of membrane gene were among the top 10 downregulated genes at 18 °C (**Table 1**). Microtubule binding, catalytic activity, odorant binding, membrane and ionotropic glutamate receptor activity genes were among the most downregulated DEGs at 28 °C, while odorant binding, nucleotide binding, ATP binding, phototransduction and transferase genes were downregulated at 32 °C. Across all temperature regimes and dpi, we observed 11 lncRNAs being among the top 10 downregulated, but no lncRNAs were present among the top 10 upregulated genes (**Table 1**). At 3 dpi, 14 genes were upregulated under all three temperature regimes in response to CHIKV infection, and comprised 10 HSPs, nucleic acid binding and genes involved in regulation of cell cycle (**Table 2**). By 7 dpi, 22 genes that were upregulated across all temperature regimes (**Table 2**) with a similar pattern observed in the categories of genes involved. There were no downregulated DEGs in common across the three temperatures, for either time points. Overall, the results indicate similar categories of upregulated genes, but far greater heterogeneity of downregulated DEGs, among the temperature regimes in response to CHIKV infection.

**Table 1.**
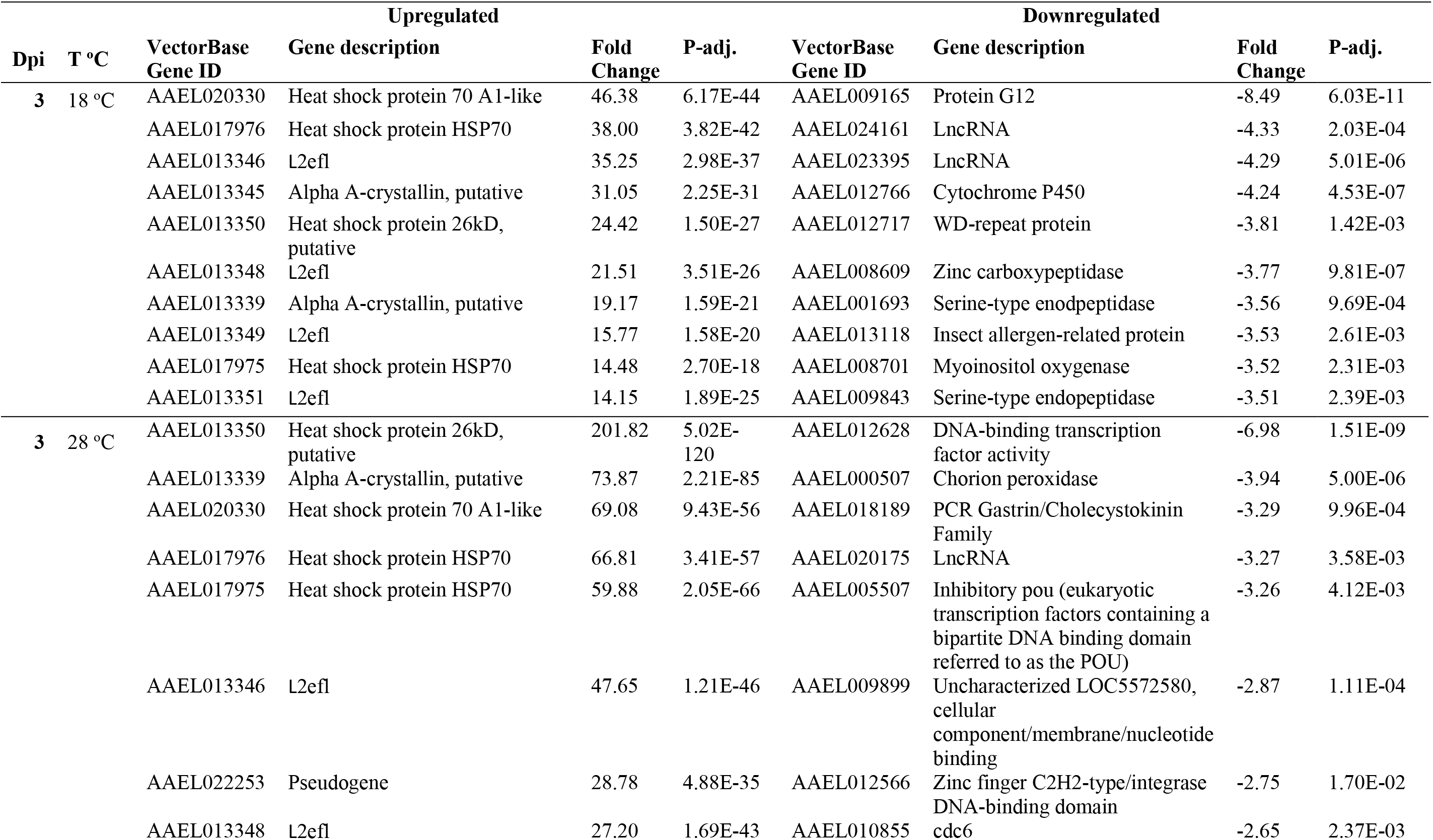
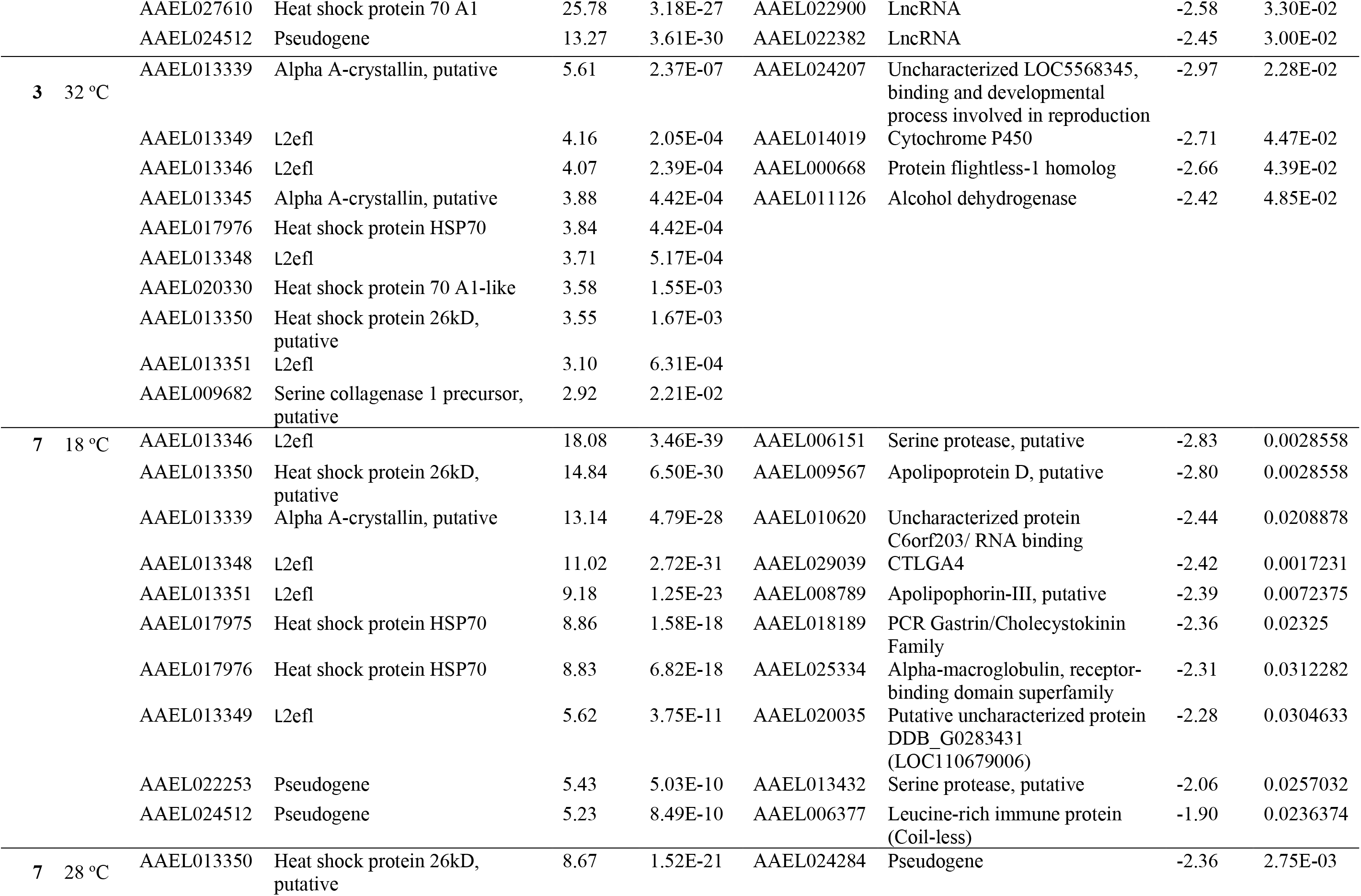
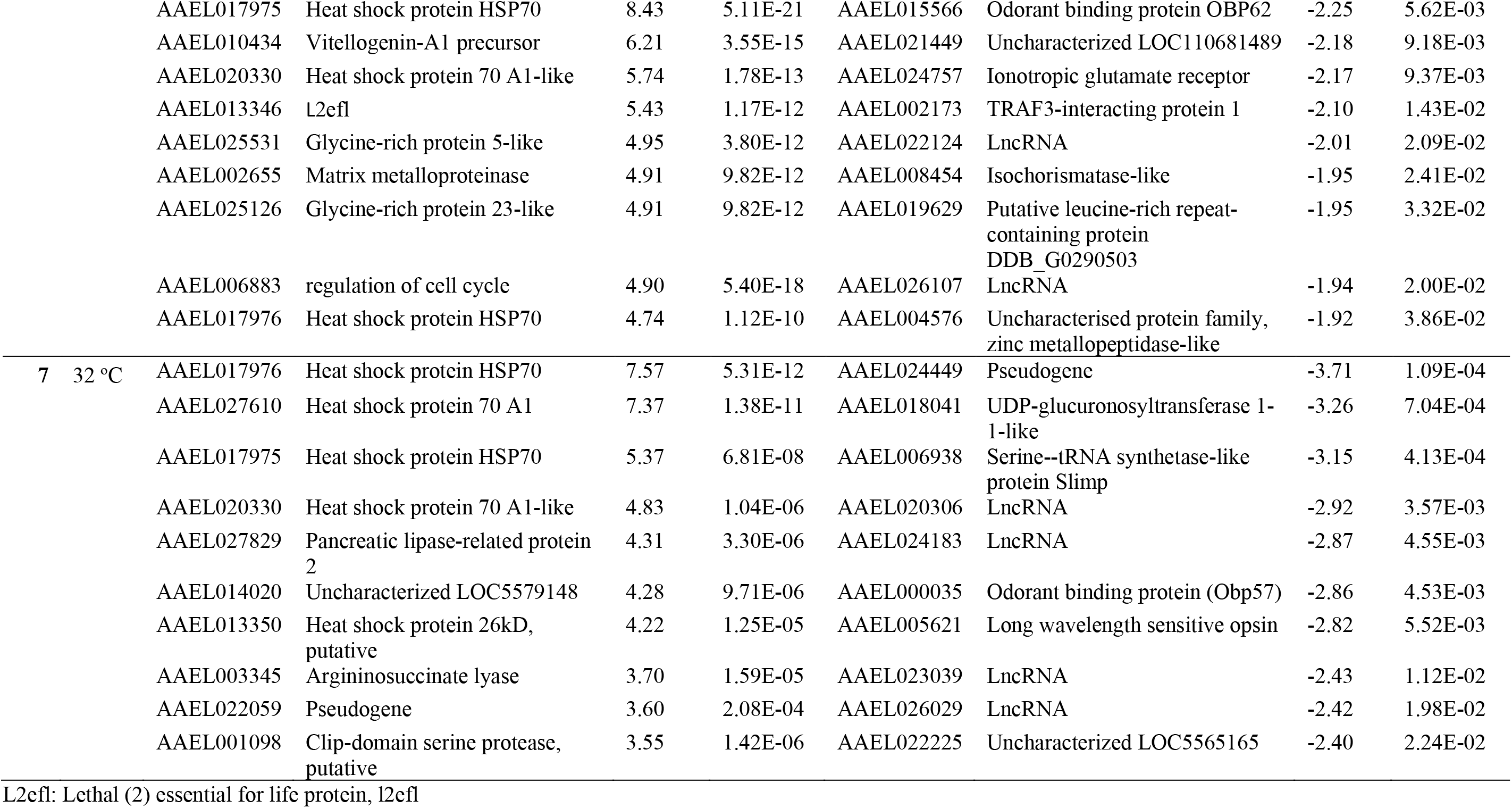
Top 10 differentially expressed genes (DEGs) ranked by fold change in CHIKV-infected *Ae. aegypti* versus uninfected controls, held at three different temperatures (T°C) and sampled at two time points post infection (dpi).

**Table 2.**
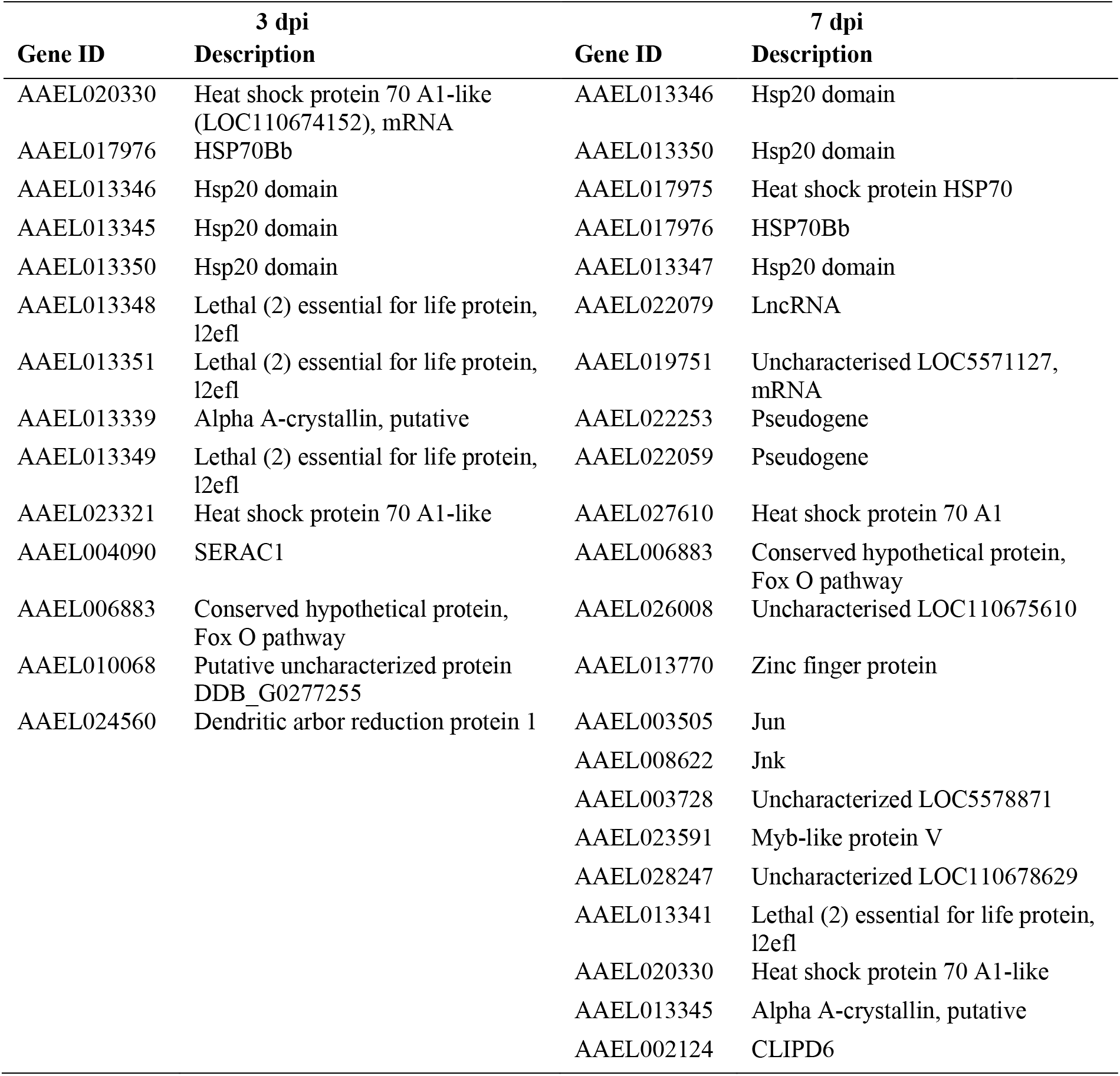
Genes observed to be differentially expressed in common across the three temperatures, in CHIKV-infected *Ae. aegypti* sampled at two time points post infection (dpi).

| 3 dpi |  | 7 dpi |  |
| --- | --- | --- | --- |
| Gene ID | Description | Gene ID | Description |
| AAEL020330 | Heat shock protein 70 A1-like (LOC110674152), mRNA | AAEL013346 | Hsp20 domain |
| AAEL017976 | HSP70Bb | AAEL013350 | Hsp20 domain |
| AAEL013346 | Hsp20 domain | AAEL017975 | Heat shock protein HSP70 |
| AAEL013345 | Hsp20 domain | AAEL017976 | HSP70Bb |
| AAEL013350 | Hsp20 domain | AAEL013347 | Hsp20 domain |
| AAEL013348 | Lethal (2) essential for life protein, l2efl | AAEL022079 | LncRNA |
| AAEL013351 | Lethal (2) essential for life protein, l2efl | AAEL019751 | Uncharacterised LOC5571127, mRNA |
| AAEL013339 | Alpha A-crystallin, putative | AAEL022253 | Pseudogene |
| AAEL013349 | Lethal (2) essential for life protein, l2efl | AAEL022059 | Pseudogene |
| AAEL023321 | Heat shock protein 70 A1-like | AAEL027610 | Heat shock protein 70 A1 |
| AAEL004090 | SERAC1 | AAEL006883 | Conserved hypothetical protein, Fox O pathway |
| AAEL006883 | Conserved hypothetical protein, Fox O pathway | AAEL026008 | Uncharacterised LOC110675610 |
| AAEL010068 | Putative uncharacterized protein DDB_G0277255 | AAEL013770 | Zinc finger protein |
| AAEL024560 | Dendritic arbor reduction protein 1 | AAEL003505 | Jun |
|  |  | AAEL008622 | Jnk |
|  |  | AAEL003728 | Uncharacterized LOC5578871 |
|  |  | AAEL023591 | Myb-like protein V |
|  |  | AAEL028247 | Uncharacterized LOC110678629 |
|  |  | AAEL013341 | Lethal (2) essential for life protein, l2efl |
|  |  | AAEL020330 | Heat shock protein 70 A1-like |
|  |  | AAEL013345 | Alpha A-crystallin, putative |
|  |  | AAEL002124 | CLIPD6 |

### Identification of *Ae. aegypti* genes involved in classical and non-classical immune response and updating annotations of AaegL5.2

To determine how the expression of known mosquito immune genes changes with temperature in response to CHIKV infection, we first identified from ImmunoDB database 445 genes listed under 27 families that were similarly annotated in AaegL1 and AaegL5.2. This indicated that 166 genes were no longer available with the same gene ID in the latest genome annotation release AaegL5.2. Since the release of AaegL1, a number of additional genes have been identified to be involved in mosquito immune response. Using a literature review, we identified 10 additional gene families, resulting in a total of 37 gene families implicated in mosquito immunity (**Supplementary Table 4**), producing a final list of 998 genes. The list was divided into classical or non-classical components (**Supplementary Tables 5 and 6**) [4, 27–34]. Genes already identified and directly involved in humoral and cellular immune response were considered as classical immune genes [35]. Genes outside of these classical immune pathways that are transcriptionally altered in response to arboviral infections in *Ae. aegypti* were considered as non-classical immune genes [36]. We next considered DEG patterns in classical immune genes, categorised under the four main processes of immune response: pathogen recognition, immune signalling, pathogen destruction and immune gene modulation.

### Pathogen recognition receptor (PRR) genes

At both time points sampled post CHIKV infection, the majority of PRR DEGs were observed in mosquitoes held at 28 °C (n DEGs=24; **Fig. 3**), with only a handful detected at the other temperatures. The number of PRR DEGs observed at 28 °C stayed largely similar at both time points (3 dpi n=24; 7 dpi n=30). Strikingly, only two PRRs were found to be differentially expressed at 32 °C at 3 dpi (**Fig. 3A**), although by 7 dpi that number had increased 6-fold (**Fig. 3B**). By contrast, the number of PRRs differentially expressed at 18 °C (n=8) was halved by 7 dpi. PRR DEGs were predominantly CLIP domain serine proteases, leucine-rich immune receptors (LRIM) and leucine-rich repeat-containing proteins (LRR), galectins, fibrinogen related proteins (FREP), ML/Niemann-pick receptors, peptidoglycan recognition proteins (PGRP), scavenger receptors (SCR), and thioester proteins (TEP). At 3 dpi, two CLIPs (AAEL003632: *CLIPB39*, AAEL012712: *CLIPC13*) and one TEP (AAEL008607: *TEP3*) were found in common to mosquitoes held at 18 °C and at 28 °C. At 7 dpi, we observed *CLIPD6* (AAEL002124) to be upregulated under all temperatures.

**Fig. 3.**
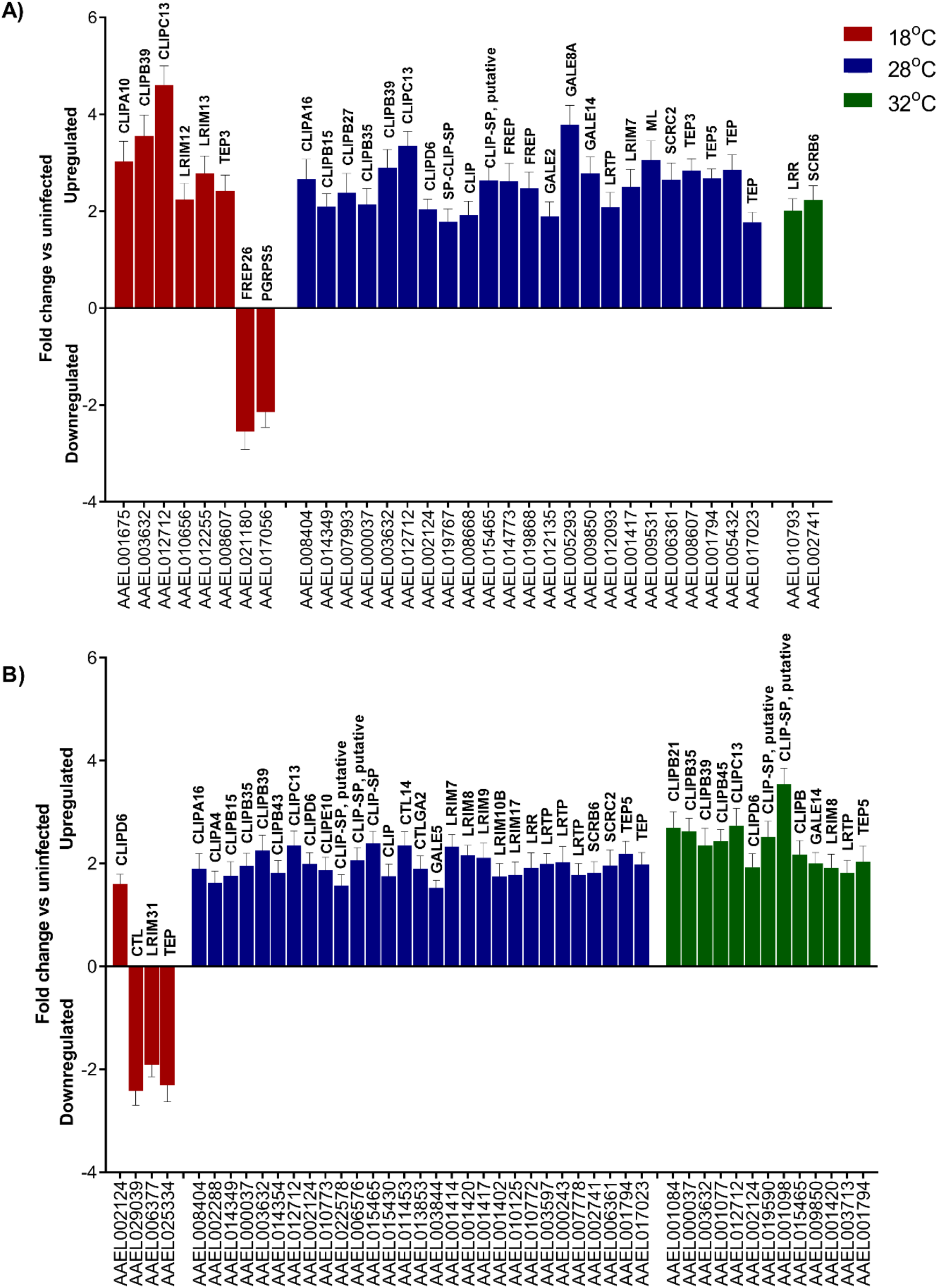
Differentially expressed genes (DEGs) related to pathogen recognition at **A**) 3 dpi and **B**) 7 dpi, in CHIKV-infected *Ae. aegypti* versus uninfected controls. CTL: C-type lectin; LRIM: leucine-rich immune receptors; LRR: leucine-rich repeat-containing proteins; LRTP: leucine-rich transmembrane protein; FREP: fibrinogen related proteins; ML: Niemann-Pick receptors; GALE: galectin; PGRP: peptidoglycan recognition proteins; SCR: scavenger receptors; TEP: thioester proteins.

### Immune signalling genes

We found a complete absence of immune signalling DEGs at 32 °C at the 3 dpi, with only two DEGs detected by 7 dpi (**Fig. 4**). Upregulation of *Cactus* (AAEL000709), *Toll 5A* (AAEL007619) and downregulation of *TRAF6* (AAEL028236) were only found at 18 °C (**Fig. 4**). *Kayak* (AAEL008953) and *AP-1* (AAEL003505) were expressed at both time points at 18 °C and 28 °C, but only *AP-1* was observed at 32 °C at 7 dpi. The repertoire of immune signalling genes was stable across the two time points at 28 °C and included two *Spätzle* genes (AAEL001929 and AAEL013434), *Relish 2* (AAEL007624) gene and a hypothetical protein from JAK-STAT pathway (AAEL009645).

**Fig. 4.**
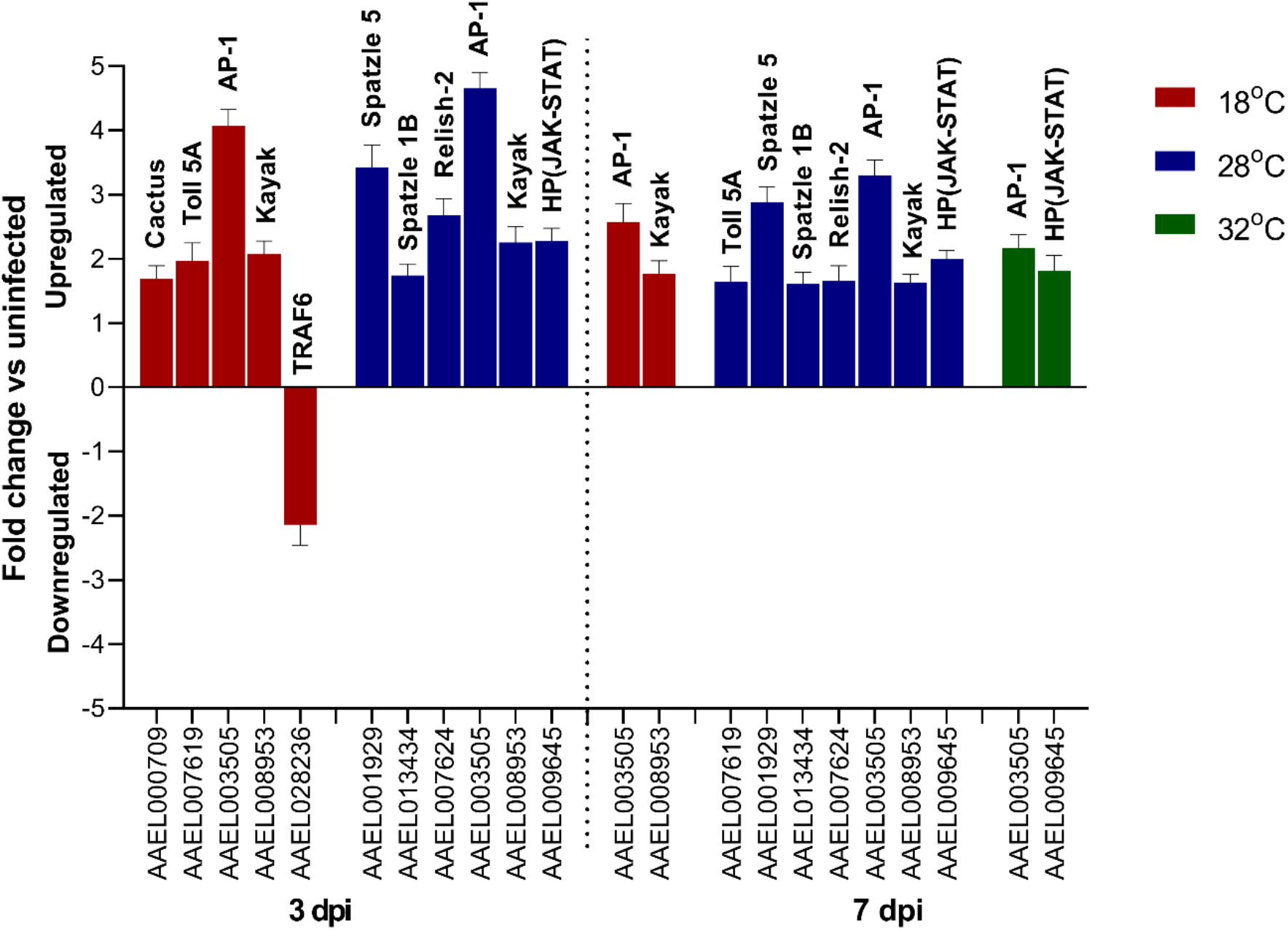
Differentially expressed genes (DEGs) related to immune signalling at **A**) 3 dpi and **B**) 7 dpi, in CHIKV-infected *Ae. aegypti* versus uninfected controls. HP indicates hypothetical protein.

### Pathogen destruction and immune modulation genes

There were no pathogen destruction or related effector DEGs in mosquitoes held at 32 °C at 3 dpi, although *Caspase 8* (AAEL014348) was upregulated at 7 dpi (**Fig. 5A**). *Caspase* 8 and *Dicer-2* (AAEL006794) were upregulated at 18 °C at 3 dpi but were not differentially expressed at 7 dpi. Upregulation of *Dicer-2* expression in response to CHIKV was only observed at 18 °C and 28 °C. In contrast, the mosquitoes kept at 28 °C differentially expressed most of the previously identified pathogen destruction mechanisms including antimicrobial peptide (*AMP*) genes (*Attacin*, *Defensin* and *Cecropin*) and apoptosis genes (*Caspase 8* and *IAP 1*). By 7 dpi, most expression of DEGs was suppressed at 18 °C and reduced at 28 °C. We did not observe any immune modulation genes significantly upregulated at 32 °C at 3 dpi, although by 7 dpi two serpins were observed (**Fig. 5 B**). Conversely, we observed three immune modulation DEGs (AAEL003182: *SRPN26*; AAEL010235: allergen; AAEL006347: *apyrase*) downregulated at 18 °C at 3 dpi but none at 7 dpi.

**Fig. 5.**
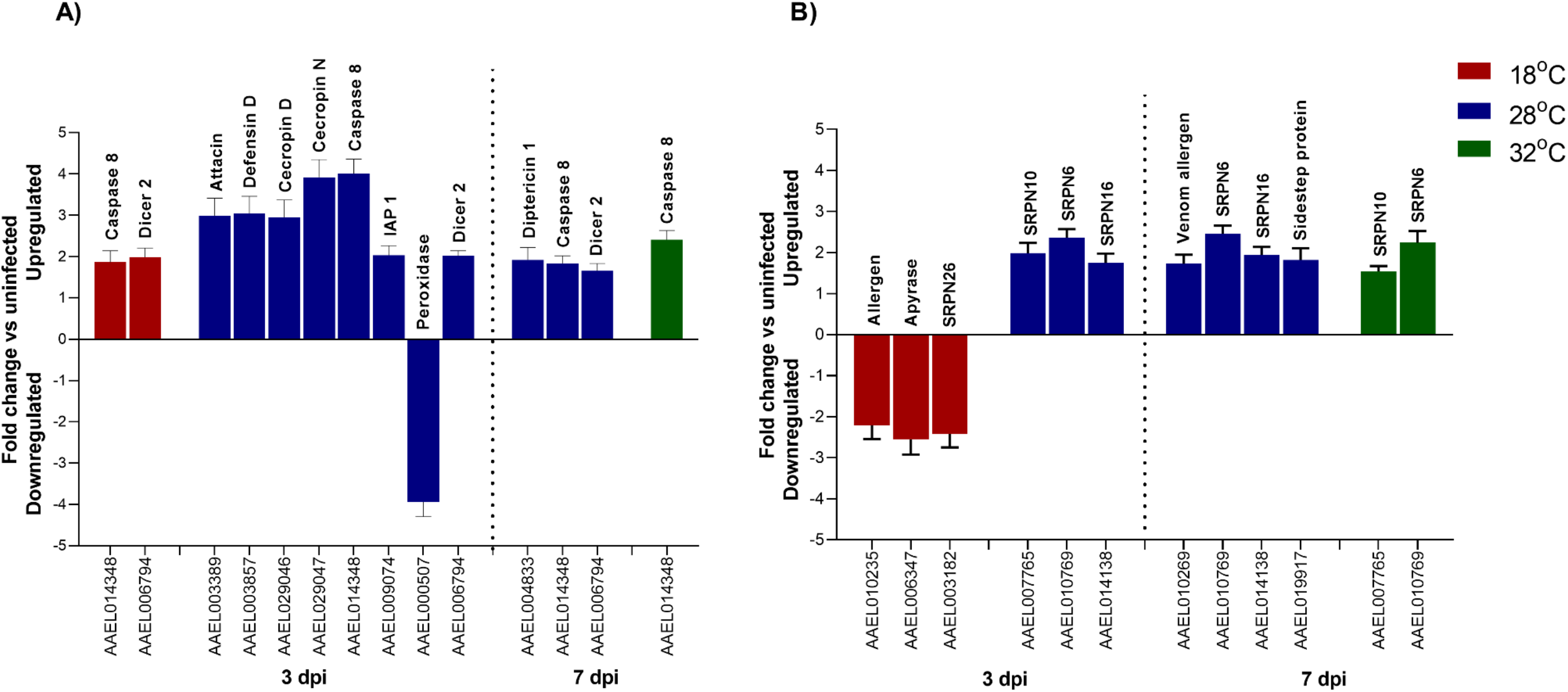
Differentially expressed genes (DEGs) related to **A**) pathogen destruction and **B**) immune modulation, in CHIKV-infected *Ae. aegypti* versus uninfected controls sampled at two time points (dpi). SRPN: *serpin*.

### Gene ontology (GO) mapping differs across temperature and time

The lists of DEGs identified for each temperature and time point sampled were submitted to the DAVID bioinformatics (V6.8) functional annotation tool. The resulting Gene Ontology (GO) terms were classified using WEGO 2.0 web gene ontology annotation plotting. Gene ontologies in the three major categories of Cellular Location, Molecular Function and Biological Processes differed according to temperature and time of sampling post infection (**Fig. 6**). At 3 dpi, a total of 14 cellular location terms were obtained from the DEGs identified (**Fig. 6A** left panel) with the majority being present at 18 °C. Consistent with the low number of DEGs found at 32 °C, only very few cellular location terms were mapped for this temperature. Only five GO terms pertaining to cellular locations were mapped at all three temperatures. At 7 dpi, a similar number of cellular locations related to DEGs was mapped as for 3 dpi (n locations = 13, **Fig. 6A** right panel). The highest number of cellular locations was observed for mosquitoes held at 28 °C. At 18 °C, almost all (8/9) cellular locations comprised upregulated DEGs. Both 28 °C and 32 °C were similar with respect to DEG expression. Cell junction, supramolecular complex, synapse and synapse part related genes were upregulated only at 28 °C.

**Fig 6.**
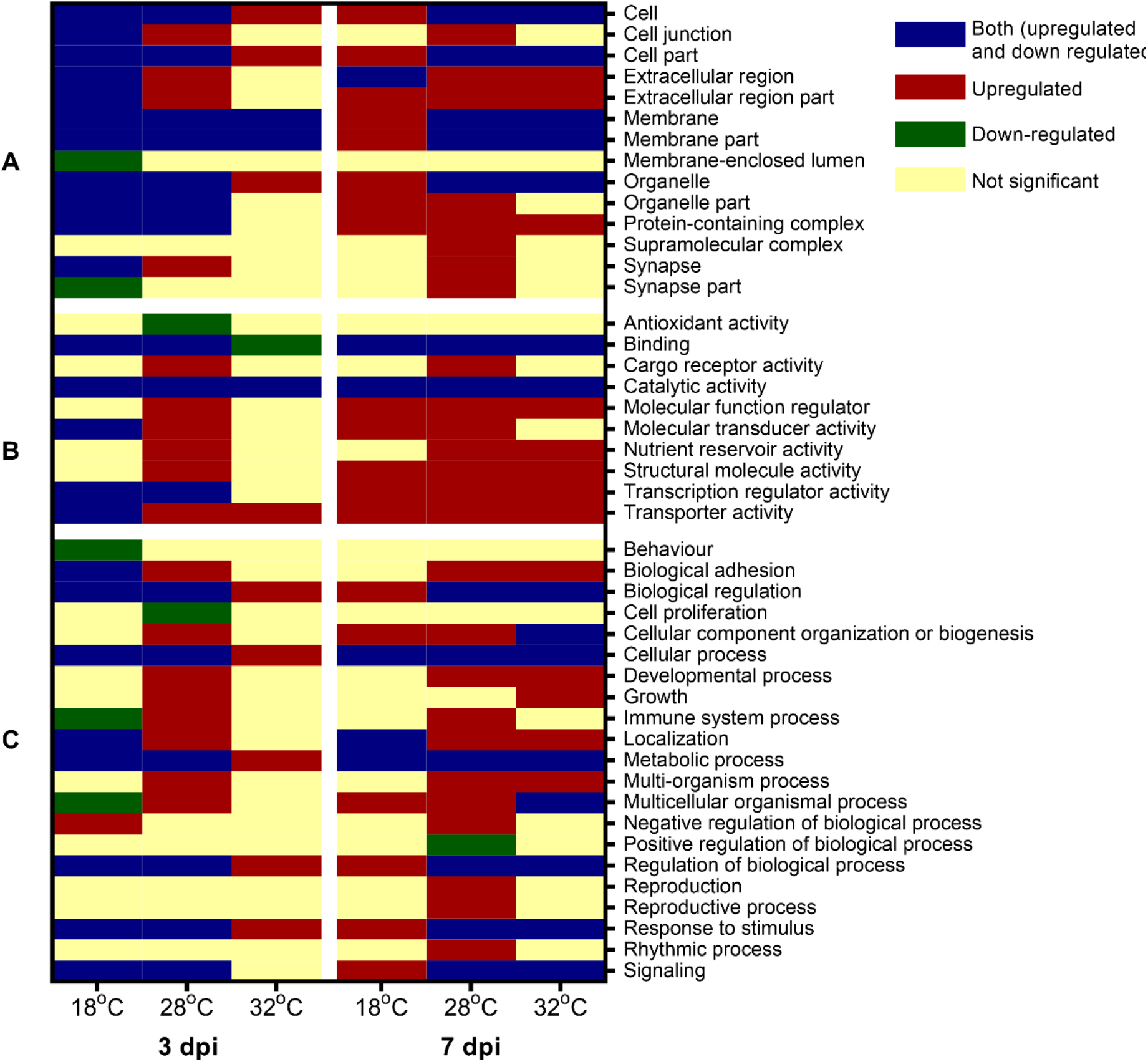
Gene Ontology analysis of differentially expressed genes (DEGs) in CHIKV-infected *Ae. aegypti* mosquitoes, held at three temperatures and sampled at two time points post infection (dpi). **A**) Cellular locations; **B**) Molecular functions; **C**) Biological processes.

At 3 dpi, DEGs were related to a total of 10 molecular functions (**Fig. 6B** left panel), with catalytic activity significantly differentially expressed at all temperatures. Five, 10 and 3 molecular functions were differentially expressed at 18 °C, 28 °C and 32 °C respectively. Five out of six molecular functions identified in mosquitoes held at 18 °C post infection had significant DEGs (both up and downregulated) while the remaining category, behaviour, was downregulated. Binding was downregulated only at 32 °C, but transporter activity was upregulated. Cargo receptor activity, molecular function regulator, nutrient reservoir activity and structural molecular activity were functions observed only at 28 °C. Mosquitoes held at the other two temperatures had no DEGs related to these molecular functions. At 7 dpi, six out of nine molecular functions were found in common across all temperatures (**Fig 6B** right panel). Cargo receptor activity was only significantly upregulated at 28 °C. Molecular transducer activity was observed to be upregulated at 18 °C and 28 °C, whereas nutrient reservoir activity was upregulated at 28 °C and 32 °C.

DEGs in CHIKV-infected mosquitoes versus controls sampled at 3 dpi related 17 biological processes, across the three different temperatures (**Fig 6C** left). Mosquitoes from 28 °C displayed the highest number (15/17) of differentially expressed biological processes, whereas those kept at 32 °C displayed the lowest (5/21). Downregulation of behaviour was unique for mosquitoes held at 18 °C. On the other hand, downregulation of cell proliferation and upregulation of cellular component organisation of biogenesis and development process were exclusively found at 28 °C. Immune system and multicellular organismal processes were downregulated at 18 °C but upregulated at 28 °C. At 7 dpi, 19 biological functions were identified from DEGs across all temperatures (**Fig 6C** right), with the highest number (n=18) identified at 28 °C. Upregulation of immune system process and negative regulation of reproduction were exclusively found at 28 °C. The majority (6/9) of the biological processes found at 18 °C consisted of upregulated DEGs. Further, 62% (8/13) of the biological processes at 32 °C were significantly altered through up- and downregulation.

### Impact of temperature on functional pathway enrichment

KEGG pathways obtained from the DAVID Bioinformatics analysis above were manually categorised into KEGG orthologies (groups of pathways) to determine functional enrichment. The vast majority of pathways altered by temperature, at both time points, were involved in metabolism (**Fig. 7**, top panels denoted by **A**). At 3 dpi, 36 metabolic pathways differed significantly between CHIKV-infected and control mosquitoes across all temperatures. The majority of pathways found at 18 °C were downregulated, suggesting some degree of metabolic shutdown in infected mosquitoes. Downregulated pathways were mostly related to carbohydrate, amino acid and glycan metabolism (**Fig. 7**, top panels denoted by **A**). At 32 °C, all three enriched pathways were downregulated and categorised under metabolic pathways and carbohydrate metabolism. By day 7 post infection, 44 metabolic pathways differed between CHIKV-infected and control mosquitoes. Unlike for 3 dpi, there was no difference in enrichment of metabolic pathways at 18 °C between the two mosquito groups. By contrast, 21 pathways were upregulated at 32 °C at this timepoint compared to only three at 3 dpi. A greater number of enriched metabolic pathways was also observed in mosquitoes at 28 °C at 7 dpi versus 3 dpi. Ascorbate and aldarate metabolism (aag00053), glycerophospholipid metabolism (aag00564) and ether lipid metabolism (aag00565) DEGs were only observed at 32 °C.

**Fig 7.**
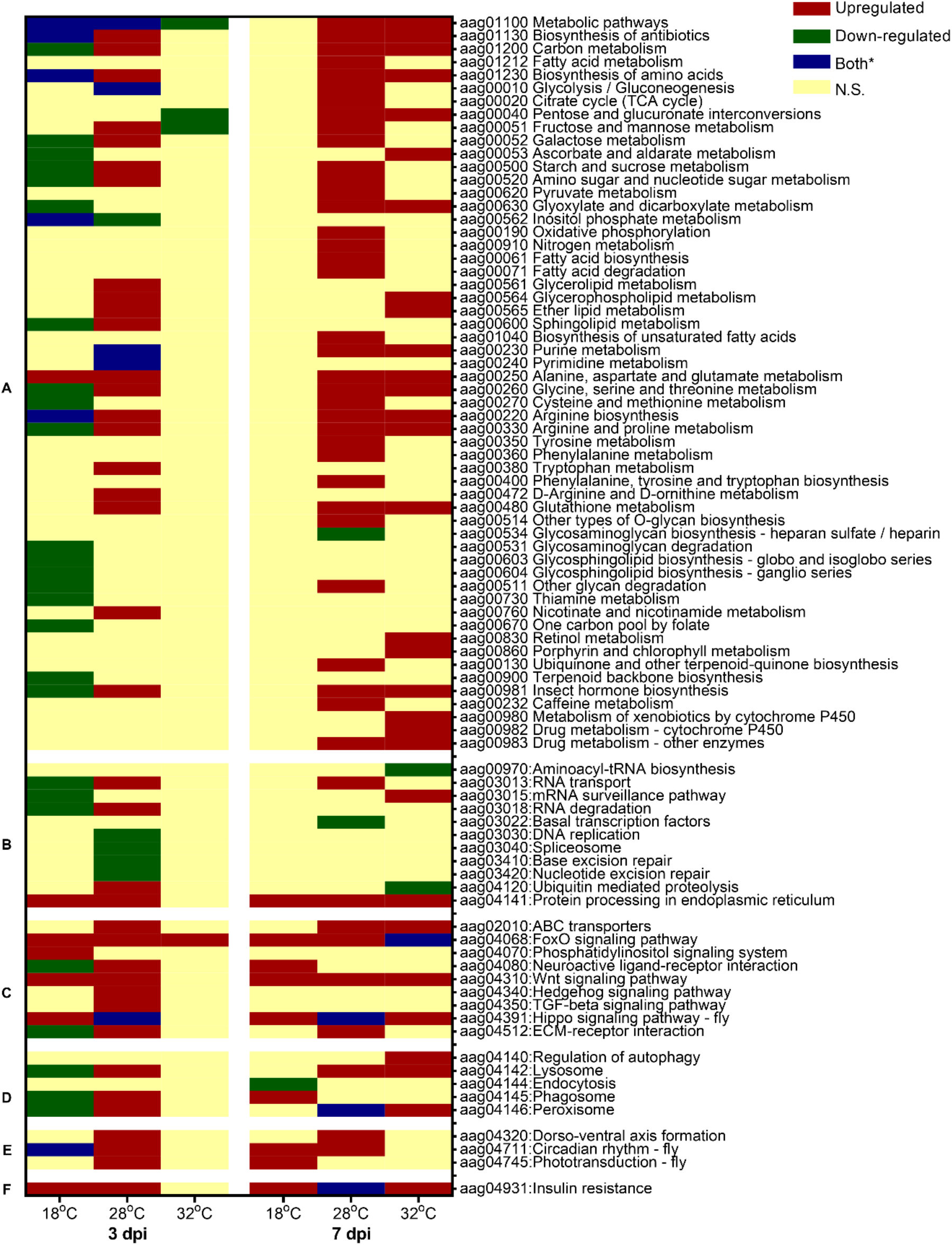
KEGG pathway analysis for DEGs found at 3dpi and 7dpi. **A**) metabolic pathways; **B**) genetic information processing; **C**) environmental information processing; **D**) cellular processes; **E**) organismal systems; **F**) human diseases; ‘aag’ followed by numbers indicate the KEGG pathway identification number. *Denotes both upregulated and downregulated components to the pathway.

Pathways involved in genetic and environmental information processing and cellular processes also showed enrichment in CHIKV-infected mosquitoes compared with uninfected controls. At 3 dpi, involvement of pathways in these categories was observed only at 18 °C and 28 °C, with only FoxO signalling present at 32 °C. Almost half of the 15 non-metabolic pathways enriched at 18 °C were downregulated (**Fig. 7B-F**). Pathways involved in response to RNA virus infection such as lysosome, phagosome, peroxisome, RNA transport and RNA degradation were significantly downregulated at 18 °C but upregulated at 28 °C (**Fig. 7C-D**). Some pathways were exclusively differentially expressed at 28 °C, including DNA replication, spliceosome, base excision repair, nucleotide excision repair, ABC transporters, Hedgehog pathway, TGF-beta signalling pathway, dorso-ventral axis formation and phototransduction-fly (**Fig. 7B-C**). At 7 dpi, the majority of pathways detected were upregulated (9/10, 10/13 and 9/12 at 18 °C, 28 °C and 32 °C, respectively; **Fig. 7**). Upregulation of phagosome and downregulation of endocytosis (aag04144) were exclusively found at 18 °C whilst upregulation of RNA transport and ECM receptor interaction were unique to 28 °C. On the other hand, upregulation of mRNA surveillance pathway and regulation of autophagy (aag04140), but downregulation of aminoacyl tRNA biosynthesis and ubiquitin-mediated proteolysis were distinctly seen at 32 °C. Most mosquito groups, except those held at 32 °C for 3dpi, showed enrichment of insulin resistance pathway.

### Unmapped genes in gene enrichment analysis and long non-coding RNA (lncRNA)

We found 266 upregulated genes and 80 downregulated genes that could not be mapped with DAVID bioinformatics cloud map (**Fig. 8**; **Supplementary Table 7**). Among these, 123 upregulated and 40 downregulated lncRNA genes were found (**Supplementary Table 8**). That is, on average, nearly 50% of unmapped genes were found to comprise lncRNAs. Overall, 10.32% of all upregulated and 16.06% of all downregulated DEGs were lncRNAs.

**Fig 8.**
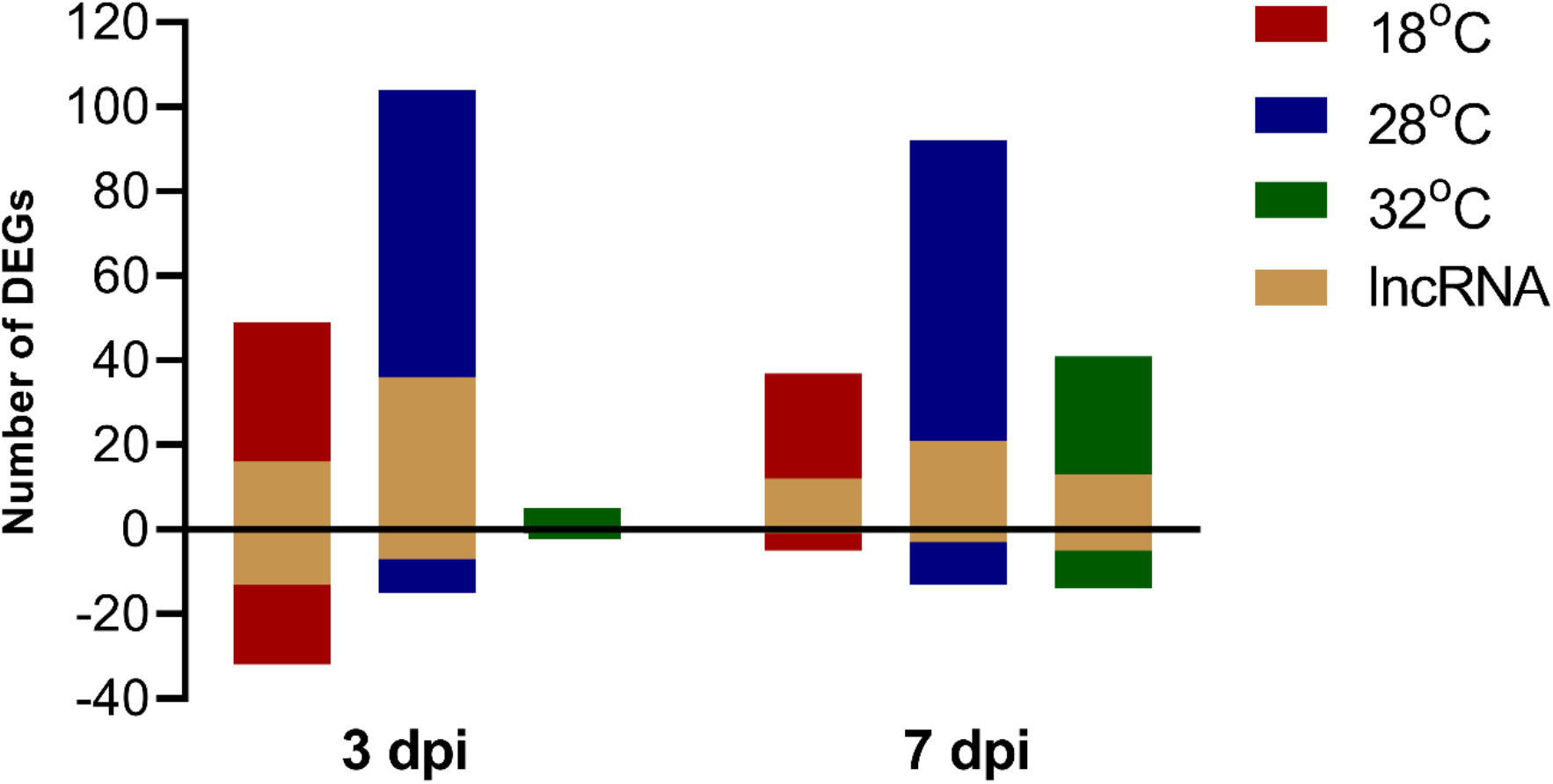
Number of differentially-expressed genes (DEGs) that could not be mapped with DAVID bioinformatics cloud map. The number of lncRNA genes is shown within the total DEGs.

## DISCUSSION

Ambient temperature influences the ability of mosquitoes to transmit viruses but the molecular mechanisms underpinning this are not well understood. Consistent with previous reports, mosquitoes in our experiment that were held at relatively low ambient temperatures showed reduced CHIKV replication. Correspondingly, we observed increased replication at 3 dpi at the highest temperature used here. Distinct mosquito gene expression profiles underpinned infection response at different ambient temperatures, both in the absolute number of DEGs but also their gene identities. Similar to our findings for 28 °C, a recent study of mosquitoes held at 30 °C found a larger number of DEGs expressed at 3 dpi in response to CHIKV [37]. However, exposure to 32 °C in our study elicited a surprisingly low number of genes being expressed in response to CHIKV compared to control uninfected mosquitoes, particularly at the earlier time point. This suppressed response is consistent with increased CHIKV replication at this temperature and suggests that, at high temperatures, mosquitoes are unable to mount an effective defence soon after virus infection.

We observed that the repertoire of immune genes differentially expressed in response to CHIKV infection differed across all temperatures at each time point, contrary to what might have been expected [38, 39]. The first step in eliciting an immune response is the recognition of pathogens by pattern recognition receptors [40]. Similar to previous studies [37–39, 41], we identified diverse PRR genes being differentially expressed in our study, including CLIP-domain serine protease family B, FREP, LRR, leucine-rich transmembrane proteins, CTLs, TEP and Galectins. However, we did not find any PRR DEGs common across all temperature treatments at 3 dpi, while at 7 dpi only *CLIPD6* was shared. There were no immune signalling or pathogen destruction-related genes found at the higher temperature (32 °C), in contrast to the mosquitoes held at 28 °C at both time points, which showed extensive upregulation of genes in both categories. Mosquitoes held at 18 °C and 28 °C showed significant upregulation of *Dicer-2*, critical in antiviral defence in *Aedes* spp. mosquitoes [42], but there was no differential expression of this at 32 °C. We observed additional immune genes become expressed at 32 °C at 7 dpi as compared to 3 dpi, however this involved far fewer genes than at 28 °C and *Dicer-2* was not differentially expressed. A limitation of our study is that we cannot rule out the presence of increased RNAi (*Dicer-2*) response immediately post infection (i.e. within 24 hours), since our earliest time point was 3 dpi. Taken together, overall, our data suggest that the immune response to CHIKV is robust at 28 °C, but key components fail to be activated at higher temperatures.

Complementing the strong immune response observed at 28 °C, comprising Toll, IMD and JAK-STAT pathway components and *Dicer-2* activity, analysis on non-classical components of immunity revealed the doubling of Cytochrome P450 and serine protease gene numbers from 3 to 7 dpi in response to CHIKV infection. Cytochromes are generally involved in cellular functions including oxidative stress, respiration, apoptosis and xenobiotic metabolism [43]. The involvement of Cytochrome P450 in midguts of *Ae. aegypti* in response to DENV has been previously reported [44]. Increases in number of genes coding for serine proteases have also been reported for *Ae. aegypti* during DENV infection [45], thought to activate immune pathways through the triggering of serine protease cascades [46]. However, serine proteases may also aid arbovirus infection through proteolysis of extracellular matrix proteins, facilitating viral attachment [47]. Despite these strong immune defences at 28 °C, we still observed an increase in CHIKV replication over time.

In our study, the highest number of downregulated genes was found at 18 °C at 3 dpi. Experiments on *Drosophila* indicate that exposure to non-optimal/ stressful low or high temperatures causes a significant reduction of lipid storage [48]. At low temperatures, there are reduced energy reserves owing to the slow accumulation of fat triggered by impaired of biochemical activities. The lowering of metabolic rates may lead to a slowdown in the biosynthesis of key host resources necessary to the virus lifecycle, resulting in reduced CHIKV replication. Consistent with this, we also observed downregulation of genes responsible for nucleic acid binding, indicating disturbance to gene regulation [49].

Mosquitoes have evolved various strategies to cope with different thermal conditions such as acclimation, adjusting behavioural activity and synthesis of heat shock proteins [50–52]. The downregulation at 32 °C and 7 dpi of two sensory genes, namely odorant binding protein and long wavelength opsin, suggest a possible effect on mediators of behavioural activity. Insect long wavelength opsins have previously been implicated in insect thermoregulatory responses [53]. Production of HSP proteins such as HSP70 and small HSPs [54] can result in more robust activation of insect defence mechanisms, and insect cells against mechanical and chemical stresses caused by damage to host tissue by invading pathogens [55]. Accordingly, we found significantly upregulated *hsp*70 gene expression in response to CHIKV infection across all temperatures in our study. Acclimation to cold temperatures also leads to elevation in HSP production [56]. Consistent with this, we found that the highest number of *hsp*70 genes expressed at 18 °C, particularly at 3 dpi. It is worth noting that a member (*l2efl*) of the small heat shock protein *hsp*20 family, which has been suggested to suppress virus entry and/ interact with viral proteins [57], was in the top 10 upregulated genes across all temperatures and time points in our study. This is the first time, to our knowledge, that *hsp*20 involvement has been identified in *Ae. aegypti* in response to CHIKV infection and may be specific to this interaction, as it has not been reportedly widely during arbovirus infection of mosquitoes.

We provide an updated list of genes involved in immune response, based on the most recent genome annotation of *Ae. aegypti*. Previous studies of transcriptomic changes may be hampered by limited annotation and minimal literature on how immune genes/gene nomenclatures have changed from the *Ae. aegypti* reference genome [58] assembly version AaegL1 released in 2007 [59] to the AaegL5.2 [60, 61] version published in 2019. A limitation of our experiment is that we cannot rule out the influence of blood meal digestion in DEG patterns observed at 3 dpi. Although digestion may be completed by 3 dpi at the higher temperatures, at 18 °C it may take longer than three days due to a slowdown in metabolism. Consistent with this, we observed significant downregulation of a zinc carboxypeptidase (AAEL008609) involved in blood meal digestion [62].

Overall, our data suggest that temperature strongly influences the ability of mosquitoes to transmit viruses and the immune response during infection. At lower temperatures, downregulation of genes and conservation of resources may drive the viral replication observed despite the activation of many classical and non-classical immune components. In contrast, at high temperatures, the impairment of immune responses may result in shorter virus extrinsic incubation periods and higher virus titers. Impaired mosquito defences may result in a greater propensity for viruses to emerge in a warming climate. Conversely, impairment may also impose additional strong selection pressure on mosquitoes at high temperatures, resulting in altered behaviour and geographic ranges.

A final important observation is that time matters when dissecting response to infection, as varying repertoires of genes may be expressed at different points. Our data challenge the assumption that immune response to infection is constant through time in persistently infected mosquitoes. There was a 10-fold decrease from 3 dpi to 7 dpi in the number of downregulated genes in the number of DEGs observed at 18 °C. Conversely, a 6.5-fold increase in the number of upregulated DEGs was observed at 32 °C. The number of DEGs at 28 °C remained more or less constant across time points but different repertoires of genes were expressed. In parallel, gene ontologies identified, and pathways enriched at these temperatures also differed.

In conclusion, we show that ambient temperature influences overall gene expression in response to CHIKV infection and suggest that high temperatures may impair mosquito immune response. Impaired mosquito responses may accelerate transmission of arboviruses and, potentially, pathogen emergence. The presence of a considerable number of lncRNA, pseudogenes and uncharacterised genes significantly differentially expressed in infected mosquitoes highlights the need for further functional studies and annotation of *Ae. aegypti* genome.

## METHODS

### Mosquitoes and virus infection

Five to 7-day old *Ae. aegypti* were orally challenged with virus-infected or sheep blood alone (control), using methods previously described [26]. A CHIKV strain from the Asian Genotype (GenBank ID: MF773560) was prepared as described in [26] and delivered in oral feeds at a final pfu of 1 × 10^7^ per ml of virus stock. Post oral challenge, mosquitoes observed to have taken a blood meal were randomly allocated to three different temperatures (18 °C, 28 °C and 32 °C) in environmental chambers, with 70% humidity at a 12h:12h day/ night cycle. Mosquitoes were then sampled at 3 and 7 post-infection. Mosquitoes were tested for the presence of CHIKV in whole bodies (minus wings and legs) using qRT-PCR as previously described [26].

### RNA extraction and NGS sequencing

RNA was extracted from individual mosquito bodies using TRIzol™ reagent (Invitrogen™, Thermo Fisher Scientific, USA). Samples were homogenised for 90 seconds with the addition of silica glass beads (Daintree Scientific, St Helens, TAS, Australia) in a MiniBeadbeater-96 (Biospec Products, Bartlesville, Oklahoma, USA). Total RNA was extracted from the homogenate according to the Trizol protocol. RNA was dissolved in Ultrapure™ water (Invitrogen™, Thermo Fisher Scientific, USA) to make a final volume of 40 μL, and frozen at −80 °C until further analysis.

### Sample selection and RNAseq

Six mosquito bodies in which we detected the presence of CHIKV were selected from each time point, for each temperature. The samples were checked for RNA quality >1.8 of A260/280 ratio and quantity by NanoDrop Lite (Thermo Fisher Scientific Inc.). RNA integrity was checked using RIN score analysis performed in an Agilent 2100 Bioanalyzer (Agilent Technologies, Palo Alto, CA, USA). Similarly, RNA extracted from uninfected control mosquitoes was subjected to quality and quantity checking. A total of 72 RNA samples, comprising 36 infected blood-fed and 36 uninfected blood-fed controls were subjected to RNAseq, with six per each combination of time point (3 dpi and 7 dpi), and temperature (18 °C, 28 °C and 32 °C). Samples were sequenced on the Illumina HiSeq platform at Genewiz, China. Illumina raw data generated for all samples were deposited at the Short Read Archive (SRA) database under the BioProject accession number PRJNA630779.

### RNAseq data analysis

Sequencing data obtained from Genewiz, China were subjected to quality control and mapping. Raw sequences in fastq format were subjected to adapter removal and quality trimming using Trim Galore (https://github.com/FelixKrueger/TrimGalore). Poor quality bases/reads with a quality score lower than 30 and sequences with a read length shorter than 50 nucleotides were removed to obtain high-quality clean data. To map the sequences, we first downloaded the reference genome sequences and annotations of AaegL5.2 from VectorBase (AaegL5) [38], the most recent annotation release of *Ae. aegypti* [51] genome. Second, RNA STAR two-pass mapping approach was used to align clean reads onto the AaegL5 reference genome sequence and obtain gene read counts [41]. The RSEM tool was then used to quantify the expression of gene isoforms [63].

### DEG identification, GO mapping and KEGG pathway analysis

Differential expression analysis was performed using the DESeq2 Bioconductor package [64]. The comparison between the mapped read counts of the virus-infected mosquitoes vs the read counts of uninfected blood-fed mosquitoes identified differentially expressed genes (DEGs). Genes were significantly differentially expressed if the adjusted p-value (false discovery rate) was <0.05 and showed an absolute fold change of ±1.5. For all DEGs, gene annotation was done using the Biomart tool provided by the VectorBase database. DEG lists were further used for GO, pathway enrichment and immune response analysis. DAVID bioinformatics (V6.8) was used to assign GO terms and KEGG pathways information to differentially expressed genes [65]. WEGO 2.0 was then used to compare and plot GO annotation results at user-specified hierarchical level 2 [65]. The KEGG pathways enriched by DAVID bioinformatics were manually categorized under six main categories and subcategories as defined by the KEGG pathway database [36]. There were some pathways/GO terms with the presence of both up and downregulated genes. This is expected when pathways encode both positive and negative regulators [66]. Thus, if the upregulated gene group contains “positive” regulators and downregulated one includes “negative” regulators, in general, the pathway can be considered as regulated [67]. We used this approach in our descriptions of GO analysis and KEGG pathway analysis. For gene IDs for which functional information could not be assigned via DAVID bioinformatics (v6.8) gene enrichment analysis tool, then such genes were subjected to annotation using VectorBase search and BLAST sequence similarity (https://blast.ncbi.nlm.nih.gov/Blast.cgi).

### Identification of classical and non-classical immune genes

We downloaded the peptide sequences of all the immune genes listed at the database ImmunoDB (http://cegg.unige.ch/Insecta/immunodb), which possesses information on insect immune-related genes and gene families [39]. We performed a similarity search using blastp with the peptides given in VectorBase (https://www.vectorbase.org/) under the annotation release AaegL5. Using an in-house script, we obtained the 10 best hits for each peptide and then selected the top hit with the largest bit score, % identity and lowest E-value. Next, we further searched the literature for articles using the search terms “*Aedes aegypti*” and “immune” published after 2007 to ensure that we identify novel gene families/ genes, that is, the genes recently discovered to link with immune response and thereby not covered in 2007 annotation of *Ae. aegypti*. In addition, a manual search for immune-related gene families/ genes identified VectorBase annotated genes using previously identified gene family names. Genes that were related to four main categories of immune response process were considered classical immune genes while the genes directly or indirectly supplementing the above four processes are regarded as non-classical immune genes. The list of DEGs were compared against the list of immune genes identified through both literature review and similarity screening using Venny 2.1. Additionally, a list of lncRNA was identified comparing conserved ID sets between AaegL3.1 and AaegL5.1 gene annotation, and by including lncRNAs reported in AaegL5.2 [63].

### CHIKV read count analysis

Estimation of chikungunya virus replication in infected and control *Ae. aegypti* was also performed by mapping high-quality adaptor-clipped Illumina pair-end reads onto the chikungunya virus genome, Asian genotype (GenBank accession no. MF773560) using Burrows-Wheeler Aligner (BWA) mem (https://arxiv.org/abs/1303.3997). Read counts of aligned reads were obtained using samtools idxstats [68]. Normalised Reads Per Million (RPM) counts were calculated to quantify viral read counts. Two samples (RY-74 and RY-75) were identified as outliers on the correlation plot between virus titer and normalized reads per million counts and were removed from further analysis. The Mann-Whitney test was used to find differences between virus read counts of any two mosquito groups. The Kruskal-Wallis test was used for comparison of viral reads of multiple groups. A p-value <0.05 was considered statistically significant. Statistical analyses were performed using SPSS 25 (IBM Statistics). Graphs were prepared using GraphPad Prism version 8.3.0.

## ACKNOWLEDGEMENTS

We thank Mr Henry Simila for technical assistance. The research presented here was funded by the National Health and Medical Research Council (NHMRC) of Australia project grant APP1125317 and the Royal Society of Tropical Medicine and Hygiene small grants program. BMC Randika Wimalasiri-Yapa was funded by the University Grants Commission Sri Lanka, The Open University of Sri Lanka and Queensland University of Technology, Brisbane, Australia.

## Notes

### Competing Interest Statement

The authors have declared no competing interest.

