## Supplementary Document for "Temperature alters gene expression in mosquitoes during arbovirus infection"

Supplementary Table 1. Mapping statistics.

| Sample ID | Raw reads R1 | Raw reads R2 | Mapped read pairs | % of mapped reads |
|---|---|---|---|---|
| **RY-01** | 40,285,352 | 40,285,352 | 37,623,151 | 93.39 |
| **RY-02** | 38,908,531 | 38,908,531 | 36,441,243 | 93.66 |
| **RY-03** | 42,009,688 | 42,009,688 | 39296180 | 93.54 |
| **RY-04** | 42,984,040 | 42,984,040 | 40141389 | 93.39 |
| **RY-05** | 46,411,651 | 46,411,651 | 43128300 | 92.93 |
| **RY-06** | 37,002,005 | 37,002,005 | 34379255 | 92.91 |
| **RY-07** | 34,130,936 | 34,130,936 | 31,994,356 | 93.74 |
| **RY-08** | 32,190,705 | 32,190,705 | 30047114 | 93.34 |
| **RY-09** | 37,578,836 | 37,578,836 | 35252656 | 93.81 |
| **RY-10** | 38,453,304 | 38,453,304 | 36101459 | 93.88 |
| **RY-12** | 39,543,441 | 39,543,441 | 37166365 | 93.99 |
| **RY-13** | 41,962,994 | 41,962,994 | 39394169 | 93.88 |
| **RY-14** | 33,822,121 | 33,822,121 | 30787910 | 91.03 |
| **RY-15** | 32,192,708 | 32,192,708 | 29,206,064 | 90.72 |
| **RY-16** | 37,765,250 | 37,765,250 | 34188323 | 90.53 |
| **RY-17** | 38,591,247 | 38,591,247 | 35751876 | 92.64 |
| **RY-18** | 40,984,117 | 40,984,117 | 38603474 | 94.19 |
| **RY-19** | 37,091,930 | 37,091,930 | 34923378 | 94.15 |
| **RY-20** | 37,073,879 | 37,073,879 | 34809347 | 93.89 |
| **RY-21** | 38,291,431 | 38,291,431 | 35914033 | 93.79 |
| **RY-22** | 43,858,575 | 43,858,575 | 41268848 | 94.1 |
| **RY-23** | 30,765,242 | 30,765,242 | 28,836,103 | 93.73 |
| **RY-24** | 30,693,689 | 30,693,689 | 28,845,836 | 93.98 |
| **RY-25** | 38,692,441 | 38,692,441 | 36,421,995 | 94.13 |
| **RY-26** | 35,516,428 | 35,516,428 | 33,342,457 | 93.88 |
| **RY-28** | 38,218,083 | 38,218,083 | 35,909,209 | 93.96 |
| **RY-29** | 30,004,369 | 30,004,369 | 28200236 | 93.99 |
| **RY-30** | 38,562,521 | 38,562,521 | 36,298,967 | 94.13 |
| **RY-31** | 35,832,268 | 35,832,268 | 33,766,480 | 94.23 |
| **RY-32** | 35,417,713 | 35,417,713 | 33,403,827 | 94.31 |

| | | | | |
|---|---|---|---|---|
| **RY-33** | 45,325,872 | 45,325,872 | 42,911,373 | 94.67 |
| **RY-34** | 42,575,000 | 42,575,000 | 40,117,155 | 94.23 |
| **RY-36** | 30,835,998 | 30,835,998 | 29,191,850 | 94.67 |
| **RY-37** | 34,601,897 | 34,601,897 | 32,717,138 | 94.55 |
| **RY-38** | 37,350,308 | 37,350,308 | 35,344,436 | 94.63 |
| **RY-39** | 39,612,660 | 39,612,660 | 37,360,181 | 94.31 |
| **RY-40** | 41,740,294 | 41,740,294 | 39,480,900 | 94.59 |
| **RY-41** | 36,497,253 | 36,497,253 | 34,491,830 | 94.51 |
| **RY-42** | 40,526,256 | 40,526,256 | 38,210,454 | 94.29 |
| **RY-43** | 40,632,982 | 40,632,982 | 38,492,323 | 94.73 |
| **RY-44** | 34,957,475 | 34,957,475 | 32,910,879 | 94.15 |
| **RY-45** | 35,231,754 | 35,231,754 | 33,230,913 | 94.32 |
| **RY-46** | 33,628,624 | 33,628,624 | 31,731,002 | 94.36 |
| **RY-47** | 46,187,306 | 46,187,306 | 38,243,073 | 82.8 |
| **RY-48** | 39,147,303 | 39,147,303 | 36,894,388 | 94.25 |
| **RY-49** | 32,333,451 | 32,333,451 | 30,061,200 | 92.97 |
| **RY-50** | 31,383,630 | 31,383,630 | 29,628,090 | 94.41 |
| **RY-51** | 34,759,953 | 34,759,953 | 32,787,103 | 94.32 |
| **RY-52** | 37,588,740 | 37,588,740 | 35352578 | 94.05 |
| **RY-53** | 40,571,112 | 40,571,112 | 38250283 | 94.28 |
| **RY-54** | 57,388,878 | 57,388,878 | 54,162,441 | 94.38 |
| **RY-56** | 40,163,339 | 40,163,339 | 37755106 | 94 |
| **RY-57** | 38,051,883 | 38,051,883 | 35830088 | 94.16 |
| **RY-58** | 36,193,477 | 36,193,477 | 34102310 | 94.22 |
| **RY-59** | 35,544,173 | 35,544,173 | 32,389,236 | 91.12 |
| **RY-60** | 44,312,210 | 44,312,210 | 40314320 | 90.98 |
| **RY-61** | 36,649,859 | 36,649,859 | 33,445,721 | 91.26 |
| **RY-63** | 36,481,409 | 36,481,409 | 33273723 | 91.21 |
| **RY-64** | 36,567,180 | 36,567,180 | 33,403,693 | 91.35 |
| **RY-65** | 36,197,587 | 36,197,587 | 33,021,251 | 91.23 |
| **RY-66** | 37,734,511 | 37,734,511 | 34,319,093 | 90.95 |
| **RY-67** | 35,750,054 | 35,750,054 | 32,682,290 | 91.42 |
| **RY-68** | 32,756,364 | 32,756,364 | 29,426,665 | 89.83 |
| **RY-69** | 36,849,343 | 36,849,343 | 33,560,238 | 91.07 |

| | | | | |
|---|---|---|---|---|
| **RY-70** | 40,083,413 | 40,083,413 | 36,610,308 | 91.34 |
| **RY-71** | 34,061,633 | 34,061,633 | 31,078,074 | 91.24 |
| **RY-72** | 38,569,119 | 38,569,119 | 35,299,390 | 91.52 |
| **RY-73** | 32,810,943 | 32,810,943 | 30,310,061 | 92.38 |
| **RY-74** | 30,112,259 | 30,112,259 | 28,008,286 | 93.01 |
| **RY-75** | 29,952,637 | 29,952,637 | 28,126,155 | 93.9 |
| **RY-76** | 33,478,029 | 33,478,029 | 31,259,937 | 93.37 |
| **RY-77** | 36,205,058 | 36,205,058 | 33,836,239 | 93.46 |
| **Total** | 2,696,236,721 | 2,696,236,721 | 2,511,065,774 | 93.13 |

Supplementary Table 2. All DEG lists.

**3 dpi 18 $^oC$ upregulated**

| GeneID | Base mean | log2(FC) | StdErr | Wald-Stats | P-value | P-adj |
|---|---|---|---|---|---|---|
| AAEL020330 | 3734.941 | 5.535313 | 0.380164 | 14.56031 | 5.02E-48 | 6.17E-44 |
| AAEL017976 | 12407.69 | 5.247997 | 0.368873 | 14.22711 | 6.22E-46 | 3.82E-42 |
| AAEL013346 | 1482.01 | 5.139391 | 0.383931 | 13.38625 | 7.28E-41 | 2.98E-37 |
| AAEL013345 | 2014.426 | 4.956533 | 0.402408 | 12.31717 | 7.32E-35 | 2.25E-31 |
| AAEL013350 | 6345.489 | 4.609722 | 0.398557 | 11.56602 | 6.13E-31 | 1.50E-27 |
| AAEL013348 | 654.2395 | 4.427207 | 0.392608 | 11.27639 | 1.72E-29 | 3.51E-26 |
| AAEL013351 | 1373.552 | 3.823111 | 0.343999 | 11.11371 | 1.08E-28 | 1.89E-25 |
| AAEL013339 | 296.5337 | 4.26091 | 0.415188 | 10.26262 | 1.04E-24 | 1.59E-21 |
| AAEL013349 | 1012.989 | 3.979345 | 0.396856 | 10.02717 | 1.16E-23 | 1.58E-20 |
| AAEL023321 | 205.6866 | 3.799456 | 0.381041 | 9.971254 | 2.04E-23 | 2.50E-20 |
| AAEL017975 | 21388.26 | 3.856276 | 0.406555 | 9.485246 | 2.42E-21 | 2.70E-18 |
| AAEL026300 | 452.9175 | 3.817729 | 0.42868 | 8.905779 | 5.30E-19 | 5.42E-16 |
| AAEL022253 | 11494.01 | 3.49052 | 0.422974 | 8.252319 | 1.55E-16 | 1.47E-13 |
| AAEL003505 | 2500.659 | 2.027496 | 0.25007 | 8.107707 | 5.16E-16 | 4.52E-13 |
| AAEL014531 | 1387.135 | 2.153974 | 0.279854 | 7.696777 | 1.40E-14 | 1.14E-11 |
| AAEL014843 | 39193.04 | 1.301246 | 0.17041 | 7.635994 | 2.24E-14 | 1.72E-11 |
| AAEL022079 | 771.3579 | 2.768739 | 0.364815 | 7.589426 | 3.21E-14 | 2.32E-11 |
| AAEL013344 | 3116.273 | 2.221841 | 0.299912 | 7.40831 | 1.28E-13 | 7.95E-11 |
| AAEL017380 | 1445.159 | 3.169652 | 0.427951 | 7.406578 | 1.30E-13 | 7.95E-11 |
| AAEL001800 | 742.0151 | 1.166323 | 0.161765 | 7.20996 | 5.60E-13 | 3.27E-10 |
| AAEL026751 | 8475.504 | 1.810765 | 0.268794 | 6.736626 | 1.62E-11 | 9.04E-09 |
| AAEL004090 | 1268.531 | 1.454383 | 0.227733 | 6.386355 | 1.70E-10 | 8.69E-08 |
| AAEL019935 | 7915.363 | 0.665429 | 0.106208 | 6.265334 | 3.72E-10 | 1.83E-07 |
| AAEL027610 | 4012.713 | 2.669849 | 0.430476 | 6.202079 | 5.57E-10 | 2.63E-07 |
| AAEL024512 | 229.4591 | 2.603068 | 0.426477 | 6.103648 | 1.04E-09 | 4.55E-07 |
| AAEL001682 | 232.9808 | 1.234114 | 0.204104 | 6.046488 | 1.48E-09 | 6.27E-07 |
| AAEL010680 | 158.535 | 2.096026 | 0.356294 | 5.882859 | 4.03E-09 | 1.60E-06 |
| AAEL006883 | 2732.59 | 2.196196 | 0.377811 | 5.812946 | 6.14E-09 | 2.31E-06 |
| AAEL001857 | 489.1288 | 2.338409 | 0.409105 | 5.71591 | 1.09E-08 | 3.94E-06 |

| | | | | | | |
|---|---|---|---|---|---|---|
| AAEL012712 | 384.0227 | 2.204887 | 0.391418 | 5.633081 | 1.77E-08 | 5.81E-06 |
| AAEL002969 | 1497.899 | 1.8545 | 0.32938 | 5.630278 | 1.80E-08 | 5.81E-06 |
| AAEL026833 | 1068.714 | 1.7364 | 0.319538 | 5.434093 | 5.51E-08 | 1.65E-05 |
| AAEL000850 | 353.5365 | 1.258177 | 0.233109 | 5.397374 | 6.76E-08 | 1.98E-05 |
| AAEL021614 | 20.26452 | 2.173809 | 0.406643 | 5.345736 | 9.01E-08 | 2.57E-05 |
| AAEL008953 | 13844.17 | 1.054405 | 0.198955 | 5.29972 | 1.16E-07 | 3.24E-05 |
| AAEL015609 | 36.04469 | 1.423921 | 0.270591 | 5.262256 | 1.42E-07 | 3.88E-05 |
| AAEL013314 | 912.1895 | 0.946075 | 0.186049 | 5.085087 | 3.67E-07 | 9.60E-05 |
| AAEL023591 | 2862.812 | 1.315018 | 0.258998 | 5.077333 | 3.83E-07 | 9.79E-05 |
| AAEL011708 | 28081.34 | 1.088194 | 0.216052 | 5.036723 | 4.74E-07 | 0.000119 |
| AAEL011371 | 3830.119 | 1.05284 | 0.211939 | 4.967662 | 6.78E-07 | 0.000166 |
| AAEL008007 | 320.1926 | 1.764049 | 0.356803 | 4.944044 | 7.65E-07 | 0.000184 |
| AAEL005558 | 5998.823 | 0.695489 | 0.143783 | 4.837075 | 1.32E-06 | 0.0003 |
| AAEL005992 | 93.65065 | 1.464637 | 0.304043 | 4.817201 | 1.46E-06 | 0.000325 |
| AAEL022059 | 6900.654 | 2.071132 | 0.432829 | 4.785107 | 1.71E-06 | 0.000368 |
| AAEL010411 | 4533.409 | 0.725523 | 0.154427 | 4.698174 | 2.62E-06 | 0.000537 |
| AAEL004589 | 11091.43 | 0.688054 | 0.146272 | 4.703938 | 2.55E-06 | 0.000537 |
| AAEL008622 | 84.70919 | 1.799117 | 0.382886 | 4.698839 | 2.62E-06 | 0.000537 |
| AAEL025126 | 1084.982 | 2.039839 | 0.434553 | 4.694109 | 2.68E-06 | 0.000539 |
| AAEL008473 | 3439.312 | 1.82938 | 0.390621 | 4.683258 | 2.82E-06 | 0.000559 |
| AAEL025531 | 346.7837 | 2.01641 | 0.431726 | 4.670576 | 3.00E-06 | 0.000585 |
| AAEL006794 | 4156.364 | 0.991009 | 0.216748 | 4.57217 | 4.83E-06 | 0.000898 |
| AAEL021302 | 477.8883 | 1.312448 | 0.29467 | 4.453957 | 8.43E-06 | 0.001418 |
| AAEL026519 | 176.072 | 1.193048 | 0.269628 | 4.424789 | 9.65E-06 | 0.001601 |
| AAEL013352 | 325.8598 | 1.829224 | 0.415108 | 4.406623 | 1.05E-05 | 0.001674 |
| AAEL023746 | 464.3846 | 0.921624 | 0.213034 | 4.326193 | 1.52E-05 | 0.002299 |
| AAEL003632 | 57.25144 | 1.830763 | 0.429099 | 4.266529 | 1.99E-05 | 0.002769 |
| AAEL019902 | 1538.573 | 1.347703 | 0.317675 | 4.242402 | 2.21E-05 | 0.00305 |
| AAEL026008 | 3566.907 | 1.412411 | 0.334227 | 4.225901 | 2.38E-05 | 0.003182 |
| AAEL026215 | 2918.491 | 0.870748 | 0.206926 | 4.208014 | 2.58E-05 | 0.0034 |
| AAEL003619 | 1726.592 | 1.540189 | 0.372122 | 4.138936 | 3.49E-05 | 0.004326 |
| AAEL004148 | 5155.337 | 0.586117 | 0.141692 | 4.136556 | 3.53E-05 | 0.004328 |
| AAEL012255 | 163.6476 | 1.475474 | 0.361034 | 4.086798 | 4.37E-05 | 0.005162 |
| AAEL014316 | 72.6363 | 0.858819 | 0.212107 | 4.048996 | 5.14E-05 | 0.005957 |

| | | | | | | |
|---|---|---|---|---|---|---|
| AAEL009055 | 738.5938 | 1.152196 | 0.288026 | 4.000323 | 6.33E-05 | 0.00719 |
| AAEL005032 | 2305.857 | 1.064697 | 0.267315 | 3.982926 | 6.81E-05 | 0.007666 |
| AAEL005656 | 472.1923 | 1.496157 | 0.376347 | 3.97547 | 7.02E-05 | 0.007754 |
| AAEL014363 | 17.91429 | 1.680746 | 0.42306 | 3.972832 | 7.10E-05 | 0.007754 |
| AAEL000812 | 490.3249 | 0.68961 | 0.173908 | 3.96538 | 7.33E-05 | 0.007754 |
| AAEL006069 | 411.6914 | 0.794996 | 0.201135 | 3.952552 | 7.73E-05 | 0.007976 |
| AAEL001565 | 231.7231 | 0.612403 | 0.155297 | 3.943432 | 8.03E-05 | 0.008216 |
| AAEL008101 | 638.0649 | 0.739177 | 0.188032 | 3.931132 | 8.45E-05 | 0.008496 |
| AAEL000338 | 1346.458 | 1.398064 | 0.357216 | 3.913773 | 9.09E-05 | 0.008852 |
| AAEL008607 | 3573.331 | 1.274558 | 0.326845 | 3.89958 | 9.64E-05 | 0.009172 |
| AAEL015631 | 1736.381 | 1.266749 | 0.32516 | 3.895768 | 9.79E-05 | 0.009172 |
| AAEL008050 | 548.0128 | 1.676839 | 0.431892 | 3.88254 | 0.000103 | 0.009262 |
| AAEL013972 | 12.31966 | 1.600317 | 0.412581 | 3.878792 | 0.000105 | 0.009338 |
| AAEL020957 | 1177.887 | 0.666941 | 0.172305 | 3.870702 | 0.000109 | 0.009503 |
| AAEL002655 | 403.1663 | 1.658564 | 0.429288 | 3.86352 | 0.000112 | 0.009527 |
| AAEL012431 | 601.0768 | 1.046079 | 0.270886 | 3.861703 | 0.000113 | 0.009532 |
| AAEL020777 | 7.414424 | 1.639095 | 0.426048 | 3.847211 | 0.000119 | 0.009976 |
| AAEL001675 | 38.09614 | 1.598547 | 0.416466 | 3.838359 | 0.000124 | 0.01018 |
| AAEL022900 | 41.52985 | 1.583063 | 0.412721 | 3.835668 | 0.000125 | 0.01018 |
| AAEL000709 | 10552.29 | 0.755919 | 0.197674 | 3.824069 | 0.000131 | 0.010601 |
| AAEL022363 | 131.5313 | 0.944937 | 0.255111 | 3.704022 | 0.000212 | 0.016018 |
| AAEL006533 | 14.95517 | 1.510581 | 0.407395 | 3.707903 | 0.000209 | 0.016018 |
| AAEL003888 | 2821.622 | 1.213512 | 0.327854 | 3.701374 | 0.000214 | 0.016018 |
| AAEL001367 | 623.7485 | 0.801681 | 0.217583 | 3.68448 | 0.000229 | 0.016744 |
| AAEL008227 | 791.4774 | 0.802876 | 0.218254 | 3.678637 | 0.000234 | 0.017031 |
| AAEL007826 | 3991.395 | 0.604745 | 0.165168 | 3.661386 | 0.000251 | 0.017903 |
| AAEL025532 | 78.26468 | 1.325002 | 0.362452 | 3.65566 | 0.000257 | 0.018201 |
| AAEL000251 | 63.28877 | 1.063487 | 0.293342 | 3.625423 | 0.000288 | 0.019765 |
| AAEL010375 | 13.0691 | 1.571183 | 0.435826 | 3.605069 | 0.000312 | 0.020755 |
| AAEL006449 | 2740.912 | 0.621239 | 0.172353 | 3.604453 | 0.000313 | 0.020755 |
| AAEL004935 | 8.52101 | 1.511901 | 0.420177 | 3.598248 | 0.00032 | 0.020918 |
| AAEL023999 | 15.31619 | 1.463008 | 0.406932 | 3.595211 | 0.000324 | 0.021051 |
| AAEL023745 | 526.295 | 1.096241 | 0.305137 | 3.592623 | 0.000327 | 0.02109 |
| AAEL007233 | 195.8894 | 1.517417 | 0.422445 | 3.591989 | 0.000328 | 0.02109 |

| | | | | | | |
|---|---|---|---|---|---|---|
| AAEL003345 | 14584 | 1.168739 | 0.327088 | 3.573165 | 0.000353 | 0.022525 |
| AAEL025894 | 37.77207 | 1.508803 | 0.427931 | 3.525805 | 0.000422 | 0.025783 |
| AAEL002959 | 465.9473 | 1.474204 | 0.419327 | 3.515641 | 0.000439 | 0.026658 |
| AAEL010068 | 2102.199 | 1.052314 | 0.299589 | 3.512524 | 0.000444 | 0.02684 |
| AAEL006197 | 99.02639 | 0.795746 | 0.226668 | 3.510627 | 0.000447 | 0.026874 |
| AAEL024540 | 312.1506 | 0.931306 | 0.267558 | 3.480759 | 0.0005 | 0.029016 |
| AAEL018304 | 124.587 | 1.104567 | 0.317662 | 3.477173 | 0.000507 | 0.029066 |
| AAEL018301 | 658.8098 | 0.654417 | 0.188382 | 3.473882 | 0.000513 | 0.029288 |
| AAEL007619 | 417.9272 | 0.977578 | 0.281919 | 3.467591 | 0.000525 | 0.029681 |
| AAEL000750 | 2885.777 | 0.742536 | 0.214199 | 3.466577 | 0.000527 | 0.029681 |
| AAEL013347 | 1493.641 | 1.061791 | 0.306269 | 3.466861 | 0.000527 | 0.029681 |
| AAEL005890 | 1374.858 | 0.608056 | 0.175604 | 3.462649 | 0.000535 | 0.029961 |
| AAEL020236 | 3572.704 | 0.670662 | 0.194072 | 3.455743 | 0.000549 | 0.030344 |
| AAEL024560 | 88.94283 | 1.043001 | 0.302643 | 3.446309 | 0.000568 | 0.030874 |
| AAEL026868 | 21.72797 | 1.494461 | 0.434361 | 3.440594 | 0.00058 | 0.031249 |
| AAEL010166 | 29.17861 | 1.482285 | 0.431462 | 3.435495 | 0.000591 | 0.031567 |
| AAEL015557 | 212.8244 | 0.90356 | 0.264156 | 3.420554 | 0.000625 | 0.032783 |
| AAEL008213 | 90.29106 | 0.89186 | 0.26122 | 3.414204 | 0.00064 | 0.033012 |
| AAEL010656 | 1644.107 | 1.161836 | 0.340652 | 3.410627 | 0.000648 | 0.033012 |
| AAEL020575 | 74.75739 | 1.384464 | 0.407632 | 3.396357 | 0.000683 | 0.034081 |
| AAEL019995 | 1158.313 | 0.785758 | 0.231543 | 3.393564 | 0.00069 | 0.034285 |
| AAEL026878 | 1423.348 | 1.41347 | 0.417999 | 3.381516 | 0.000721 | 0.034837 |
| AAEL003003 | 122.3534 | 0.73441 | 0.217367 | 3.378668 | 0.000728 | 0.035062 |
| AAEL014335 | 634.6688 | 0.78638 | 0.233875 | 3.362395 | 0.000773 | 0.036383 |
| AAEL021595 | 155.5841 | 0.583431 | 0.173379 | 3.365067 | 0.000765 | 0.036383 |
| AAEL018241 | 157.9798 | 1.265813 | 0.376281 | 3.364006 | 0.000768 | 0.036383 |
| AAEL011412 | 460.3259 | 0.711086 | 0.211569 | 3.361015 | 0.000777 | 0.036383 |
| AAEL012853 | 230.0456 | 1.288257 | 0.383079 | 3.362902 | 0.000771 | 0.036383 |
| AAEL019637 | 1155.443 | 1.089482 | 0.328978 | 3.311713 | 0.000927 | 0.04139 |
| AAEL028247 | 71.85983 | 1.245511 | 0.377009 | 3.30366 | 0.000954 | 0.04229 |
| AAEL011836 | 411.2745 | 0.76168 | 0.230797 | 3.300221 | 0.000966 | 0.042657 |
| AAEL018189 | 31.04136 | 1.295138 | 0.395846 | 3.271821 | 0.001069 | 0.045075 |
| AAEL004124 | 575.2964 | 0.742328 | 0.228034 | 3.255346 | 0.001133 | 0.046495 |
| AAEL013974 | 532.2355 | 0.87561 | 0.269554 | 3.248369 | 0.001161 | 0.047334 |

| | | | | | | |
|---|---|---|---|---|---|---|
| AAEL026194 | 18.47451 | 1.373897 | 0.423511 | 3.244064 | 0.001178 | 0.04768 |
| AAEL006362 | 3303.221 | 0.58143 | 0.179262 | 3.243469 | 0.001181 | 0.04768 |
| AAEL001498 | 393.9792 | 1.215518 | 0.375145 | 3.240125 | 0.001195 | 0.047772 |
| AAEL027096 | 178.8959 | 0.848482 | 0.261846 | 3.240379 | 0.001194 | 0.047772 |
| AAEL002075 | 27.44772 | 1.093068 | 0.337582 | 3.23793 | 0.001204 | 0.047984 |
| AAEL014910 | 139.7114 | 0.959766 | 0.298058 | 3.220062 | 0.001282 | 0.049551 |
| AAEL009038 | 2123.108 | 0.809267 | 0.25118 | 3.22186 | 0.001274 | 0.049551 |
| AAEL014348 | 306.4843 | 0.895364 | 0.278098 | 3.219605 | 0.001284 | 0.049551 |
| AAEL027238 | 198.671 | 0.7071 | 0.219534 | 3.220908 | 0.001278 | 0.049551 |
| AAEL019751 | 742.0433 | 0.88313 | 0.274171 | 3.221094 | 0.001277 | 0.049551 |

FC: fold change; P-adj: adjusted *p* value

**3 dpi 18 ºC downregulated**

| GeneID | Base mean | log2(FC) | StdErr | Wald-Stats | P-value | P-adj |
|---|---|---|---|---|---|---|
| AAEL009165 | 438.394 | -3.08522 | 0.413725 | -7.45717 | 8.84E-14 | 6.03E-11 |
| AAEL007450 | 124.6644 | -1.55767 | 0.239976 | -6.49094 | 8.53E-11 | 4.55E-08 |
| AAEL012766 | 23.81061 | -2.08265 | 0.340855 | -6.11008 | 9.96E-10 | 4.53E-07 |
| AAEL008609 | 223.5211 | -1.9133 | 0.320578 | -5.96828 | 2.40E-09 | 9.81E-07 |
| AAEL015304 | 766.0237 | -1.18589 | 0.204076 | -5.81103 | 6.21E-09 | 2.31E-06 |
| AAEL023395 | 82.28582 | -2.09976 | 0.370658 | -5.66495 | 1.47E-08 | 5.01E-06 |
| AAEL008602 | 294.326 | -1.16002 | 0.206542 | -5.6164 | 1.95E-08 | 6.10E-06 |
| AAEL012884 | 640.7386 | -0.79055 | 0.140843 | -5.61299 | 1.99E-08 | 6.10E-06 |
| AAEL024161 | 359.2593 | -2.11474 | 0.429758 | -4.92077 | 8.62E-07 | 0.000203 |
| AAEL004283 | 222.2479 | -0.81149 | 0.16569 | -4.89766 | 9.70E-07 | 0.000225 |
| AAEL007664 | 136.8199 | -1.76802 | 0.367691 | -4.80844 | 1.52E-06 | 0.000333 |
| AAEL023753 | 722.3696 | -1.44348 | 0.311751 | -4.63023 | 3.65E-06 | 0.000701 |
| AAEL001607 | 1074.553 | -1.23204 | 0.268854 | -4.58257 | 4.59E-06 | 0.000867 |
| AAEL001693 | 180.6399 | -1.83209 | 0.402402 | -4.5529 | 5.29E-06 | 0.000969 |
| AAEL023158 | 127.7007 | -0.88786 | 0.195577 | -4.53971 | 5.63E-06 | 0.001017 |
| AAEL021138 | 278.383 | -1.24699 | 0.276271 | -4.51364 | 6.37E-06 | 0.001134 |
| AAEL009214 | 1221.181 | -1.36277 | 0.304488 | -4.47563 | 7.62E-06 | 0.001336 |
| AAEL012717 | 39.94538 | -1.92836 | 0.432801 | -4.45554 | 8.37E-06 | 0.001418 |
| AAEL007317 | 180.7115 | -0.84021 | 0.190458 | -4.41152 | 1.03E-05 | 0.001658 |
| AAEL027166 | 678.8159 | -1.79064 | 0.40985 | -4.36901 | 1.25E-05 | 0.001964 |

| | | | | | | |
|---|---|---|---|---|---|---|
| AAEL014864 | 127.8909 | -1.01439 | 0.232407 | -4.36473 | 1.27E-05 | 0.001978 |
| AAEL004313 | 1195.406 | -1.6177 | 0.371598 | -4.35337 | 1.34E-05 | 0.002057 |
| AAEL008701 | 223.0075 | -1.81708 | 0.420361 | -4.32266 | 1.54E-05 | 0.002308 |
| AAEL009843 | 60.57035 | -1.81209 | 0.420225 | -4.31218 | 1.62E-05 | 0.002391 |
| AAEL000500 | 1787.485 | -1.68356 | 0.392061 | -4.29413 | 1.75E-05 | 0.002563 |
| AAEL003443 | 318.847 | -1.78304 | 0.415707 | -4.28917 | 1.79E-05 | 0.00259 |
| AAEL013118 | 3375.385 | -1.81879 | 0.424468 | -4.28488 | 1.83E-05 | 0.00261 |
| AAEL008063 | 581.9485 | -0.61889 | 0.14459 | -4.28034 | 1.87E-05 | 0.002633 |
| AAEL004700 | 112.9155 | -1.20243 | 0.284015 | -4.2337 | 2.30E-05 | 0.003135 |
| AAEL003601 | 577.9445 | -1.23636 | 0.295569 | -4.183 | 2.88E-05 | 0.003757 |
| AAEL008876 | 274.1447 | -1.61046 | 0.387061 | -4.16073 | 3.17E-05 | 0.004099 |
| AAEL003100 | 80.70409 | -1.66985 | 0.401941 | -4.15446 | 3.26E-05 | 0.004169 |
| AAEL024064 | 52.7001 | -1.79892 | 0.433341 | -4.15127 | 3.31E-05 | 0.004184 |
| AAEL011766 | 48.33218 | -1.05116 | 0.253682 | -4.14363 | 3.42E-05 | 0.004282 |
| AAEL008561 | 707.1124 | -0.60988 | 0.147605 | -4.13186 | 3.60E-05 | 0.004373 |
| AAEL024926 | 5.064363 | -1.77784 | 0.432335 | -4.11218 | 3.92E-05 | 0.004717 |
| AAEL010196 | 4565.047 | -1.74898 | 0.425987 | -4.10571 | 4.03E-05 | 0.004804 |
| AAEL019650 | 534.3518 | -1.15792 | 0.284307 | -4.07277 | 4.65E-05 | 0.005431 |
| AAEL015458 | 2057.041 | -1.68346 | 0.424537 | -3.96541 | 7.33E-05 | 0.007754 |
| AAEL002904 | 35.40926 | -1.19036 | 0.299887 | -3.96936 | 7.21E-05 | 0.007754 |
| AAEL012932 | 1730.283 | -0.73613 | 0.185405 | -3.97039 | 7.18E-05 | 0.007754 |
| AAEL013712 | 1776.191 | -1.71109 | 0.430227 | -3.97719 | 6.97E-05 | 0.007754 |
| AAEL005008 | 774.5354 | -1.55791 | 0.393108 | -3.96307 | 7.40E-05 | 0.007763 |
| AAEL025199 | 2331.33 | -1.41866 | 0.35822 | -3.96029 | 7.49E-05 | 0.007787 |
| AAEL000146 | 32.40373 | -1.29028 | 0.327616 | -3.93838 | 8.20E-05 | 0.008322 |
| AAEL013554 | 76.32456 | -1.57441 | 0.400667 | -3.92947 | 8.51E-05 | 0.008496 |
| AAEL013001 | 139.4364 | -1.56577 | 0.399543 | -3.91889 | 8.90E-05 | 0.008806 |
| AAEL021086 | 47.59014 | -1.30114 | 0.332338 | -3.91512 | 9.04E-05 | 0.008852 |
| AAEL003182 | 543.8475 | -1.27316 | 0.3267 | -3.89703 | 9.74E-05 | 0.009172 |
| AAEL006381 | 741.8508 | -1.66804 | 0.429611 | -3.88267 | 0.000103 | 0.009262 |
| AAEL017056 | 5.354633 | -1.69349 | 0.435699 | -3.88685 | 0.000102 | 0.009262 |
| AAEL019834 | 509.3668 | -1.44168 | 0.370473 | -3.89147 | 9.96E-05 | 0.009262 |
| AAEL011741 | 2328.323 | -0.83752 | 0.2157 | -3.88281 | 0.000103 | 0.009262 |
| AAEL003742 | 287.8265 | -0.9903 | 0.256044 | -3.8677 | 0.00011 | 0.009503 |

| | | | | | | |
|---|---|---|---|---|---|---|
| AAEL010276 | 235.6529 | -1.66376 | 0.430305 | -3.86645 | 0.00011 | 0.009503 |
| AAEL006903 | 59.28145 | -1.65685 | 0.430071 | -3.8525 | 0.000117 | 0.00983 |
| AAEL000757 | 854.9099 | -1.00989 | 0.26288 | -3.84162 | 0.000122 | 0.010137 |
| AAEL000471 | 759.0046 | -1.48929 | 0.388222 | -3.83619 | 0.000125 | 0.01018 |
| AAEL008488 | 40.27667 | -1.35145 | 0.35603 | -3.79589 | 0.000147 | 0.011803 |
| AAEL024475 | 203.5882 | -1.14069 | 0.30366 | -3.75648 | 0.000172 | 0.013646 |
| AAEL018216 | 67.65922 | -0.90195 | 0.24131 | -3.73773 | 0.000186 | 0.014611 |
| AAEL002775 | 124.9256 | -0.69675 | 0.187053 | -3.72487 | 0.000195 | 0.015171 |
| AAEL003066 | 5406.107 | -1.44144 | 0.387124 | -3.72345 | 0.000197 | 0.015171 |
| AAEL006615 | 110.1771 | -1.60289 | 0.433354 | -3.69881 | 0.000217 | 0.016018 |
| AAEL003600 | 1127.051 | -1.08632 | 0.294022 | -3.6947 | 0.00022 | 0.016182 |
| AAEL013421 | 111.7923 | -1.29698 | 0.353452 | -3.66946 | 0.000243 | 0.017448 |
| AAEL012318 | 262.092 | -1.33428 | 0.366013 | -3.64544 | 0.000267 | 0.018758 |
| AAEL015336 | 488.5393 | -1.07378 | 0.294591 | -3.64497 | 0.000267 | 0.018758 |
| AAEL008751 | 69.78643 | -1.30677 | 0.358706 | -3.64302 | 0.000269 | 0.018793 |
| AAEL024221 | 5.345853 | -1.55643 | 0.429452 | -3.62422 | 0.00029 | 0.019765 |
| AAEL006347 | 1959.661 | -1.34767 | 0.372936 | -3.61368 | 0.000302 | 0.020361 |
| AAEL006454 | 68.70137 | -1.15657 | 0.320311 | -3.61077 | 0.000305 | 0.020478 |
| AAEL028002 | 9.806623 | -1.28049 | 0.355546 | -3.60148 | 0.000316 | 0.02077 |
| AAEL004676 | 329.8385 | -1.03017 | 0.285967 | -3.60239 | 0.000315 | 0.02077 |
| AAEL007668 | 478.7056 | -1.33325 | 0.373241 | -3.57207 | 0.000354 | 0.022525 |
| AAEL004301 | 431.6955 | -1.0681 | 0.29933 | -3.56831 | 0.000359 | 0.022734 |
| AAEL004097 | 326.8902 | -0.9176 | 0.257328 | -3.56587 | 0.000363 | 0.022828 |
| AAEL003806 | 71.00272 | -1.55063 | 0.435198 | -3.56306 | 0.000367 | 0.022957 |
| AAEL006483 | 663.1591 | -1.53862 | 0.433829 | -3.54661 | 0.00039 | 0.024315 |
| AAEL020477 | 634.2157 | -0.86628 | 0.245056 | -3.53505 | 0.000408 | 0.025275 |
| AAEL006333 | 403.5089 | -1.28681 | 0.364248 | -3.53278 | 0.000411 | 0.025365 |
| AAEL000223 | 201.1629 | -1.43697 | 0.407537 | -3.526 | 0.000422 | 0.025783 |
| AAEL024122 | 49.64152 | -1.25495 | 0.357578 | -3.50958 | 0.000449 | 0.026874 |
| AAEL015450 | 1765.982 | -1.31602 | 0.375199 | -3.50752 | 0.000452 | 0.026944 |
| AAEL004728 | 1624.614 | -1.187 | 0.339004 | -3.50143 | 0.000463 | 0.027179 |
| AAEL021513 | 118.4388 | -1.04908 | 0.30041 | -3.49217 | 0.000479 | 0.028005 |
| AAEL003722 | 104.7567 | -0.84764 | 0.243565 | -3.48015 | 0.000501 | 0.029016 |
| AAEL028236 | 47.74142 | -1.10188 | 0.316866 | -3.47744 | 0.000506 | 0.029066 |

| | | | | | | |
|---|---|---|---|---|---|---|
| AAEL008769 | 502.7905 | -1.48734 | 0.429667 | -3.4616 | 0.000537 | 0.029961 |
| AAEL023844 | 208.2892 | -1.01111 | 0.293357 | -3.44669 | 0.000568 | 0.030874 |
| AAEL004931 | 1059.659 | -1.44743 | 0.420047 | -3.44588 | 0.000569 | 0.030874 |
| AAEL000885 | 180.1278 | -0.76649 | 0.22249 | -3.44505 | 0.000571 | 0.030874 |
| AAEL021180 | 38.37013 | -1.19695 | 0.348403 | -3.43554 | 0.000591 | 0.031567 |
| AAEL018125 | 45.97186 | -1.43962 | 0.420605 | -3.42274 | 0.00062 | 0.032783 |
| AAEL007926 | 783.3932 | -1.29407 | 0.378206 | -3.42162 | 0.000623 | 0.032783 |
| AAEL001674 | 396.8265 | -1.44401 | 0.423314 | -3.4112 | 0.000647 | 0.033012 |
| AAEL002185 | 143.3106 | -1.15058 | 0.337187 | -3.4123 | 0.000644 | 0.033012 |
| AAEL003318 | 646.4753 | -1.37486 | 0.402531 | -3.41554 | 0.000637 | 0.033012 |
| AAEL013054 | 89.28896 | -0.70108 | 0.205719 | -3.40795 | 0.000655 | 0.0332 |
| AAEL010318 | 96968.52 | -0.70391 | 0.206996 | -3.40061 | 0.000672 | 0.033964 |
| AAEL010235 | 1711.07 | -1.14545 | 0.337265 | -3.39631 | 0.000683 | 0.034081 |
| AAEL013375 | 6.526511 | -1.446 | 0.42573 | -3.39653 | 0.000682 | 0.034081 |
| AAEL002801 | 162.5786 | -1.02585 | 0.302658 | -3.38948 | 0.0007 | 0.034377 |
| AAEL005776 | 75.35789 | -0.9483 | 0.27984 | -3.38871 | 0.000702 | 0.034377 |
| AAEL019698 | 13.24113 | -1.36899 | 0.40402 | -3.38843 | 0.000703 | 0.034377 |
| AAEL026023 | 258.7575 | -1.38667 | 0.409744 | -3.38424 | 0.000714 | 0.03463 |
| AAEL007907 | 160.1749 | -0.84771 | 0.250468 | -3.38451 | 0.000713 | 0.03463 |
| AAEL008773 | 1346.675 | -0.89811 | 0.266904 | -3.36492 | 0.000766 | 0.036383 |
| AAEL013276 | 45.17908 | -1.33383 | 0.396792 | -3.36154 | 0.000775 | 0.036383 |
| AAEL000689 | 44.09866 | -1.31763 | 0.392774 | -3.35469 | 0.000795 | 0.037084 |
| AAEL025667 | 299.8847 | -1.44163 | 0.429911 | -3.35332 | 0.000798 | 0.037126 |
| AAEL023431 | 7.05051 | -1.29718 | 0.387656 | -3.34622 | 0.000819 | 0.037946 |
| AAEL008753 | 509.4985 | -1.42447 | 0.426082 | -3.34319 | 0.000828 | 0.03803 |
| AAEL009449 | 175.5888 | -1.26647 | 0.38027 | -3.33044 | 0.000867 | 0.039421 |
| AAEL023874 | 15.27692 | -1.34954 | 0.405863 | -3.32511 | 0.000884 | 0.039974 |
| AAEL019588 | 119.2231 | -0.96967 | 0.291674 | -3.3245 | 0.000886 | 0.039974 |
| AAEL024149 | 5.098964 | -1.42067 | 0.428807 | -3.31308 | 0.000923 | 0.041339 |
| AAEL012687 | 68.23247 | -1.33568 | 0.405288 | -3.29563 | 0.000982 | 0.043051 |
| AAEL008424 | 739.1683 | -0.96785 | 0.293938 | -3.29272 | 0.000992 | 0.043161 |
| AAEL003483 | 643.4096 | -1.39197 | 0.423096 | -3.28996 | 0.001002 | 0.043161 |
| AAEL002656 | 402.7658 | -0.99556 | 0.30252 | -3.29088 | 0.000999 | 0.043161 |
| AAEL014830 | 385.1722 | -1.38864 | 0.422694 | -3.28521 | 0.001019 | 0.043738 |

| | | | | | | |
|---|---|---|---|---|---|---|
| AAEL008141 | 853.4426 | -0.62107 | 0.18939 | -3.2793 | 0.001041 | 0.044355 |
| AAEL002675 | 193.839 | -1.41964 | 0.43377 | -3.27279 | 0.001065 | 0.045075 |
| AAEL007669 | 54.41489 | -1.4053 | 0.429402 | -3.2727 | 0.001065 | 0.045075 |
| AAEL005199 | 90.53596 | -1.31955 | 0.40354 | -3.26995 | 0.001076 | 0.045208 |
| AAEL017212 | 7682.897 | -0.71193 | 0.217779 | -3.26905 | 0.001079 | 0.045208 |
| AAEL021576 | 33.63242 | -0.99791 | 0.305587 | -3.26555 | 0.001093 | 0.045615 |
| AAEL008860 | 221.8753 | -0.70897 | 0.217207 | -3.26404 | 0.001098 | 0.045703 |
| AAEL006587 | 288.8305 | -0.74326 | 0.227802 | -3.26276 | 0.001103 | 0.045754 |
| AAEL006406 | 1720.222 | -1.0076 | 0.310897 | -3.24096 | 0.001191 | 0.047772 |
| AAEL017320 | 134.5747 | -1.38428 | 0.428618 | -3.22963 | 0.00124 | 0.049042 |
| AAEL024146 | 41.99415 | -1.39058 | 0.431907 | -3.21963 | 0.001284 | 0.049551 |
| AAEL000101 | 293.866 | -1.01511 | 0.315583 | -3.21662 | 0.001297 | 0.049912 |

FC: fold change; P-adj: adjusted *p* value

**3 dpi 28 ºC upregulated**

| GeneID | Base mean | log2(FC) | StdErr | Wald-Stats | P-value | P-adj |
|---|---|---|---|---|---|---|
| AAEL013350 | 5784.056 | 7.656931 | 0.323131 | 23.69607 | 3.96E-124 | 5.02E-120 |
| AAEL013339 | 190.3038 | 6.206891 | 0.30999 | 20.02289 | 3.48E-89 | 2.21E-85 |
| AAEL017975 | 7345.993 | 5.90406 | 0.333718 | 17.69179 | 4.85E-70 | 2.05E-66 |
| AAEL017976 | 6584.064 | 6.062034 | 0.368852 | 16.43487 | 1.08E-60 | 3.41E-57 |
| AAEL020330 | 1361.706 | 6.110232 | 0.37674 | 16.21868 | 3.72E-59 | 9.43E-56 |
| AAEL013346 | 1155.747 | 5.574374 | 0.375043 | 14.86329 | 5.71E-50 | 1.21E-46 |
| AAEL013348 | 421.3866 | 4.765501 | 0.331875 | 14.35932 | 9.31E-47 | 1.69E-43 |
| AAEL023321 | 283.7587 | 3.400928 | 0.251276 | 13.53461 | 9.77E-42 | 1.55E-38 |
| AAEL026215 | 1692.454 | 1.37514 | 0.106498 | 12.91241 | 3.83E-38 | 4.88E-35 |
| AAEL022253 | 1987.402 | 4.84703 | 0.375388 | 12.91204 | 3.85E-38 | 4.88E-35 |
| AAEL024512 | 174.4702 | 3.730291 | 0.31059 | 12.01032 | 3.14E-33 | 3.61E-30 |
| AAEL006883 | 1027.71 | 2.569106 | 0.214682 | 11.96702 | 5.29E-33 | 5.59E-30 |
| AAEL019751 | 901.0433 | 2.227562 | 0.19015 | 11.71475 | 1.07E-31 | 1.04E-28 |
| AAEL027610 | 6435.418 | 4.688384 | 0.410716 | 11.41516 | 3.51E-30 | 3.18E-27 |
| AAEL013351 | 1269.289 | 3.296215 | 0.30531 | 10.7963 | 3.58E-27 | 3.03E-24 |
| AAEL008622 | 70.84556 | 3.009712 | 0.286288 | 10.51289 | 7.53E-26 | 5.97E-23 |
| AAEL026751 | 9525.754 | 2.211781 | 0.211952 | 10.43528 | 1.71E-25 | 1.28E-22 |
| AAEL025531 | 65.35001 | 3.261551 | 0.342989 | 9.509189 | 1.92E-21 | 1.35E-18 |

| | | | | | | |
|---|---|---|---|---|---|---|
| AAEL023591 | 2776.274 | 1.985301 | 0.210816 | 9.417219 | 4.63E-21 | 3.09E-18 |
| AAEL011038 | 923.8275 | 1.967118 | 0.211192 | 9.314348 | 1.23E-20 | 7.78E-18 |
| AAEL010068 | 1679.027 | 2.230035 | 0.242467 | 9.197262 | 3.67E-20 | 2.22E-17 |
| AAEL026833 | 1376.325 | 2.809921 | 0.307319 | 9.143345 | 6.05E-20 | 3.49E-17 |
| AAEL013349 | 800.6453 | 3.593085 | 0.393457 | 9.132086 | 6.72E-20 | 3.70E-17 |
| AAEL003505 | 3237.838 | 2.219512 | 0.247298 | 8.975035 | 2.83E-19 | 1.50E-16 |
| AAEL026008 | 4188.54 | 2.715168 | 0.311314 | 8.721637 | 2.74E-18 | 1.39E-15 |
| AAEL006794 | 4168.475 | 1.017043 | 0.118469 | 8.584925 | 9.09E-18 | 4.43E-15 |
| AAEL022059 | 3368.455 | 3.626993 | 0.423239 | 8.569612 | 1.04E-17 | 4.87E-15 |
| AAEL005992 | 152.9939 | 2.341997 | 0.274841 | 8.521268 | 1.58E-17 | 7.14E-15 |
| AAEL010680 | 70.01603 | 2.522056 | 0.296131 | 8.51668 | 1.64E-17 | 7.18E-15 |
| AAEL002467 | 7464.032 | 3.451633 | 0.416294 | 8.291341 | 1.12E-16 | 4.73E-14 |
| AAEL003728 | 458.8365 | 2.612568 | 0.327448 | 7.978577 | 1.48E-15 | 6.05E-13 |
| AAEL013347 | 1090.537 | 2.140365 | 0.275172 | 7.778275 | 7.35E-15 | 2.91E-12 |
| AAEL001857 | 219.3792 | 2.584523 | 0.333103 | 7.758935 | 8.56E-15 | 3.29E-12 |
| AAEL021302 | 460.6418 | 1.787031 | 0.230846 | 7.741223 | 9.85E-15 | 3.67E-12 |
| AAEL001794 | 1467.342 | 1.423436 | 0.19009 | 7.488224 | 6.98E-14 | 2.53E-11 |
| AAEL009762 | 310.1128 | 2.801133 | 0.380186 | 7.367798 | 1.73E-13 | 6.11E-11 |
| AAEL003726 | 134.2793 | 2.865447 | 0.392324 | 7.303776 | 2.80E-13 | 9.46E-11 |
| AAEL011371 | 5108.795 | 1.269037 | 0.173792 | 7.302035 | 2.83E-13 | 9.46E-11 |
| AAEL004870 | 217.5289 | 1.889874 | 0.262189 | 7.208052 | 5.68E-13 | 1.84E-10 |
| AAEL021072 | 57.51679 | 2.684856 | 0.3752 | 7.155806 | 8.32E-13 | 2.64E-10 |
| AAEL009171 | 3057.94 | 1.660286 | 0.232613 | 7.137558 | 9.50E-13 | 2.94E-10 |
| AAEL015631 | 1068.381 | 1.803133 | 0.260341 | 6.926052 | 4.33E-12 | 1.31E-09 |
| AAEL017380 | 161.5912 | 2.839763 | 0.413419 | 6.868969 | 6.47E-12 | 1.86E-09 |
| AAEL025126 | 274.4642 | 2.810271 | 0.415808 | 6.758581 | 1.39E-11 | 3.93E-09 |
| AAEL013984 | 1618.659 | 1.47697 | 0.219324 | 6.734186 | 1.65E-11 | 4.54E-09 |
| AAEL021795 | 4203.723 | 2.428758 | 0.361348 | 6.721388 | 1.80E-11 | 4.85E-09 |
| AAEL007344 | 902.5024 | 2.380682 | 0.364705 | 6.527694 | 6.68E-11 | 1.76E-08 |
| AAEL028247 | 37.18132 | 2.325267 | 0.356505 | 6.522391 | 6.92E-11 | 1.79E-08 |
| AAEL012567 | 51.72097 | 2.160349 | 0.33344 | 6.478975 | 9.23E-11 | 2.34E-08 |
| AAEL002853 | 414.3521 | 1.426999 | 0.222454 | 6.414796 | 1.41E-10 | 3.50E-08 |
| AAEL005903 | 337.8735 | 1.939593 | 0.305302 | 6.353036 | 2.11E-10 | 5.15E-08 |
| AAEL002610 | 2168.662 | 1.641394 | 0.261188 | 6.284342 | 3.29E-10 | 7.87E-08 |

| | | | | | | |
|---|---|---|---|---|---|---|
| AAEL006795 | 321.4665 | 1.841573 | 0.29576 | 6.226574 | 4.77E-10 | 1.12E-07 |
| AAEL013352 | 798.3605 | 2.470338 | 0.398806 | 6.19434 | 5.85E-10 | 1.35E-07 |
| AAEL002655 | 142.1698 | 2.037876 | 0.33391 | 6.10307 | 1.04E-09 | 2.36E-07 |
| AAEL008607 | 3804.703 | 1.504206 | 0.247252 | 6.083701 | 1.17E-09 | 2.61E-07 |
| AAEL010769 | 935.7548 | 1.240058 | 0.206443 | 6.006778 | 1.89E-09 | 4.14E-07 |
| AAEL014531 | 905.2883 | 1.084319 | 0.180737 | 5.999426 | 1.98E-09 | 4.25E-07 |
| AAEL009645 | 6326.606 | 1.186614 | 0.198253 | 5.985353 | 2.16E-09 | 4.49E-07 |
| AAEL017334 | 4229.773 | 1.92533 | 0.321629 | 5.986191 | 2.15E-09 | 4.49E-07 |
| AAEL019564 | 531.9792 | 1.464263 | 0.244865 | 5.97989 | 2.23E-09 | 4.57E-07 |
| AAEL022079 | 181.5742 | 2.163054 | 0.362815 | 5.961863 | 2.49E-09 | 5.02E-07 |
| AAEL009952 | 172.918 | 1.50934 | 0.255248 | 5.913221 | 3.35E-09 | 6.64E-07 |
| AAEL014541 | 397.9313 | 2.022085 | 0.346435 | 5.836838 | 5.32E-09 | 1.04E-06 |
| AAEL005428 | 6536.217 | 2.116166 | 0.362993 | 5.829777 | 5.55E-09 | 1.07E-06 |
| AAEL012712 | 227.1891 | 1.745037 | 0.299754 | 5.821569 | 5.83E-09 | 1.10E-06 |
| AAEL008106 | 812.1433 | 1.492137 | 0.257569 | 5.793154 | 6.91E-09 | 1.29E-06 |
| AAEL002757 | 25.01229 | 2.039375 | 0.354207 | 5.757581 | 8.53E-09 | 1.57E-06 |
| AAEL023999 | 15.79765 | 1.933722 | 0.336125 | 5.752984 | 8.77E-09 | 1.59E-06 |
| AAEL002969 | 989.6688 | 2.008806 | 0.350082 | 5.738108 | 9.57E-09 | 1.71E-06 |
| AAEL009055 | 512.7365 | 1.758351 | 0.306746 | 5.732279 | 9.91E-09 | 1.74E-06 |
| AAEL014348 | 185.223 | 2.002745 | 0.349573 | 5.729125 | 1.01E-08 | 1.75E-06 |
| AAEL000811 | 315.0489 | 1.503285 | 0.262761 | 5.721119 | 1.06E-08 | 1.81E-06 |
| AAEL004090 | 2135.773 | 1.759188 | 0.312888 | 5.622413 | 1.88E-08 | 3.18E-06 |
| AAEL007624 | 818.8665 | 1.424095 | 0.254414 | 5.597555 | 2.17E-08 | 3.63E-06 |
| AAEL006708 | 81.64057 | 1.883651 | 0.338754 | 5.560519 | 2.69E-08 | 4.43E-06 |
| AAEL002075 | 44.26663 | 2.035739 | 0.367087 | 5.545652 | 2.93E-08 | 4.76E-06 |
| AAEL003740 | 279.2502 | 1.904356 | 0.346661 | 5.49342 | 3.94E-08 | 6.25E-06 |
| AAEL010411 | 6156.341 | 0.808247 | 0.147511 | 5.479229 | 4.27E-08 | 6.69E-06 |
| AAEL006323 | 508.8304 | 1.965614 | 0.362572 | 5.421304 | 5.92E-08 | 9.15E-06 |
| AAEL008936 | 564.0859 | 1.602409 | 0.296276 | 5.408497 | 6.36E-08 | 9.71E-06 |
| AAEL021929 | 262.6133 | 2.041578 | 0.3788 | 5.389598 | 7.06E-08 | 1.07E-05 |
| AAEL001100 | 759.5531 | 1.845488 | 0.34436 | 5.359183 | 8.36E-08 | 1.25E-05 |
| AAEL001232 | 630.6709 | 1.182004 | 0.22068 | 5.356191 | 8.50E-08 | 1.25E-05 |
| AAEL010338 | 122.3551 | 1.774577 | 0.33417 | 5.310408 | 1.09E-07 | 1.59E-05 |
| AAEL017345 | 1353.901 | 1.301753 | 0.245565 | 5.301062 | 1.15E-07 | 1.66E-05 |

| | | | | | | |
|---|---|---|---|---|---|---|
| AAEL010076 | 176.9812 | 1.401221 | 0.264481 | 5.298 | 1.17E-07 | 1.67E-05 |
| AAEL028635 | 337.1032 | 2.140899 | 0.41845 | 5.116264 | 3.12E-07 | 4.39E-05 |
| AAEL001690 | 1042.88 | 2.14875 | 0.420905 | 5.105068 | 3.31E-07 | 4.61E-05 |
| AAEL007092 | 1666.175 | 0.984136 | 0.194172 | 5.068368 | 4.01E-07 | 5.52E-05 |
| AAEL022387 | 2408.535 | 1.8226 | 0.35972 | 5.066721 | 4.05E-07 | 5.52E-05 |
| AAEL001929 | 57.63435 | 1.77557 | 0.351062 | 5.057718 | 4.24E-07 | 5.72E-05 |
| AAEL001513 | 64.07459 | 1.532098 | 0.304763 | 5.027181 | 4.98E-07 | 6.64E-05 |
| AAEL008190 | 362.6617 | 1.261397 | 0.254119 | 4.963805 | 6.91E-07 | 9.13E-05 |
| AAEL002412 | 1098.94 | 1.340911 | 0.270462 | 4.957859 | 7.13E-07 | 9.31E-05 |
| AAEL005432 | 3110.994 | 1.514457 | 0.307333 | 4.927742 | 8.32E-07 | 0.0001076 |
| AAEL026819 | 103.2923 | 1.801787 | 0.367513 | 4.902648 | 9.46E-07 | 0.0001187 |
| AAEL022334 | 969.9629 | 1.652164 | 0.337944 | 4.88887 | 1.01E-06 | 0.0001254 |
| AAEL003978 | 144.74 | 1.303115 | 0.266597 | 4.887959 | 1.02E-06 | 0.0001254 |
| AAEL002124 | 735.3393 | 1.026379 | 0.210275 | 4.881121 | 1.05E-06 | 0.0001286 |
| AAEL004715 | 539.5686 | 1.268765 | 0.261237 | 4.856753 | 1.19E-06 | 0.0001441 |
| AAEL001503 | 338.8052 | 1.266419 | 0.262135 | 4.831165 | 1.36E-06 | 0.0001623 |
| AAEL005293 | 1313.266 | 1.92198 | 0.398187 | 4.826826 | 1.39E-06 | 0.0001643 |
| AAEL011203 | 106.977 | 1.869969 | 0.389089 | 4.806015 | 1.54E-06 | 0.0001807 |
| AAEL001565 | 124.8969 | 1.392703 | 0.290008 | 4.802295 | 1.57E-06 | 0.0001824 |
| AAEL004688 | 2524.55 | 0.944062 | 0.199424 | 4.733936 | 2.20E-06 | 0.0002538 |
| AAEL017139 | 187.815 | 1.911766 | 0.40473 | 4.723563 | 2.32E-06 | 0.0002647 |
| AAEL027019 | 94.76075 | 1.53148 | 0.325302 | 4.707866 | 2.50E-06 | 0.0002833 |
| AAEL010434 | 159.8025 | 1.897974 | 0.403811 | 4.700159 | 2.60E-06 | 0.0002916 |
| AAEL006990 | 1028.282 | 1.738972 | 0.371248 | 4.684128 | 2.81E-06 | 0.0003126 |
| AAEL008953 | 13309.45 | 1.170589 | 0.250402 | 4.674837 | 2.94E-06 | 0.0003243 |
| AAEL026603 | 42.51229 | 1.547599 | 0.333241 | 4.644089 | 3.42E-06 | 0.0003729 |
| AAEL012702 | 3369.073 | 1.529489 | 0.329453 | 4.642512 | 3.44E-06 | 0.0003729 |
| AAEL029047 | 47.98867 | 1.971092 | 0.426172 | 4.625108 | 3.74E-06 | 0.0004022 |
| AAEL019684 | 11.20749 | 1.747526 | 0.378185 | 4.620825 | 3.82E-06 | 0.0004071 |
| AAEL008105 | 7217.976 | 0.801024 | 0.173472 | 4.617602 | 3.88E-06 | 0.0004101 |
| AAEL026175 | 2198.047 | 1.405885 | 0.304609 | 4.615384 | 3.92E-06 | 0.000411 |
| AAEL001800 | 950.5305 | 0.935029 | 0.204034 | 4.582722 | 4.59E-06 | 0.0004769 |
| AAEL006533 | 23.2452 | 1.638814 | 0.359215 | 4.562206 | 5.06E-06 | 0.0005217 |
| AAEL014754 | 330.1854 | 1.491431 | 0.331149 | 4.503808 | 6.67E-06 | 0.0006769 |

| | | | | | | |
|---|---|---|---|---|---|---|
| AAEL011727 | 21.82691 | 1.887064 | 0.419431 | 4.499103 | 6.82E-06 | 0.0006865 |
| AAEL023348 | 338.2233 | 0.868756 | 0.19344 | 4.491087 | 7.09E-06 | 0.0007073 |
| AAEL009904 | 181.8715 | 1.884171 | 0.419793 | 4.488331 | 7.18E-06 | 0.0007109 |
| AAEL009074 | 8645.459 | 1.019847 | 0.228361 | 4.465948 | 7.97E-06 | 0.0007833 |
| AAEL007926 | 1651.043 | 1.639836 | 0.368175 | 4.453961 | 8.43E-06 | 0.000822 |
| AAEL010688 | 172.4118 | 1.386629 | 0.31161 | 4.44988 | 8.59E-06 | 0.0008314 |
| AAEL013434 | 330.6199 | 0.798817 | 0.180432 | 4.427252 | 9.54E-06 | 0.0009028 |
| AAEL015465 | 220.0909 | 1.400984 | 0.316382 | 4.428138 | 9.51E-06 | 0.0009028 |
| AAEL008473 | 2504.421 | 1.254764 | 0.284979 | 4.403007 | 1.07E-05 | 0.0009962 |
| AAEL023729 | 325.2253 | 1.876554 | 0.426262 | 4.402351 | 1.07E-05 | 0.0009962 |
| AAEL007342 | 2699.181 | 1.278658 | 0.29103 | 4.393562 | 1.12E-05 | 0.0010243 |
| AAEL000859 | 1239.467 | 1.861165 | 0.425578 | 4.373264 | 1.22E-05 | 0.0011083 |
| AAEL026531 | 121.5693 | 0.901493 | 0.206381 | 4.368094 | 1.25E-05 | 0.0011211 |
| AAEL026031 | 67.43212 | 1.114144 | 0.255089 | 4.367659 | 1.26E-05 | 0.0011211 |
| AAEL008157 | 167.8229 | 1.553542 | 0.35625 | 4.360817 | 1.30E-05 | 0.0011406 |
| AAEL006276 | 179.9901 | 1.366195 | 0.313569 | 4.356926 | 1.32E-05 | 0.0011524 |
| AAEL009850 | 96.99051 | 1.474627 | 0.338563 | 4.355547 | 1.33E-05 | 0.0011524 |
| AAEL012536 | 832.206 | 1.176214 | 0.270873 | 4.342311 | 1.41E-05 | 0.0012076 |
| AAEL017095 | 41.09661 | 1.786439 | 0.411297 | 4.343424 | 1.40E-05 | 0.0012076 |
| AAEL002353 | 84.33612 | 1.540883 | 0.355015 | 4.340331 | 1.42E-05 | 0.0012103 |
| AAEL003886 | 46.91248 | 1.722213 | 0.397482 | 4.332804 | 1.47E-05 | 0.0012441 |
| AAEL002269 | 1148.651 | 1.444293 | 0.333561 | 4.329917 | 1.49E-05 | 0.0012522 |
| AAEL010596 | 192.8158 | 1.478195 | 0.342404 | 4.317107 | 1.58E-05 | 0.0013184 |
| AAEL012698 | 394.4726 | 1.588359 | 0.368058 | 4.315511 | 1.59E-05 | 0.0013192 |
| AAEL021257 | 71.32764 | 1.376562 | 0.31922 | 4.312265 | 1.62E-05 | 0.0013301 |
| AAEL023395 | 41.78586 | 1.80173 | 0.418197 | 4.308334 | 1.64E-05 | 0.0013452 |
| AAEL027700 | 263.9977 | 1.540998 | 0.359122 | 4.29102 | 1.78E-05 | 0.0014452 |
| AAEL007197 | 1251.759 | 0.976455 | 0.228348 | 4.276166 | 1.90E-05 | 0.0015352 |
| AAEL012099 | 1309.418 | 1.021207 | 0.240332 | 4.249157 | 2.15E-05 | 0.0017107 |
| AAEL009387 | 9606.411 | 0.619584 | 0.145967 | 4.244688 | 2.19E-05 | 0.0017342 |
| AAEL001243 | 89.52798 | 1.073912 | 0.253295 | 4.239776 | 2.24E-05 | 0.0017507 |
| AAEL005342 | 799.5598 | 0.821281 | 0.193663 | 4.24078 | 2.23E-05 | 0.0017507 |
| AAEL025894 | 12.7806 | 1.802038 | 0.425469 | 4.235418 | 2.28E-05 | 0.0017738 |
| AAEL018241 | 342.8322 | 1.444882 | 0.341251 | 4.234077 | 2.29E-05 | 0.0017738 |

| | | | | | | |
|---|---|---|---|---|---|---|
| AAEL024838 | 514.8578 | 0.87682 | 0.207826 | 4.219001 | 2.45E-05 | 0.0018852 |
| AAEL001548 | 574.2691 | 1.084587 | 0.257205 | 4.21681 | 2.48E-05 | 0.0018921 |
| AAEL001965 | 1088.742 | 1.402411 | 0.332704 | 4.215188 | 2.50E-05 | 0.0018943 |
| AAEL006069 | 504.3606 | 1.30271 | 0.309209 | 4.213035 | 2.52E-05 | 0.0019011 |
| AAEL017144 | 1836.946 | 1.156814 | 0.274713 | 4.21099 | 2.54E-05 | 0.0019071 |
| AAEL002959 | 175.2784 | 1.549768 | 0.368929 | 4.200728 | 2.66E-05 | 0.0019723 |
| AAEL008096 | 512.3748 | 1.291174 | 0.310802 | 4.154332 | 3.26E-05 | 0.0023767 |
| AAEL006815 | 256.5448 | 1.087856 | 0.262123 | 4.150172 | 3.32E-05 | 0.0024065 |
| AAEL014510 | 244.9046 | 1.110525 | 0.268041 | 4.14312 | 3.43E-05 | 0.0024676 |
| AAEL007206 | 1011.047 | 1.095975 | 0.265351 | 4.130285 | 3.62E-05 | 0.0025947 |
| AAEL020603 | 54.94935 | 1.18837 | 0.288531 | 4.118684 | 3.81E-05 | 0.0027135 |
| AAEL014349 | 207.4646 | 1.071446 | 0.26027 | 4.116675 | 3.84E-05 | 0.002722 |
| AAEL006361 | 16.66943 | 1.408174 | 0.344199 | 4.091161 | 4.29E-05 | 0.003006 |
| AAEL009660 | 94.57614 | 1.152919 | 0.282251 | 4.084728 | 4.41E-05 | 0.0030735 |
| AAEL027188 | 39.3365 | 1.510878 | 0.370816 | 4.07447 | 4.61E-05 | 0.0031946 |
| AAEL001307 | 98.75584 | 1.499311 | 0.368137 | 4.072695 | 4.65E-05 | 0.0032015 |
| AAEL003640 | 370.2143 | 1.046696 | 0.257521 | 4.064503 | 4.81E-05 | 0.0032982 |
| AAEL007902 | 172.9648 | 1.16835 | 0.287866 | 4.058662 | 4.94E-05 | 0.0033636 |
| AAEL026843 | 7790.01 | 1.160491 | 0.286056 | 4.056872 | 4.97E-05 | 0.0033713 |
| AAEL019528 | 1488.322 | 0.892322 | 0.220601 | 4.044963 | 5.23E-05 | 0.0035098 |
| AAEL009531 | 203.9522 | 1.610651 | 0.399908 | 4.027552 | 5.64E-05 | 0.003721 |
| AAEL007765 | 4599.867 | 0.990208 | 0.246687 | 4.014026 | 5.97E-05 | 0.0039205 |
| AAEL011867 | 103.0433 | 1.145468 | 0.285495 | 4.012217 | 6.02E-05 | 0.0039303 |
| AAEL029031 | 190.9685 | 1.168092 | 0.291701 | 4.004412 | 6.22E-05 | 0.0040415 |
| AAEL005666 | 566.7667 | 1.052103 | 0.26297 | 4.000854 | 6.31E-05 | 0.0040818 |
| AAEL027093 | 3.850387 | 1.668462 | 0.41737 | 3.997558 | 6.40E-05 | 0.0040894 |
| AAEL003632 | 31.74144 | 1.532191 | 0.383149 | 3.998939 | 6.36E-05 | 0.0040894 |
| AAEL006754 | 177.584 | 1.065677 | 0.26671 | 3.995634 | 6.45E-05 | 0.0040894 |
| AAEL003888 | 1937.104 | 1.435117 | 0.359065 | 3.996812 | 6.42E-05 | 0.0040894 |
| AAEL013812 | 571.2525 | 1.594401 | 0.399522 | 3.990774 | 6.59E-05 | 0.0041328 |
| AAEL026447 | 4.585397 | 1.67971 | 0.421203 | 3.987887 | 6.67E-05 | 0.0041627 |
| AAEL017023 | 760.2604 | 0.825842 | 0.207597 | 3.978098 | 6.95E-05 | 0.0042955 |
| AAEL006280 | 45.33233 | 1.287139 | 0.32376 | 3.975591 | 7.02E-05 | 0.0043055 |
| AAEL009630 | 257.3899 | 1.001203 | 0.25186 | 3.975235 | 7.03E-05 | 0.0043055 |

| | | | | | | |
|---|---|---|---|---|---|---|
| AAEL010625 | 8.018426 | 1.642687 | 0.41373 | 3.970431 | 7.17E-05 | 0.0043512 |
| AAEL013341 | 1207.901 | 1.21579 | 0.306183 | 3.970794 | 7.16E-05 | 0.0043512 |
| AAEL019578 | 203.8957 | 1.193456 | 0.300841 | 3.967064 | 7.28E-05 | 0.0043921 |
| AAEL010479 | 1728.988 | 0.988333 | 0.249786 | 3.956721 | 7.60E-05 | 0.0045649 |
| AAEL014618 | 34.6267 | 1.686867 | 0.426507 | 3.955074 | 7.65E-05 | 0.0045748 |
| AAEL007792 | 278.9527 | 1.207479 | 0.306052 | 3.945344 | 7.97E-05 | 0.0047201 |
| AAEL008354 | 111.4486 | 1.057035 | 0.267885 | 3.945848 | 7.95E-05 | 0.0047201 |
| AAEL001373 | 33.94338 | 1.283716 | 0.32588 | 3.939227 | 8.17E-05 | 0.0047972 |
| AAEL014076 | 41.77342 | 1.653518 | 0.419674 | 3.940003 | 8.15E-05 | 0.0047972 |
| AAEL003737 | 172.2251 | 1.090585 | 0.277245 | 3.933651 | 8.37E-05 | 0.0048873 |
| AAEL007703 | 743.8899 | 1.651893 | 0.420655 | 3.926951 | 8.60E-05 | 0.0050023 |
| AAEL008699 | 77.33347 | 1.672026 | 0.426238 | 3.922755 | 8.75E-05 | 0.005067 |
| AAEL024560 | 92.78694 | 1.200297 | 0.308026 | 3.896736 | 9.75E-05 | 0.0056177 |
| AAEL018351 | 517.4097 | 0.966635 | 0.24819 | 3.894738 | 9.83E-05 | 0.0056385 |
| AAEL019868 | 142.1417 | 1.306783 | 0.336771 | 3.880331 | 0.0001043 | 0.0059563 |
| AAEL026300 | 36.49657 | 1.620907 | 0.418581 | 3.872383 | 0.0001078 | 0.0061263 |
| AAEL006394 | 10.63294 | 1.387161 | 0.358467 | 3.8697 | 0.000109 | 0.0061665 |
| AAEL022674 | 138.1285 | 1.458834 | 0.378489 | 3.85436 | 0.000116 | 0.006537 |
| AAEL003051 | 92.12374 | 1.288741 | 0.334764 | 3.849699 | 0.0001183 | 0.0066332 |
| AAEL008628 | 17.79751 | 1.411862 | 0.368837 | 3.827872 | 0.0001293 | 0.0071862 |
| AAEL003857 | 72.99003 | 1.604194 | 0.419271 | 3.82615 | 0.0001302 | 0.007205 |
| AAEL027699 | 13.28489 | 1.386421 | 0.363289 | 3.816303 | 0.0001355 | 0.0073558 |
| AAEL007090 | 256.6072 | 1.298519 | 0.340496 | 3.813614 | 0.0001369 | 0.0073558 |
| AAEL024038 | 138.8779 | 1.010885 | 0.265026 | 3.814294 | 0.0001366 | 0.0073558 |
| AAEL023882 | 10.13949 | 1.549059 | 0.405857 | 3.816764 | 0.0001352 | 0.0073558 |
| AAEL011980 | 130.3171 | 1.120688 | 0.294684 | 3.803022 | 0.0001429 | 0.0075912 |
| AAEL027632 | 623.31 | 1.436387 | 0.377728 | 3.802699 | 0.0001431 | 0.0075912 |
| AAEL020340 | 7667.399 | 0.903366 | 0.238488 | 3.787893 | 0.0001519 | 0.0080244 |
| AAEL002175 | 2093.977 | 0.587213 | 0.155435 | 3.777878 | 0.0001582 | 0.0083194 |
| AAEL025750 | 156.3401 | 1.21716 | 0.323445 | 3.763119 | 0.0001678 | 0.0086822 |
| AAEL003527 | 60.39437 | 1.041641 | 0.276793 | 3.763242 | 0.0001677 | 0.0086822 |
| AAEL013257 | 82.30367 | 1.165331 | 0.309554 | 3.764553 | 0.0001668 | 0.0086822 |
| AAEL025839 | 30.1891 | 1.555673 | 0.413083 | 3.766007 | 0.0001659 | 0.0086822 |
| AAEL005491 | 235.1988 | 1.077765 | 0.28707 | 3.754358 | 0.0001738 | 0.0088827 |

| | | | | | | |
|---|---|---|---|---|---|---|
| AAEL001519 | 144.9248 | 1.34742 | 0.358811 | 3.755233 | 0.0001732 | 0.0088827 |
| AAEL002254 | 591.4642 | 0.876364 | 0.233521 | 3.752835 | 0.0001748 | 0.008901 |
| AAEL014773 | 7.563296 | 1.387879 | 0.370479 | 3.746172 | 0.0001796 | 0.0090817 |
| AAEL003389 | 37.29007 | 1.578976 | 0.421534 | 3.745788 | 0.0001798 | 0.0090817 |
| AAEL029046 | 13.03295 | 1.559203 | 0.416459 | 3.743951 | 0.0001811 | 0.0091121 |
| AAEL025552 | 100.2473 | 1.224406 | 0.327524 | 3.738373 | 0.0001852 | 0.0092433 |
| AAEL012859 | 101.3332 | 1.388411 | 0.371717 | 3.735133 | 0.0001876 | 0.0093264 |
| AAEL009126 | 41.73404 | 1.589443 | 0.426507 | 3.726651 | 0.000194 | 0.0095707 |
| AAEL012856 | 399.5885 | 1.165543 | 0.313075 | 3.72289 | 0.000197 | 0.0096768 |
| AAEL009487 | 3417.461 | 1.014725 | 0.27331 | 3.712721 | 0.000205 | 0.0099583 |
| AAEL001417 | 229.3783 | 1.326564 | 0.357297 | 3.712779 | 0.000205 | 0.0099583 |
| AAEL028221 | 66.42982 | 1.059166 | 0.285634 | 3.708123 | 0.0002088 | 0.0100637 |
| AAEL012184 | 150.9025 | 0.996729 | 0.269675 | 3.696034 | 0.000219 | 0.0104754 |
| AAEL014316 | 53.42481 | 1.162642 | 0.314961 | 3.691385 | 0.000223 | 0.0106286 |
| AAEL013600 | 815.7912 | 0.75663 | 0.205186 | 3.687537 | 0.0002264 | 0.01071 |
| AAEL020033 | 453.895 | 1.239684 | 0.336099 | 3.688453 | 0.0002256 | 0.01071 |
| AAEL015644 | 237.9285 | 1.10361 | 0.299777 | 3.681441 | 0.0002319 | 0.0108882 |
| AAEL007322 | 1890.898 | 0.954807 | 0.259324 | 3.681912 | 0.0002315 | 0.0108882 |
| AAEL006114 | 7.660662 | 1.513053 | 0.412564 | 3.667436 | 0.000245 | 0.0114596 |
| AAEL014539 | 48.80451 | 1.38572 | 0.37831 | 3.662926 | 0.0002494 | 0.0115779 |
| AAEL021583 | 474.0557 | 1.154128 | 0.316011 | 3.65218 | 0.00026 | 0.0120294 |
| AAEL025392 | 10.03591 | 1.496365 | 0.41126 | 3.638491 | 0.0002742 | 0.0125497 |
| AAEL026537 | 13.80957 | 1.257432 | 0.346143 | 3.632699 | 0.0002805 | 0.0127887 |
| AAEL006126 | 93.96053 | 1.502308 | 0.413837 | 3.630191 | 0.0002832 | 0.0128673 |
| AAEL020078 | 15.49207 | 1.341617 | 0.369746 | 3.628487 | 0.0002851 | 0.0129063 |
| AAEL012091 | 149.8871 | 0.844295 | 0.233129 | 3.621572 | 0.0002928 | 0.0131623 |
| AAEL026440 | 1936.124 | 1.064952 | 0.294319 | 3.618365 | 0.0002965 | 0.0132794 |
| AAEL009837 | 408.4249 | 1.014612 | 0.280807 | 3.613194 | 0.0003024 | 0.0134994 |
| AAEL005130 | 66.79305 | 0.914671 | 0.253274 | 3.61139 | 0.0003046 | 0.013546 |
| AAEL014343 | 22.30363 | 1.267359 | 0.352035 | 3.600088 | 0.0003181 | 0.01405 |
| AAEL025125 | 5.258394 | 1.532462 | 0.426213 | 3.595532 | 0.0003237 | 0.0142486 |
| AAEL009249 | 296.5556 | 1.05174 | 0.294025 | 3.577048 | 0.0003475 | 0.0150343 |
| AAEL002964 | 579.7919 | 0.95955 | 0.268206 | 3.577665 | 0.0003467 | 0.0150343 |
| AAEL003420 | 140.9826 | 1.049211 | 0.293817 | 3.570966 | 0.0003557 | 0.0153348 |

| | | | | | | |
|---|---|---|---|---|---|---|
| AAEL024468 | 7.567725 | 1.517422 | 0.425983 | 3.562164 | 0.0003678 | 0.0158047 |
| AAEL001536 | 30.53871 | 1.240871 | 0.348656 | 3.559012 | 0.0003723 | 0.0159415 |
| AAEL013875 | 5029.768 | 0.899848 | 0.252949 | 3.557425 | 0.0003745 | 0.015984 |
| AAEL009148 | 332.5632 | 1.042286 | 0.293367 | 3.55284 | 0.0003811 | 0.0161565 |
| AAEL027655 | 13.87487 | 1.330922 | 0.374997 | 3.549156 | 0.0003865 | 0.0163296 |
| AAEL005768 | 447.8671 | 1.361932 | 0.384271 | 3.544191 | 0.0003938 | 0.0165849 |
| AAEL007403 | 236.0034 | 0.961803 | 0.272173 | 3.533787 | 0.0004097 | 0.0171377 |
| AAEL001818 | 387.54 | 1.117286 | 0.31656 | 3.529456 | 0.0004164 | 0.0173634 |
| AAEL021016 | 200.1772 | 0.772463 | 0.219288 | 3.522597 | 0.0004273 | 0.0177605 |
| AAEL024540 | 137.1792 | 0.991282 | 0.281532 | 3.521032 | 0.0004299 | 0.0178073 |
| AAEL007502 | 320.1961 | 0.846843 | 0.241569 | 3.505591 | 0.0004556 | 0.0188114 |
| AAEL010881 | 80.64011 | 1.212936 | 0.346136 | 3.504221 | 0.0004579 | 0.0188471 |
| AAEL010451 | 73.26203 | 1.116715 | 0.319472 | 3.495501 | 0.0004732 | 0.0193482 |
| AAEL001928 | 1682.96 | 1.018857 | 0.291817 | 3.491427 | 0.0004804 | 0.0195825 |
| AAEL021982 | 17.62089 | 1.166193 | 0.334432 | 3.487086 | 0.0004883 | 0.019776 |
| AAEL008489 | 242.942 | 0.967137 | 0.27772 | 3.48242 | 0.0004969 | 0.0200555 |
| AAEL002390 | 851.7374 | 1.068684 | 0.30695 | 3.481624 | 0.0004984 | 0.0200555 |
| AAEL024669 | 730.7402 | 1.029516 | 0.2965 | 3.472226 | 0.0005162 | 0.0207053 |
| AAEL001693 | 176.3984 | 1.406068 | 0.405565 | 3.466939 | 0.0005264 | 0.0210502 |
| AAEL014910 | 30.89665 | 1.34186 | 0.388051 | 3.457945 | 0.0005443 | 0.0216292 |
| AAEL006138 | 142.0151 | 1.469892 | 0.425706 | 3.452833 | 0.0005547 | 0.0219059 |
| AAEL023243 | 28.10678 | 1.117499 | 0.324289 | 3.446003 | 0.0005689 | 0.0223973 |
| AAEL022600 | 400.9745 | 1.363779 | 0.396013 | 3.443773 | 0.0005737 | 0.0225082 |
| AAEL014138 | 164.986 | 0.802264 | 0.233013 | 3.442994 | 0.0005753 | 0.0225082 |
| AAEL012412 | 22.49855 | 1.275602 | 0.370778 | 3.44034 | 0.000581 | 0.0225906 |
| AAEL004719 | 340.5183 | 0.879947 | 0.255738 | 3.440813 | 0.00058 | 0.0225906 |
| AAEL002908 | 49.15145 | 1.465278 | 0.426499 | 3.435598 | 0.0005912 | 0.0229195 |
| AAEL008404 | 19.86394 | 1.411783 | 0.412881 | 3.419349 | 0.0006277 | 0.0242588 |
| AAEL009869 | 1228.05 | 0.608743 | 0.178301 | 3.414127 | 0.0006399 | 0.0243841 |
| AAEL019844 | 58.7366 | 0.90623 | 0.265459 | 3.413827 | 0.0006406 | 0.0243841 |
| AAEL019537 | 442.6477 | 1.402509 | 0.410702 | 3.414902 | 0.0006381 | 0.0243841 |
| AAEL011779 | 149.1592 | 0.870939 | 0.255109 | 3.413989 | 0.0006402 | 0.0243841 |
| AAEL013525 | 597.8701 | 0.888488 | 0.260494 | 3.410784 | 0.0006478 | 0.024584 |
| AAEL012093 | 571.972 | 1.056681 | 0.310395 | 3.40431 | 0.0006633 | 0.025099 |

| | | | | | | |
|---|---|---|---|---|---|---|
| AAEL013345 | 2091.104 | 1.351629 | 0.397873 | 3.397132 | 0.000681 | 0.025538 |
| AAEL011967 | 912.1809 | 0.877246 | 0.258229 | 3.397158 | 0.0006809 | 0.025538 |
| AAEL024112 | 607.4509 | 1.092194 | 0.321654 | 3.395553 | 0.0006849 | 0.0256101 |
| AAEL019893 | 337.8388 | 0.60392 | 0.17792 | 3.394337 | 0.000688 | 0.0256484 |
| AAEL005849 | 8412.767 | 1.155223 | 0.340574 | 3.391991 | 0.0006939 | 0.0257931 |
| AAEL012549 | 1065.228 | 0.976689 | 0.288427 | 3.386262 | 0.0007085 | 0.0262175 |
| AAEL007693 | 311.1743 | 1.157429 | 0.341837 | 3.385913 | 0.0007094 | 0.0262175 |
| AAEL003345 | 10284.03 | 1.172598 | 0.34724 | 3.376911 | 0.000733 | 0.0269337 |
| AAEL027362 | 31.42138 | 1.391915 | 0.41233 | 3.375731 | 0.0007362 | 0.0269712 |
| AAEL027829 | 171.7545 | 1.338293 | 0.396624 | 3.374211 | 0.0007403 | 0.0270425 |
| AAEL026041 | 13.17522 | 1.33437 | 0.395992 | 3.369688 | 0.0007525 | 0.0273327 |
| AAEL008832 | 2258.953 | 0.698426 | 0.207252 | 3.369935 | 0.0007519 | 0.0273327 |
| AAEL000242 | 45.51674 | 1.024009 | 0.304451 | 3.363464 | 0.0007697 | 0.0277182 |
| AAEL008278 | 412.195 | 1.326029 | 0.394747 | 3.359186 | 0.0007817 | 0.0280712 |
| AAEL013906 | 19.18083 | 1.356959 | 0.404346 | 3.355934 | 0.000791 | 0.0283232 |
| AAEL022829 | 9.194961 | 1.414172 | 0.422182 | 3.349674 | 0.0008091 | 0.0288894 |
| AAEL025079 | 8.309693 | 1.407754 | 0.420599 | 3.347024 | 0.0008168 | 0.028928 |
| AAEL009081 | 1992.548 | 0.644463 | 0.192496 | 3.347939 | 0.0008142 | 0.028928 |
| AAEL004805 | 2452.177 | 1.189846 | 0.356103 | 3.341295 | 0.0008339 | 0.029362 |
| AAEL002130 | 2748.739 | 0.739045 | 0.221291 | 3.339691 | 0.0008387 | 0.0294504 |
| AAEL004572 | 127.7069 | 1.028086 | 0.30798 | 3.338163 | 0.0008433 | 0.029531 |
| AAEL006329 | 17.39274 | 1.202431 | 0.361327 | 3.327817 | 0.0008753 | 0.0303979 |
| AAEL027493 | 152.3785 | 0.754039 | 0.226649 | 3.326902 | 0.0008782 | 0.0304145 |
| AAEL012450 | 121.4309 | 0.868228 | 0.262584 | 3.306471 | 0.0009448 | 0.0323681 |
| AAEL005651 | 6837.951 | 0.707463 | 0.213943 | 3.306783 | 0.0009437 | 0.0323681 |
| AAEL007191 | 411.6934 | 1.08428 | 0.32835 | 3.302211 | 0.0009593 | 0.0327751 |
| AAEL013163 | 932.7747 | 1.002523 | 0.304206 | 3.29554 | 0.0009823 | 0.0333833 |
| AAEL019773 | 78.99006 | 1.398552 | 0.424672 | 3.29325 | 0.0009904 | 0.0335665 |
| AAEL000618 | 34.66495 | 1.210145 | 0.368289 | 3.285855 | 0.0010167 | 0.0341536 |
| AAEL000037 | 418.4091 | 1.093827 | 0.332938 | 3.285376 | 0.0010185 | 0.0341536 |
| AAEL009888 | 481.8975 | 0.805992 | 0.245405 | 3.284341 | 0.0010222 | 0.0341888 |
| AAEL024175 | 55.25191 | 0.911478 | 0.277895 | 3.279933 | 0.0010383 | 0.0345452 |
| AAEL002661 | 10.03927 | 1.396822 | 0.426485 | 3.275194 | 0.0010559 | 0.035038 |
| AAEL006352 | 247.1212 | 1.071658 | 0.327645 | 3.270791 | 0.0010725 | 0.0354951 |

| | | | | | | |
|---|---|---|---|---|---|---|
| AAEL001349 | 60.11651 | 0.855291 | 0.262136 | 3.262776 | 0.0011033 | 0.0361209 |
| AAEL004213 | 55.24829 | 1.285933 | 0.394022 | 3.263609 | 0.0011 | 0.0361209 |
| AAEL008668 | 44.47421 | 0.940649 | 0.288628 | 3.259041 | 0.0011179 | 0.0364279 |
| AAEL008336 | 26.22536 | 0.957256 | 0.293909 | 3.256985 | 0.001126 | 0.036505 |
| AAEL019438 | 372.2857 | 1.24651 | 0.382814 | 3.25618 | 0.0011292 | 0.0365154 |
| AAEL022427 | 44.84072 | 1.180727 | 0.36314 | 3.25144 | 0.0011482 | 0.0370352 |
| AAEL000164 | 106.7757 | 1.220664 | 0.375644 | 3.24952 | 0.001156 | 0.0371915 |
| AAEL010738 | 1982.918 | 0.822461 | 0.253202 | 3.248236 | 0.0011612 | 0.0372651 |
| AAEL027514 | 412.0888 | 0.701038 | 0.216342 | 3.240417 | 0.0011936 | 0.0381095 |
| AAEL007535 | 268.9715 | 0.807575 | 0.24955 | 3.236131 | 0.0012116 | 0.0385891 |
| AAEL000102 | 1418.612 | 0.99988 | 0.309177 | 3.234008 | 0.0012207 | 0.0387556 |
| AAEL001646 | 154.4133 | 1.290689 | 0.399209 | 3.233116 | 0.0012245 | 0.0387556 |
| AAEL000967 | 611.8377 | 0.792888 | 0.245267 | 3.232757 | 0.001226 | 0.0387556 |
| AAEL010067 | 14.60293 | 1.075311 | 0.332704 | 3.23203 | 0.0012291 | 0.0387577 |
| AAEL015416 | 9.73182 | 1.291735 | 0.400928 | 3.221862 | 0.0012736 | 0.0399963 |
| AAEL003738 | 117.2808 | 1.200916 | 0.372971 | 3.219863 | 0.0012825 | 0.0400424 |
| AAEL010413 | 4.499809 | 1.318839 | 0.40987 | 3.217696 | 0.0012922 | 0.0400494 |
| AAEL012341 | 568.3146 | 0.97098 | 0.301644 | 3.218965 | 0.0012865 | 0.0400494 |
| AAEL019849 | 447.027 | 0.79886 | 0.248324 | 3.217001 | 0.0012954 | 0.0400494 |
| AAEL007143 | 48.75739 | 1.058717 | 0.329087 | 3.217133 | 0.0012948 | 0.0400494 |
| AAEL009464 | 40.25879 | 1.103822 | 0.343521 | 3.213257 | 0.0013124 | 0.0403527 |
| AAEL006582 | 31869.16 | 0.690617 | 0.215008 | 3.212048 | 0.0013179 | 0.0403527 |
| AAEL001062 | 277.9282 | 0.948194 | 0.295525 | 3.208506 | 0.0013343 | 0.0407546 |
| AAEL015458 | 380.1295 | 1.09973 | 0.343444 | 3.202062 | 0.0013645 | 0.0415667 |
| AAEL002347 | 155.4017 | 1.063669 | 0.332247 | 3.201443 | 0.0013674 | 0.0415667 |
| AAEL023560 | 68.28717 | 0.970427 | 0.303694 | 3.195409 | 0.0013963 | 0.0421425 |
| AAEL000834 | 559.4591 | 1.144341 | 0.358232 | 3.194416 | 0.0014011 | 0.0421873 |
| AAEL022876 | 8.691477 | 1.173543 | 0.367778 | 3.190904 | 0.0014183 | 0.0425015 |
| AAEL009038 | 518.4585 | 1.011739 | 0.317871 | 3.182855 | 0.0014583 | 0.0434953 |
| AAEL027106 | 190.1763 | 1.027908 | 0.323191 | 3.180491 | 0.0014703 | 0.0437487 |
| AAEL004533 | 129.7041 | 0.914981 | 0.288245 | 3.174316 | 0.0015019 | 0.0443425 |
| AAEL003703 | 19.06008 | 1.269577 | 0.400009 | 3.173872 | 0.0015042 | 0.0443425 |
| AAEL000428 | 1438.551 | 1.014774 | 0.319595 | 3.175189 | 0.0014974 | 0.0443425 |
| AAEL006113 | 677.8402 | 0.730493 | 0.230422 | 3.170244 | 0.0015231 | 0.0447956 |

| | | | | | | |
|---|---|---|---|---|---|---|
| AAEL025332 | 10.69345 | 1.283329 | 0.40543 | 3.165353 | 0.001549 | 0.0454472 |
| AAEL008274 | 1215.456 | 0.888387 | 0.280718 | 3.164699 | 0.0015524 | 0.0454472 |
| AAEL005676 | 86.96242 | 1.143142 | 0.36143 | 3.162828 | 0.0015624 | 0.0456349 |
| AAEL003229 | 2207.826 | 0.59259 | 0.187414 | 3.161923 | 0.0015673 | 0.0456718 |
| AAEL011180 | 3243.92 | 1.028451 | 0.325333 | 3.161228 | 0.0015711 | 0.0456759 |
| AAEL019767 | 806.3366 | 0.835747 | 0.264885 | 3.155132 | 0.0016043 | 0.0465343 |
| AAEL001094 | 955.4217 | 0.693881 | 0.220024 | 3.153662 | 0.0016124 | 0.0466627 |
| AAEL005177 | 419.3467 | 0.887037 | 0.282033 | 3.145156 | 0.00166 | 0.0479315 |
| AAEL006904 | 1363.188 | 0.58096 | 0.184792 | 3.143852 | 0.0016674 | 0.0480363 |
| AAEL012135 | 286.5131 | 0.923976 | 0.294007 | 3.142698 | 0.001674 | 0.0481167 |
| AAEL027270 | 10.83758 | 1.293931 | 0.412253 | 3.13868 | 0.0016971 | 0.048561 |
| AAEL019463 | 415.8936 | 0.801131 | 0.255357 | 3.137294 | 0.0017052 | 0.0486813 |
| AAEL007993 | 21.5142 | 1.25305 | 0.399686 | 3.135083 | 0.0017181 | 0.0488298 |
| AAEL005017 | 1179.668 | 0.716334 | 0.228478 | 3.135243 | 0.0017171 | 0.0488298 |
| AAEL010379 | 496.8672 | 0.802462 | 0.256622 | 3.127014 | 0.0017659 | 0.049966 |
| AAEL001318 | 17.67618 | 1.063269 | 0.339988 | 3.127375 | 0.0017637 | 0.049966 |

FC: fold change; P-adj: adjusted *p* value

**3 dpi 28 ºC downregulated**

| GeneID | Base mean | log2(FC) | StdErr | Wald-Stats | P-value | P-adj |
|---|---|---|---|---|---|---|
| AAEL012628 | 474.5021 | -2.8039 | 0.406212 | -6.90257 | 5.11E-12 | 1.51E-09 |
| AAEL000507 | 59.32845 | -1.97832 | 0.357445 | -5.53463 | 3.12E-08 | 5.00E-06 |
| AAEL009899 | 1587.204 | -1.52068 | 0.309149 | -4.91894 | 8.70E-07 | 0.0001114 |
| AAEL010891 | 88.07134 | -1.21154 | 0.24657 | -4.91356 | 8.94E-07 | 0.0001134 |
| AAEL018189 | 59.77118 | -1.71655 | 0.39002 | -4.40117 | 1.08E-05 | 0.0009962 |
| AAEL025401 | 592.1465 | -0.86644 | 0.198485 | -4.36527 | 1.27E-05 | 0.0011255 |
| AAEL028022 | 94.78526 | -0.8702 | 0.204456 | -4.25616 | 2.08E-05 | 0.0016685 |
| AAEL022506 | 603.8726 | -0.90026 | 0.214022 | -4.20637 | 2.60E-05 | 0.001935 |
| AAEL010855 | 777.8795 | -1.40369 | 0.337708 | -4.15651 | 3.23E-05 | 0.0023677 |
| AAEL002785 | 647.5887 | -0.71852 | 0.177452 | -4.04913 | 5.14E-05 | 0.0034663 |
| AAEL020175 | 24.92202 | -1.70906 | 0.4231 | -4.03937 | 5.36E-05 | 0.0035756 |
| AAEL003916 | 2524.846 | -0.65688 | 0.163013 | -4.0296 | 5.59E-05 | 0.003708 |
| AAEL005507 | 193.9252 | -1.70281 | 0.4265 | -3.99251 | 6.54E-05 | 0.0041229 |
| AAEL011983 | 1222.265 | -0.83232 | 0.208822 | -3.98577 | 6.73E-05 | 0.0041794 |

| | | | | | | |
|---|---|---|---|---|---|---|
| AAEL006313 | 294.6973 | -1.11518 | 0.291314 | -3.82809 | 0.0001291 | 0.0071862 |
| AAEL024022 | 1257.093 | -0.66379 | 0.173974 | -3.81545 | 0.0001359 | 0.0073558 |
| AAEL014295 | 769.078 | -0.92575 | 0.243127 | -3.80767 | 0.0001403 | 0.0075029 |
| AAEL011710 | 642.353 | -0.95844 | 0.255195 | -3.75572 | 0.0001728 | 0.0088827 |
| AAEL011911 | 565.861 | -0.62327 | 0.166709 | -3.73866 | 0.000185 | 0.0092433 |
| AAEL005577 | 278.9741 | -0.76512 | 0.205044 | -3.73146 | 0.0001904 | 0.0094264 |
| AAEL014263 | 66.07671 | -1.15402 | 0.310698 | -3.71428 | 0.0002038 | 0.0099583 |
| AAEL011510 | 132.9247 | -1.06661 | 0.28816 | -3.70144 | 0.0002144 | 0.0102933 |
| AAEL012455 | 376.7221 | -0.763 | 0.208191 | -3.66488 | 0.0002475 | 0.011532 |
| AAEL006614 | 282.8583 | -0.73253 | 0.202205 | -3.62268 | 0.0002916 | 0.0131526 |
| AAEL019677 | 1164.556 | -0.90249 | 0.253846 | -3.55526 | 0.0003776 | 0.0160621 |
| AAEL012566 | 70.42868 | -1.46172 | 0.413295 | -3.53675 | 0.0004051 | 0.017003 |
| AAEL026775 | 414.3637 | -0.83955 | 0.249591 | -3.36371 | 0.000769 | 0.0277182 |
| AAEL026260 | 85.39223 | -1.1222 | 0.333522 | -3.36468 | 0.0007663 | 0.0277182 |
| AAEL022382 | 36.42032 | -1.29198 | 0.387719 | -3.33226 | 0.0008615 | 0.0300415 |
| AAEL026241 | 507.6766 | -0.59299 | 0.178602 | -3.32017 | 0.0008996 | 0.0310731 |
| AAEL022900 | 50.1307 | -1.36645 | 0.414138 | -3.29951 | 0.0009685 | 0.033003 |
| AAEL024298 | 83.20629 | -1.1401 | 0.34678 | -3.28769 | 0.0010101 | 0.0341457 |
| AAEL013816 | 1131.992 | -0.59861 | 0.182456 | -3.28084 | 0.001035 | 0.0345246 |
| AAEL021303 | 127.5461 | -1.29047 | 0.39539 | -3.26379 | 0.0010993 | 0.0361209 |
| AAEL006488 | 1316.479 | -0.58701 | 0.182219 | -3.22148 | 0.0012753 | 0.0399963 |
| AAEL025549 | 22.5495 | -0.80927 | 0.253398 | -3.19369 | 0.0014047 | 0.0421933 |

FC: fold change; P-adj: adjusted *p* value

**3 dpi 32 ºC upregulated**

| GeneID | Base mean | log2(FC) | StdErr | Wald-stats | P-value | P-adj |
|---|---|---|---|---|---|---|
| AAEL013339 | 87.53917 | 2.487519 | 0.370644 | 6.711338 | 1.93E-11 | 2.37E-07 |
| AAEL004090 | 1851.692 | 1.374891 | 0.247752 | 5.549469 | 2.87E-08 | 0.000176 |
| AAEL013349 | 673.0459 | 2.055251 | 0.377009 | 5.451465 | 5.00E-08 | 0.000205 |
| AAEL013346 | 755.9516 | 2.024833 | 0.376885 | 5.372545 | 7.76E-08 | 0.000239 |
| AAEL017976 | 6791.25 | 1.939832 | 0.374036 | 5.186217 | 2.15E-07 | 0.000442 |
| AAEL013345 | 1284.808 | 1.955582 | 0.377127 | 5.185477 | 2.15E-07 | 0.000442 |
| AAEL013348 | 219.7709 | 1.89139 | 0.3689 | 5.127108 | 2.94E-07 | 5.17E-04 |
| AAEL013351 | 775.3334 | 1.630181 | 0.321914 | 5.064033 | 4.10E-07 | 6.31E-04 |

| | | | | | | |
|---|---|---|---|---|---|---|
| AAEL020330 | 1528.054 | 1.838949 | 0.37786 | 4.866747 | 1.13E-06 | 0.001551 |
| AAEL013350 | 4300.273 | 1.82661 | 0.378096 | 4.831071 | 1.36E-06 | 0.00167 |
| AAEL006883 | 1071.751 | 1.315066 | 0.311461 | 4.222249 | 2.42E-05 | 0.021401 |
| AAEL006886 | 129.0013 | 1.504414 | 0.354119 | 4.248332 | 2.15E-05 | 0.021401 |
| AAEL013257 | 86.11675 | 1.150659 | 0.273634 | 4.205105 | 2.61E-05 | 0.021401 |
| AAEL021012 | 439.7245 | 1.43843 | 0.338622 | 4.247893 | 2.16E-05 | 0.021401 |
| AAEL010068 | 1739.129 | 1.141382 | 0.27103 | 4.211268 | 2.54E-05 | 0.021401 |
| AAEL009682 | 7.136665 | 1.547103 | 0.369836 | 4.183215 | 2.87E-05 | 0.022097 |
| AAEL010793 | 302.8245 | 1.010064 | 0.244752 | 4.126884 | 3.68E-05 | 0.023585 |
| AAEL012184 | 144.1262 | 0.885474 | 0.214867 | 4.121029 | 3.77E-05 | 0.023585 |
| AAEL009660 | 114.36 | 0.905478 | 0.219925 | 4.11722 | 3.83E-05 | 0.023585 |
| AAEL023321 | 363.3567 | 1.400596 | 0.342886 | 4.084733 | 4.41E-05 | 0.025848 |
| AAEL006126 | 191.997 | 1.229758 | 0.30412 | 4.043663 | 5.26E-05 | 0.028144 |
| AAEL012351 | 162.5252 | 1.471971 | 0.363539 | 4.049009 | 5.14E-05 | 0.028144 |
| AAEL024560 | 144.7041 | 1.161194 | 0.291786 | 3.979602 | 6.90E-05 | 0.035288 |
| AAEL004478 | 15.70645 | 1.311482 | 0.330305 | 3.970516 | 7.17E-05 | 0.035288 |
| AAEL002741 | 36.46224 | 1.158966 | 0.297029 | 3.901856 | 9.55E-05 | 0.043942 |
| AAEL005977 | 533.8883 | 0.761845 | 0.196042 | 3.886133 | 0.000102 | 0.044682 |

FC: fold change; P-adj: adjusted *p* value

**3 dpi 32 ºC downregulated**

| GeneID | Base mean | log2(FC) | StdErr | Wald-Stats | P-value | P-adj |
|---|---|---|---|---|---|---|
| AAEL024207 | 101.1526 | -1.57107 | 0.377418 | -4.16268 | 3.15E-05 | 0.022759 |
| AAEL000668 | 67.81704 | -1.40938 | 0.361439 | -3.89935 | 9.64E-05 | 0.043942 |
| AAEL014019 | 197.0831 | -1.43832 | 0.370897 | -3.87796 | 0.000105 | 0.044682 |
| AAEL011126 | 406.0354 | -1.27551 | 0.331352 | -3.84943 | 0.000118 | 0.048546 |

FC: fold change; P-adj: adjusted *p* value

**7 dpi 18 ºC upregulated**

| GeneID | Base mean | log2(FC) | StdErr | Wald-Stats | P-value | P-adj |
|---|---|---|---|---|---|---|
| AAEL013346 | 420.3165 | 4.176572 | 0.302726 | 13.79656 | 2.67E-43 | 3.46E-39 |
| AAEL013348 | 150.1522 | 3.462339 | 0.280088 | 12.36163 | 4.22E-35 | 2.72E-31 |
| AAEL013350 | 2381.49 | 3.89112 | 0.32236 | 12.07075 | 1.51E-33 | 6.50E-30 |
| AAEL013339 | 63.31234 | 3.71537 | 0.317903 | 11.68712 | 1.48E-31 | 4.79E-28 |
| AAEL013351 | 631.1653 | 3.197844 | 0.296957 | 10.76872 | 4.84E-27 | 1.25E-23 |

| | | | | | | |
|---|---|---|---|---|---|---|
| AAEL017975 | 8101.265 | 3.147815 | 0.327586 | 9.609137 | 7.32E-22 | 1.58E-18 |
| AAEL017976 | 4130.422 | 3.142821 | 0.332892 | 9.440963 | 3.69E-21 | 6.82E-18 |
| AAEL013347 | 800.9459 | 1.953186 | 0.217144 | 8.994871 | 2.37E-19 | 3.82E-16 |
| AAEL022079 | 396.6688 | 2.314959 | 0.279665 | 8.277602 | 1.26E-16 | 1.81E-13 |
| AAEL019751 | 955.3816 | 1.67476 | 0.212444 | 7.883309 | 3.19E-15 | 4.12E-12 |
| AAEL013349 | 274.9653 | 2.490805 | 0.328152 | 7.590404 | 3.19E-14 | 3.75E-11 |
| AAEL006352 | 298.3305 | 1.251859 | 0.172568 | 7.254283 | 4.04E-13 | 4.35E-10 |
| AAEL022253 | 4523.436 | 2.439641 | 0.337721 | 7.223847 | 5.05E-13 | 5.03E-10 |
| AAEL024512 | 162.5634 | 2.388031 | 0.334364 | 7.142006 | 9.20E-13 | 8.49E-10 |
| AAEL022059 | 2395.029 | 2.202666 | 0.33608 | 6.553999 | 5.60E-11 | 4.83E-08 |
| AAEL023321 | 151.8003 | 2.029257 | 0.323287 | 6.276943 | 3.45E-10 | 2.79E-07 |
| AAEL023395 | 121.0724 | 2.045191 | 0.328381 | 6.228096 | 4.72E-10 | 3.59E-07 |
| AAEL027610 | 2684.146 | 2.056135 | 0.337229 | 6.097151 | 1.08E-09 | 7.76E-07 |
| AAEL001857 | 134.3301 | 1.906687 | 0.336916 | 5.659238 | 1.52E-08 | 1.03E-05 |
| AAEL006883 | 927.2206 | 1.564359 | 0.287865 | 5.434357 | 5.50E-08 | 3.55E-05 |
| AAEL003737 | 149.6336 | 1.054883 | 0.197164 | 5.350283 | 8.78E-08 | 5.41E-05 |
| AAEL026833 | 557.9738 | 1.531724 | 0.290752 | 5.26815 | 1.38E-07 | 8.10E-05 |
| AAEL009881 | 354.4552 | 0.934855 | 0.181892 | 5.139604 | 2.75E-07 | 0.000155 |
| AAEL024597 | 267.3648 | 0.828234 | 0.163736 | 5.058341 | 4.23E-07 | 0.000228 |
| AAEL002209 | 189.6914 | 1.11988 | 0.223811 | 5.003687 | 5.62E-07 | 0.000291 |
| AAEL026008 | 1605.445 | 1.410719 | 0.28551 | 4.941043 | 7.77E-07 | 0.000386 |
| AAEL021072 | 40.69005 | 1.404413 | 0.289463 | 4.851787 | 1.22E-06 | 0.000565 |
| AAEL013077 | 1172.399 | 1.259947 | 0.260178 | 4.842638 | 1.28E-06 | 0.000571 |
| AAEL002075 | 33.58957 | 1.425952 | 0.303065 | 4.7051 | 2.54E-06 | 0.001093 |
| AAEL013770 | 92.40521 | 1.069836 | 0.228253 | 4.687066 | 2.77E-06 | 0.001156 |
| AAEL003505 | 2530.577 | 1.361659 | 0.29527 | 4.611572 | 4.00E-06 | 0.001603 |
| AAEL006276 | 238.8906 | 1.189589 | 0.25823 | 4.606706 | 4.09E-06 | 0.001603 |
| AAEL003726 | 44.47114 | 1.516708 | 0.329791 | 4.598992 | 4.25E-06 | 0.00161 |
| AAEL000915 | 310.5454 | 1.320343 | 0.294409 | 4.484733 | 7.30E-06 | 0.002484 |
| AAEL025894 | 23.83281 | 1.477798 | 0.33409 | 4.423353 | 9.72E-06 | 0.003064 |
| AAEL006902 | 726.9372 | 1.383176 | 0.314324 | 4.400477 | 1.08E-05 | 0.003314 |
| AAEL003886 | 54.19795 | 1.174559 | 0.267184 | 4.396067 | 1.10E-05 | 0.003314 |
| AAEL007206 | 983.3226 | 1.088468 | 0.249164 | 4.368476 | 1.25E-05 | 0.003676 |
| AAEL005772 | 485.0575 | 1.387753 | 0.319198 | 4.347618 | 1.38E-05 | 0.003954 |

| | | | | | | |
|---|---|---|---|---|---|---|
| AAEL024038 | 206.2591 | 0.944778 | 0.218621 | 4.321531 | 1.55E-05 | 0.004355 |
| AAEL020092 | 18951.2 | 0.941047 | 0.221991 | 4.239122 | 2.24E-05 | 0.006172 |
| AAEL008622 | 57.86806 | 1.386567 | 0.333706 | 4.155058 | 3.25E-05 | 0.00858 |
| AAEL004919 | 229.0628 | 0.94733 | 0.231526 | 4.091688 | 4.28E-05 | 0.011073 |
| AAEL008953 | 11806.12 | 0.820875 | 0.202235 | 4.059016 | 4.93E-05 | 0.011912 |
| AAEL003728 | 141.0934 | 1.194262 | 0.294594 | 4.053923 | 5.04E-05 | 0.011912 |
| AAEL010262 | 1913.182 | 0.99813 | 0.246301 | 4.052482 | 5.07E-05 | 0.011912 |
| AAEL002661 | 7.098293 | 1.367789 | 0.337293 | 4.055196 | 5.01E-05 | 0.011912 |
| AAEL012352 | 1303.102 | 0.888156 | 0.218583 | 4.063244 | 4.84E-05 | 0.011912 |
| AAEL007200 | 4.079949 | 1.34003 | 0.332504 | 4.030112 | 5.58E-05 | 0.01287 |
| AAEL023591 | 2404.098 | 0.875281 | 0.219422 | 3.989024 | 6.63E-05 | 0.015048 |
| AAEL028247 | 55.4548 | 1.340884 | 0.336799 | 3.981262 | 6.86E-05 | 0.01528 |
| AAEL026031 | 105.5293 | 0.868658 | 0.218818 | 3.969771 | 7.19E-05 | 0.015764 |
| AAEL000334 | 62.87674 | 1.303493 | 0.331749 | 3.929158 | 8.52E-05 | 0.018367 |
| AAEL027802 | 75.8778 | 1.0622 | 0.273047 | 3.890171 | 0.0001 | 0.020888 |
| AAEL024257 | 9.453414 | 1.296162 | 0.337273 | 3.843067 | 0.000122 | 0.023637 |
| AAEL013352 | 267.5548 | 1.291916 | 0.336256 | 3.842063 | 0.000122 | 0.023637 |
| AAEL000445 | 312.4624 | 0.871034 | 0.226984 | 3.837427 | 0.000124 | 0.023637 |
| AAEL013341 | 1175.571 | 0.920469 | 0.241839 | 3.806125 | 0.000141 | 0.025703 |
| AAEL002130 | 3246.238 | 0.881223 | 0.231354 | 3.808978 | 0.00014 | 0.025703 |
| AAEL003883 | 1403.115 | 0.944435 | 0.248609 | 3.798873 | 0.000145 | 0.026099 |
| AAEL024887 | 410.0083 | 0.825397 | 0.218224 | 3.782349 | 0.000155 | 0.026278 |
| AAEL010712 | 1792.319 | 0.910695 | 0.241301 | 3.774108 | 0.000161 | 0.026278 |
| AAEL010120 | 333.82 | 0.855971 | 0.225969 | 3.787994 | 0.000152 | 0.026278 |
| AAEL010242 | 512.101 | 1.274405 | 0.337638 | 3.774468 | 0.00016 | 0.026278 |
| AAEL001928 | 2105.611 | 0.953398 | 0.252352 | 3.778043 | 0.000158 | 0.026278 |
| AAEL000545 | 286.5021 | 1.215807 | 0.320857 | 3.789245 | 0.000151 | 0.026278 |
| AAEL005829 | 244.4502 | 0.81641 | 0.21755 | 3.752741 | 0.000175 | 0.026813 |
| AAEL020330 | 770.7974 | 1.202723 | 0.320253 | 3.755543 | 0.000173 | 0.026813 |
| AAEL025731 | 16.17947 | 1.225227 | 0.325725 | 3.761532 | 0.000169 | 0.026813 |
| AAEL019688 | 34.81125 | 1.268243 | 0.337741 | 3.755078 | 0.000173 | 0.026813 |
| AAEL019704 | 304.8896 | 0.876778 | 0.233212 | 3.759574 | 0.00017 | 0.026813 |
| AAEL000780 | 222.1955 | 0.805576 | 0.214776 | 3.75077 | 0.000176 | 0.026813 |
| AAEL020804 | 278.0058 | 0.618901 | 0.165558 | 3.738272 | 0.000185 | 0.027221 |

| | | | | | | |
|---|---|---|---|---|---|---|
| AAEL007225 | 22.29841 | 1.200419 | 0.320995 | 3.739683 | 0.000184 | 0.027221 |
| AAEL023002 | 15.16646 | 1.262493 | 0.337715 | 3.738335 | 0.000185 | 0.027221 |
| AAEL010260 | 1553.447 | 0.798033 | 0.213993 | 3.729244 | 0.000192 | 0.027898 |
| AAEL019684 | 35.98889 | 1.114147 | 0.299567 | 3.719188 | 0.0002 | 0.028709 |
| AAEL013345 | 672.0795 | 1.143719 | 0.308117 | 3.711969 | 0.000206 | 0.028899 |
| AAEL003966 | 237.7414 | 0.767735 | 0.206747 | 3.713401 | 0.000204 | 0.028899 |
| AAEL003888 | 2273.902 | 1.087065 | 0.294347 | 3.693142 | 0.000222 | 0.030463 |
| AAEL011371 | 3573.692 | 0.750477 | 0.203999 | 3.678828 | 0.000234 | 0.031228 |
| AAEL020028 | 179.8287 | 0.911619 | 0.24755 | 3.682565 | 0.000231 | 0.031228 |
| AAEL005967 | 32.7544 | 1.172761 | 0.319496 | 3.670662 | 0.000242 | 0.031592 |
| AAEL010680 | 65.99109 | 1.230371 | 0.336492 | 3.656461 | 0.000256 | 0.032732 |
| AAEL004023 | 308.9545 | 0.714887 | 0.196669 | 3.634984 | 0.000278 | 0.034489 |
| AAEL002853 | 589.2938 | 0.863334 | 0.237795 | 3.630584 | 0.000283 | 0.034489 |
| AAEL019899 | 7.83657 | 1.223632 | 0.336482 | 3.636547 | 0.000276 | 0.034489 |
| AAEL011520 | 701.7817 | 0.784751 | 0.216038 | 3.632467 | 0.000281 | 0.034489 |
| AAEL023555 | 37.80518 | 1.094452 | 0.302312 | 3.620269 | 0.000294 | 0.035558 |
| AAEL022829 | 12.63532 | 1.18367 | 0.327399 | 3.615373 | 0.0003 | 0.035572 |
| AAEL025332 | 16.18794 | 1.087082 | 0.300661 | 3.615639 | 0.0003 | 0.035572 |
| AAEL029030 | 11.87014 | 1.213811 | 0.336172 | 3.610691 | 0.000305 | 0.035891 |
| AAEL002124 | 879.8008 | 0.680503 | 0.189022 | 3.600127 | 0.000318 | 0.036813 |
| AAEL009541 | 411.2336 | 0.763981 | 0.212251 | 3.599425 | 0.000319 | 0.036813 |
| AAEL013190 | 5.974054 | 1.197853 | 0.334176 | 3.584491 | 0.000338 | 0.038639 |
| AAEL006844 | 18.22367 | 1.172727 | 0.328817 | 3.566498 | 0.000362 | 0.041027 |
| AAEL006132 | 35.57626 | 1.048723 | 0.294406 | 3.562165 | 0.000368 | 0.041348 |
| AAEL010881 | 138.9453 | 1.169439 | 0.329736 | 3.546592 | 0.00039 | 0.043493 |
| AAEL026848 | 42.01983 | 1.190459 | 0.337019 | 3.53232 | 0.000412 | 0.045517 |
| AAEL005961 | 9323.726 | 0.925108 | 0.262746 | 3.520927 | 0.00043 | 0.047115 |

FC: fold change; P-adj: adjusted *p* value

**7 dpi 18 ºC downregulated**

| GeneID | Base mean | log2(FC) | StdErr | Wald-Stats | P-value | P-adj |
|---|---|---|---|---|---|---|
| AAEL023617 | 399.222 | -0.71837 | 0.147716 | -4.86317 | 1.16E-06 | 0.000553 |
| AAEL012219 | 1561.792 | -0.63644 | 0.138551 | -4.59351 | 4.36E-06 | 0.00161 |
| AAEL029039 | 21.97969 | -1.27506 | 0.278798 | -4.57342 | 4.80E-06 | 0.001723 |

| | | | | | | |
|---|---|---|---|---|---|---|
| AAEL006151 | 215.0111 | -1.50159 | 0.337762 | -4.4457 | 8.76E-06 | 0.002856 |
| AAEL009567 | 119.6869 | -1.48522 | 0.33422 | -4.44386 | 8.84E-06 | 0.002856 |
| AAEL008789 | 4835.027 | -1.25408 | 0.2987 | -4.19847 | 2.69E-05 | 0.007238 |
| AAEL010620 | 42.17635 | -1.28654 | 0.330677 | -3.89062 | 1.00E-04 | 0.020888 |
| AAEL011263 | 93.60525 | -0.89784 | 0.23138 | -3.88035 | 0.000104 | 0.021404 |
| AAEL018189 | 56.956 | -1.23673 | 0.320701 | -3.85634 | 0.000115 | 0.02325 |
| AAEL006377 | 320.5194 | -0.92791 | 0.241602 | -3.84068 | 0.000123 | 0.023637 |
| AAEL013432 | 206.882 | -1.04263 | 0.273691 | -3.80951 | 0.000139 | 0.025703 |
| AAEL020035 | 135.6844 | -1.18964 | 0.321919 | -3.69547 | 0.000219 | 0.030463 |
| AAEL025334 | 93.20759 | -1.20526 | 0.327448 | -3.68077 | 0.000233 | 0.031228 |

FC: fold change; P-adj: adjusted *p* value

**7 dpi 28 ºC upregulated**

| GeneID | Base mean | log2(FC) | StdErr | Wald-Stats | P-value | P-adj |
|---|---|---|---|---|---|---|
| AAEL013350 | 728.892 | 3.116381 | 0.297873 | 10.4621 | 1.29E-25 | 1.52E-21 |
| AAEL017975 | 2959.508 | 3.074894 | 0.299122 | 10.27974 | 8.70E-25 | 5.11E-21 |
| AAEL006883 | 667.5833 | 2.292157 | 0.240929 | 9.513818 | 1.84E-21 | 5.40E-18 |
| AAEL023591 | 2032.563 | 1.554986 | 0.163415 | 9.515592 | 1.81E-21 | 5.40E-18 |
| AAEL009645 | 5142.394 | 1.068227 | 0.115341 | 9.261451 | 2.02E-20 | 4.74E-17 |
| AAEL010434 | 178.1841 | 2.635448 | 0.30056 | 8.76845 | 1.81E-18 | 3.55E-15 |
| AAEL020330 | 328.7641 | 2.522 | 0.303941 | 8.29765 | 1.06E-16 | 1.78E-13 |
| AAEL013346 | 171.5745 | 2.440115 | 0.302939 | 8.054812 | 7.96E-16 | 1.17E-12 |
| AAEL025531 | 99.93837 | 2.307221 | 0.292246 | 7.894794 | 2.91E-15 | 3.80E-12 |
| AAEL002655 | 139.2552 | 2.295781 | 0.296012 | 7.755711 | 8.79E-15 | 9.82E-12 |
| AAEL025126 | 151.0833 | 2.294829 | 0.296109 | 7.749954 | 9.19E-15 | 9.82E-12 |
| AAEL022079 | 113.6611 | 1.816592 | 0.235693 | 7.707444 | 1.28E-14 | 1.16E-11 |
| AAEL026008 | 2540.816 | 1.846535 | 0.239494 | 7.710156 | 1.26E-14 | 1.16E-11 |
| AAEL010068 | 1021.116 | 1.601684 | 0.209201 | 7.656194 | 1.92E-14 | 1.61E-11 |
| AAEL017976 | 1384.711 | 2.24417 | 0.303548 | 7.393134 | 1.43E-13 | 1.12E-10 |
| AAEL009387 | 7637.988 | 0.653415 | 0.089925 | 7.266183 | 3.70E-13 | 2.72E-10 |
| AAEL004169 | 367.0314 | 1.321907 | 0.184538 | 7.163348 | 7.87E-13 | 5.44E-10 |
| AAEL007902 | 185.6905 | 1.35207 | 0.189582 | 7.131846 | 9.90E-13 | 6.47E-10 |
| AAEL022253 | 1757.554 | 2.143421 | 0.303135 | 7.07085 | 1.54E-12 | 9.53E-10 |
| AAEL003505 | 2334.232 | 1.719665 | 0.244088 | 7.04527 | 1.85E-12 | 1.09E-09 |

| | | | | | | |
|---|---|---|---|---|---|---|
| AAEL017380 | 93.3928 | 2.127009 | 0.302186 | 7.038741 | 1.94E-12 | 1.09E-09 |
| AAEL021302 | 473.302 | 1.652228 | 0.235395 | 7.01897 | 2.24E-12 | 1.19E-09 |
| AAEL009487 | 3705.967 | 0.796327 | 0.113736 | 7.001517 | 2.53E-12 | 1.29E-09 |
| AAEL008622 | 67.37195 | 1.985214 | 0.289337 | 6.861246 | 6.83E-12 | 3.34E-09 |
| AAEL011371 | 4347.899 | 0.98818 | 0.147226 | 6.711997 | 1.92E-11 | 9.03E-09 |
| AAEL003728 | 250.8779 | 1.682361 | 0.255804 | 6.576753 | 4.81E-11 | 2.10E-08 |
| AAEL003345 | 5246.289 | 1.764473 | 0.268303 | 6.57643 | 4.82E-11 | 2.10E-08 |
| AAEL007126 | 122.7807 | 1.415538 | 0.216427 | 6.540483 | 6.13E-11 | 2.49E-08 |
| AAEL010769 | 738.499 | 1.296685 | 0.198184 | 6.54284 | 6.04E-11 | 2.49E-08 |
| AAEL013348 | 58.85481 | 1.851582 | 0.287246 | 6.445975 | 1.15E-10 | 4.50E-08 |
| AAEL006126 | 71.78147 | 1.881567 | 0.296864 | 6.338145 | 2.33E-10 | 8.82E-08 |
| AAEL024838 | 419.5059 | 1.126007 | 0.178562 | 6.305962 | 2.86E-10 | 1.05E-07 |
| AAEL010379 | 398.7471 | 1.350192 | 0.214908 | 6.28266 | 3.33E-10 | 1.19E-07 |
| AAEL004688 | 2124.404 | 0.60024 | 0.096003 | 6.252293 | 4.04E-10 | 1.40E-07 |
| AAEL006904 | 1299.323 | 0.695429 | 0.11258 | 6.177196 | 6.53E-10 | 2.19E-07 |
| AAEL009171 | 1750.662 | 0.833242 | 0.138185 | 6.029908 | 1.64E-09 | 5.36E-07 |
| AAEL019463 | 327.5993 | 0.914593 | 0.15194 | 6.01944 | 1.75E-09 | 5.56E-07 |
| AAEL001857 | 81.81225 | 1.553 | 0.262458 | 5.917143 | 3.28E-09 | 9.87E-07 |
| AAEL008635 | 788.3385 | 1.493264 | 0.255311 | 5.848804 | 4.95E-09 | 1.46E-06 |
| AAEL002610 | 1599.883 | 1.588379 | 0.273095 | 5.816219 | 6.02E-09 | 1.73E-06 |
| AAEL021929 | 320.0132 | 1.748408 | 0.303424 | 5.762268 | 8.30E-09 | 2.32E-06 |
| AAEL015631 | 557.4039 | 1.066857 | 0.186659 | 5.715537 | 1.09E-08 | 2.99E-06 |
| AAEL013857 | 245.9264 | 1.518453 | 0.26597 | 5.709103 | 1.14E-08 | 3.03E-06 |
| AAEL001420 | 5636.799 | 1.109987 | 0.195588 | 5.675132 | 1.39E-08 | 3.62E-06 |
| AAEL026603 | 38.27262 | 1.597974 | 0.284453 | 5.617714 | 1.93E-08 | 4.94E-06 |
| AAEL008953 | 8775.947 | 0.709779 | 0.126438 | 5.613638 | 1.98E-08 | 4.96E-06 |
| AAEL009126 | 64.96642 | 1.676161 | 0.301159 | 5.565701 | 2.61E-08 | 6.39E-06 |
| AAEL019751 | 508.2229 | 1.160973 | 0.209594 | 5.53914 | 3.04E-08 | 7.29E-06 |
| AAEL026833 | 890.4588 | 1.575777 | 0.284982 | 5.529398 | 3.21E-08 | 7.55E-06 |
| AAEL023321 | 107.0405 | 1.434187 | 0.260189 | 5.512088 | 3.55E-08 | 8.02E-06 |
| AAEL001929 | 32.82776 | 1.346273 | 0.244091 | 5.515464 | 3.48E-08 | 8.02E-06 |
| AAEL004803 | 107.9252 | 0.861928 | 0.156547 | 5.505866 | 3.67E-08 | 8.15E-06 |
| AAEL015465 | 208.0624 | 1.260426 | 0.229887 | 5.482799 | 4.19E-08 | 9.11E-06 |
| AAEL012856 | 434.0041 | 1.590409 | 0.290414 | 5.476358 | 4.34E-08 | 9.28E-06 |

| | | | | | | |
|---|---|---|---|---|---|---|
| AAEL027610 | 762.9641 | 1.629492 | 0.298347 | 5.461729 | 4.72E-08 | 9.90E-06 |
| AAEL005768 | 252.2436 | 1.283613 | 0.235575 | 5.448853 | 5.07E-08 | 1.05E-05 |
| AAEL009762 | 149.2643 | 1.509447 | 0.278382 | 5.422207 | 5.89E-08 | 1.19E-05 |
| AAEL002969 | 586.2887 | 1.410403 | 0.263625 | 5.35004 | 8.79E-08 | 1.75E-05 |
| AAEL006138 | 126.1631 | 1.598621 | 0.299556 | 5.336639 | 9.47E-08 | 1.86E-05 |
| AAEL001414 | 3151.493 | 1.219908 | 0.232022 | 5.25773 | 1.46E-07 | 2.77E-05 |
| AAEL004048 | 281.9358 | 1.006605 | 0.192029 | 5.241931 | 1.59E-07 | 2.96E-05 |
| AAEL007197 | 782.7576 | 0.689791 | 0.132277 | 5.214737 | 1.84E-07 | 3.28E-05 |
| AAEL002499 | 573.4986 | 0.883405 | 0.170364 | 5.185385 | 2.16E-07 | 3.78E-05 |
| AAEL012089 | 770.8618 | 0.992716 | 0.193761 | 5.123406 | 3.00E-07 | 5.19E-05 |
| AAEL008887 | 522.4587 | 1.43256 | 0.280674 | 5.103993 | 3.33E-07 | 5.67E-05 |
| AAEL009055 | 279.3157 | 1.151009 | 0.227245 | 5.065057 | 4.08E-07 | 6.86E-05 |
| AAEL007969 | 72.9558 | 1.186819 | 0.235566 | 5.038153 | 4.70E-07 | 7.75E-05 |
| AAEL017345 | 1159.848 | 1.19892 | 0.238336 | 5.030378 | 4.90E-07 | 7.88E-05 |
| AAEL008050 | 52.41532 | 1.521366 | 0.30315 | 5.018518 | 5.21E-07 | 8.27E-05 |
| AAEL005992 | 109.7268 | 1.470094 | 0.29646 | 4.958831 | 7.09E-07 | 0.000107 |
| AAEL006543 | 353.2948 | 0.734391 | 0.14816 | 4.956759 | 7.17E-07 | 0.000107 |
| AAEL015298 | 1103.794 | 0.754626 | 0.152215 | 4.957644 | 7.14E-07 | 0.000107 |
| AAEL011038 | 519.7914 | 1.035706 | 0.209131 | 4.95242 | 7.33E-07 | 0.000108 |
| AAEL003888 | 1267.242 | 1.231022 | 0.249439 | 4.935158 | 8.01E-07 | 0.000116 |
| AAEL021555 | 163.3378 | 0.930546 | 0.190445 | 4.886154 | 1.03E-06 | 0.000147 |
| AAEL014138 | 109.9276 | 0.956871 | 0.195996 | 4.882103 | 1.05E-06 | 0.000149 |
| AAEL003597 | 194.5737 | 0.994111 | 0.203906 | 4.875352 | 1.09E-06 | 0.000152 |
| AAEL006269 | 221.219 | 0.930504 | 0.19112 | 4.868691 | 1.12E-06 | 0.000155 |
| AAEL013840 | 114.4479 | 0.757733 | 0.156048 | 4.855775 | 1.20E-06 | 0.000164 |
| AAEL008473 | 1848.89 | 1.401784 | 0.288906 | 4.852041 | 1.22E-06 | 0.000165 |
| AAEL018343 | 221.6811 | 0.975159 | 0.201241 | 4.845725 | 1.26E-06 | 0.000169 |
| AAEL013984 | 1037.616 | 1.090209 | 0.225189 | 4.841296 | 1.29E-06 | 0.00017 |
| AAEL018340 | 214.9941 | 0.958824 | 0.198471 | 4.831046 | 1.36E-06 | 0.000174 |
| AAEL014618 | 40.02552 | 1.446133 | 0.300669 | 4.809722 | 1.51E-06 | 0.000191 |
| AAEL011453 | 102.1792 | 1.238665 | 0.258308 | 4.795309 | 1.62E-06 | 0.000203 |
| AAEL021099 | 252.744 | 1.132791 | 0.237184 | 4.77601 | 1.79E-06 | 0.000214 |
| AAEL008471 | 396.7432 | 0.640615 | 0.134893 | 4.749073 | 2.04E-06 | 0.000243 |
| AAEL005763 | 427.0991 | 0.75767 | 0.160337 | 4.725491 | 2.30E-06 | 0.00027 |

| | | | | | | |
|---|---|---|---|---|---|---|
| AAEL001794 | 1045.009 | 1.133738 | 0.241373 | 4.697033 | 2.64E-06 | 0.000301 |
| AAEL006815 | 180.6059 | 0.928441 | 0.197772 | 4.69449 | 2.67E-06 | 0.000302 |
| AAEL010102 | 242.2174 | 0.684095 | 0.145861 | 4.690049 | 2.73E-06 | 0.000303 |
| AAEL007773 | 256.9152 | 0.780883 | 0.166465 | 4.690962 | 2.72E-06 | 0.000303 |
| AAEL013770 | 53.71287 | 1.269363 | 0.272387 | 4.660153 | 3.16E-06 | 0.000347 |
| AAEL008106 | 809.437 | 1.12908 | 0.242515 | 4.655707 | 3.23E-06 | 0.000351 |
| AAEL022059 | 926.7941 | 1.340581 | 0.28813 | 4.65269 | 3.28E-06 | 0.000353 |
| AAEL003051 | 58.52314 | 0.828919 | 0.180728 | 4.586556 | 4.51E-06 | 0.000469 |
| AAEL003527 | 40.58973 | 0.877781 | 0.192604 | 4.557451 | 5.18E-06 | 0.00053 |
| AAEL001965 | 752.0782 | 1.124173 | 0.246693 | 4.556964 | 5.19E-06 | 0.00053 |
| AAEL002124 | 737.2991 | 0.996084 | 0.218918 | 4.550032 | 5.36E-06 | 0.000539 |
| AAEL006990 | 541.2391 | 1.19069 | 0.262658 | 4.533239 | 5.81E-06 | 0.000579 |
| AAEL024512 | 169.479 | 1.362707 | 0.30104 | 4.526671 | 5.99E-06 | 0.000587 |
| AAEL003640 | 290.3398 | 0.783619 | 0.173109 | 4.526736 | 5.99E-06 | 0.000587 |
| AAEL013345 | 270.9954 | 1.298092 | 0.289007 | 4.491561 | 7.07E-06 | 0.000687 |
| AAEL024583 | 1618.893 | 0.784361 | 0.175192 | 4.477154 | 7.56E-06 | 0.000729 |
| AAEL025903 | 169.268 | 1.038508 | 0.235003 | 4.419121 | 9.91E-06 | 0.000932 |
| AAEL026175 | 1736.262 | 1.062213 | 0.240835 | 4.410539 | 1.03E-05 | 0.000962 |
| AAEL001575 | 27.2747 | 0.993797 | 0.22556 | 4.405901 | 1.05E-05 | 0.000975 |
| AAEL009863 | 1853.093 | 0.697844 | 0.159377 | 4.378583 | 1.19E-05 | 0.001097 |
| AAEL008495 | 295.8199 | 0.829751 | 0.190175 | 4.363096 | 1.28E-05 | 0.001161 |
| AAEL026300 | 26.30127 | 1.25414 | 0.287464 | 4.362779 | 1.28E-05 | 0.001161 |
| AAEL019537 | 189.0761 | 1.020824 | 0.234192 | 4.358914 | 1.31E-05 | 0.001173 |
| AAEL008963 | 629.9184 | 0.84189 | 0.193384 | 4.353455 | 1.34E-05 | 0.001193 |
| AAEL010206 | 334.0111 | 1.097843 | 0.252299 | 4.351361 | 1.35E-05 | 0.001196 |
| AAEL007092 | 1327.612 | 0.596863 | 0.137465 | 4.341927 | 1.41E-05 | 0.00123 |
| AAEL005533 | 51.15832 | 1.313907 | 0.303443 | 4.329991 | 1.49E-05 | 0.001286 |
| AAEL000304 | 707.0519 | 0.655257 | 0.15137 | 4.328856 | 1.50E-05 | 0.001286 |
| AAEL012712 | 187.5658 | 1.231312 | 0.284744 | 4.324269 | 1.53E-05 | 0.001288 |
| AAEL014566 | 133.3158 | 0.980285 | 0.227344 | 4.311902 | 1.62E-05 | 0.001331 |
| AAEL002600 | 1788.124 | 0.798765 | 0.185465 | 4.306818 | 1.66E-05 | 0.001343 |
| AAEL013554 | 18.85148 | 1.134401 | 0.264411 | 4.290301 | 1.78E-05 | 0.001417 |
| AAEL002049 | 670.6905 | 0.899753 | 0.209911 | 4.286343 | 1.82E-05 | 0.001433 |
| AAEL006576 | 1406.726 | 1.045628 | 0.24444 | 4.277641 | 1.89E-05 | 0.00147 |

| | | | | | | |
|---|---|---|---|---|---|---|
| AAEL008625 | 52.39674 | 0.851006 | 0.198909 | 4.278372 | 1.88E-05 | 0.00147 |
| AAEL001232 | 297.9011 | 0.685302 | 0.160411 | 4.272175 | 1.94E-05 | 0.001487 |
| AAEL007128 | 277.9654 | 1.032843 | 0.24215 | 4.265311 | 2.00E-05 | 0.001519 |
| AAEL024291 | 93.41137 | 0.695878 | 0.163489 | 4.256417 | 2.08E-05 | 0.001545 |
| AAEL002959 | 109.1231 | 1.199275 | 0.282411 | 4.246553 | 2.17E-05 | 0.001585 |
| AAEL003426 | 36.92346 | 0.998294 | 0.235068 | 4.246838 | 2.17E-05 | 0.001585 |
| AAEL019658 | 968.931 | 0.624681 | 0.148328 | 4.211477 | 2.54E-05 | 0.001812 |
| AAEL023524 | 233.7648 | 0.843538 | 0.20032 | 4.210947 | 2.54E-05 | 0.001812 |
| AAEL017023 | 718.0172 | 0.98708 | 0.2347 | 4.205699 | 2.60E-05 | 0.001831 |
| AAEL011810 | 461.4027 | 0.89686 | 0.213235 | 4.205963 | 2.60E-05 | 0.001831 |
| AAEL027700 | 141.6591 | 1.000672 | 0.238004 | 4.204441 | 2.62E-05 | 0.001831 |
| AAEL008028 | 607.322 | 0.78128 | 0.185952 | 4.201517 | 2.65E-05 | 0.001841 |
| AAEL010151 | 275.9798 | 0.702215 | 0.167171 | 4.20059 | 2.66E-05 | 0.001841 |
| AAEL026843 | 5521.472 | 0.957165 | 0.229077 | 4.178348 | 2.94E-05 | 0.002001 |
| AAEL013163 | 811.1976 | 0.797777 | 0.191223 | 4.171964 | 3.02E-05 | 0.00204 |
| AAEL013339 | 16.58957 | 1.210306 | 0.29106 | 4.15827 | 3.21E-05 | 0.002142 |
| AAEL018334 | 329.0879 | 0.681031 | 0.164547 | 4.138836 | 3.49E-05 | 0.002318 |
| AAEL002467 | 1059.453 | 1.249607 | 0.302181 | 4.135296 | 3.54E-05 | 0.002341 |
| AAEL005660 | 209.4714 | 0.626625 | 0.151797 | 4.128043 | 3.66E-05 | 0.002389 |
| AAEL001307 | 44.62804 | 0.890785 | 0.216366 | 4.117035 | 3.84E-05 | 0.002492 |
| AAEL006794 | 3469.437 | 0.666262 | 0.162045 | 4.111581 | 3.93E-05 | 0.002538 |
| AAEL008190 | 277.0882 | 0.807751 | 0.197677 | 4.086228 | 4.38E-05 | 0.00275 |
| AAEL007619 | 401.9259 | 0.896309 | 0.220135 | 4.07163 | 4.67E-05 | 0.002888 |
| AAEL021375 | 213.6006 | 1.039871 | 0.255942 | 4.062915 | 4.85E-05 | 0.002952 |
| AAEL001646 | 64.59255 | 1.091138 | 0.2685 | 4.063831 | 4.83E-05 | 0.002952 |
| AAEL008144 | 2003.838 | 0.73361 | 0.180981 | 4.053512 | 5.05E-05 | 0.003057 |
| AAEL019564 | 326.2307 | 0.896106 | 0.221539 | 4.044918 | 5.23E-05 | 0.003139 |
| AAEL012981 | 262.2759 | 0.626864 | 0.155021 | 4.043747 | 5.26E-05 | 0.003139 |
| AAEL014348 | 118.0114 | 0.813352 | 0.201371 | 4.039074 | 5.37E-05 | 0.003186 |
| AAEL021263 | 89.96674 | 1.04891 | 0.260213 | 4.030975 | 5.55E-05 | 0.003265 |
| AAEL022932 | 62.93918 | 1.037877 | 0.258007 | 4.022672 | 5.75E-05 | 0.00335 |
| AAEL028635 | 201.4996 | 1.205347 | 0.299882 | 4.019398 | 5.83E-05 | 0.003379 |
| AAEL003726 | 32.35003 | 1.206955 | 0.30326 | 3.979935 | 6.89E-05 | 0.003915 |
| AAEL022674 | 62.18833 | 1.134412 | 0.285747 | 3.96999 | 7.19E-05 | 0.004062 |

| | | | | | | |
|---|---|---|---|---|---|---|
| AAEL008157 | 110.7213 | 0.937194 | 0.23688 | 3.956402 | 7.61E-05 | 0.004239 |
| AAEL003139 | 113.5658 | 0.738432 | 0.187222 | 3.94415 | 8.01E-05 | 0.00444 |
| AAEL003844 | 1921.863 | 0.607159 | 0.154337 | 3.933972 | 8.36E-05 | 0.004611 |
| AAEL009213 | 373.5242 | 0.792267 | 0.201455 | 3.932736 | 8.40E-05 | 0.004613 |
| AAEL000049 | 182.7444 | 0.71724 | 0.182497 | 3.930155 | 8.49E-05 | 0.004641 |
| AAEL000037 | 254.1255 | 0.967964 | 0.246773 | 3.922491 | 8.76E-05 | 0.004769 |
| AAEL005017 | 892.4331 | 0.616856 | 0.15761 | 3.913805 | 9.09E-05 | 0.00488 |
| AAEL027188 | 30.93828 | 1.128159 | 0.288263 | 3.913649 | 9.09E-05 | 0.00488 |
| AAEL007444 | 5000.773 | 0.617793 | 0.158059 | 3.908614 | 9.28E-05 | 0.00496 |
| AAEL011452 | 136.996 | 0.865411 | 0.221666 | 3.904129 | 9.46E-05 | 0.00503 |
| AAEL006069 | 299.3025 | 0.68483 | 0.175563 | 3.900773 | 9.59E-05 | 0.005077 |
| AAEL024337 | 420.0972 | 0.656047 | 0.168875 | 3.884807 | 0.000102 | 0.005342 |
| AAEL006535 | 160.339 | 0.974352 | 0.250856 | 3.884107 | 0.000103 | 0.005342 |
| AAEL003632 | 26.05914 | 1.170244 | 0.301375 | 3.883012 | 0.000103 | 0.005343 |
| AAEL002741 | 37.61778 | 0.861112 | 0.222433 | 3.871337 | 0.000108 | 0.005581 |
| AAEL003626 | 245.7445 | 0.931098 | 0.240752 | 3.867457 | 0.00011 | 0.005621 |
| AAEL007191 | 182.0107 | 0.618911 | 0.160236 | 3.862503 | 0.000112 | 0.005662 |
| AAEL002683 | 650.203 | 0.619203 | 0.160689 | 3.853431 | 0.000116 | 0.005851 |
| AAEL001960 | 341.1649 | 0.795728 | 0.207012 | 3.843878 | 0.000121 | 0.005963 |
| AAEL001417 | 154.6086 | 1.080971 | 0.281668 | 3.83775 | 0.000124 | 0.006074 |
| AAEL014605 | 1723.851 | 0.905679 | 0.236037 | 3.837017 | 0.000125 | 0.006074 |
| AAEL001864 | 4113.008 | 0.986691 | 0.257224 | 3.835927 | 0.000125 | 0.006076 |
| AAEL028247 | 20.69271 | 1.164048 | 0.30383 | 3.831245 | 0.000127 | 0.006168 |
| AAEL017334 | 2081.961 | 1.094645 | 0.285838 | 3.829592 | 0.000128 | 0.006184 |
| AAEL008365 | 401.2089 | 1.002519 | 0.262019 | 3.826133 | 0.00013 | 0.00622 |
| AAEL007778 | 4061.046 | 0.834316 | 0.21842 | 3.819778 | 0.000134 | 0.006303 |
| AAEL013347 | 602.0945 | 1.011676 | 0.265541 | 3.809868 | 0.000139 | 0.00646 |
| AAEL010688 | 103.3768 | 0.757898 | 0.19934 | 3.802038 | 0.000144 | 0.006616 |
| AAEL023844 | 216.4577 | 1.008718 | 0.265749 | 3.795751 | 0.000147 | 0.006707 |
| AAEL017144 | 1478.453 | 1.0164 | 0.267694 | 3.796878 | 0.000147 | 0.006707 |
| AAEL010884 | 2076.732 | 1.075631 | 0.283806 | 3.790025 | 0.000151 | 0.006837 |
| AAEL017139 | 58.74374 | 0.971667 | 0.258125 | 3.764321 | 0.000167 | 0.007493 |
| AAEL026751 | 4615.016 | 0.765989 | 0.203747 | 3.759505 | 0.00017 | 0.007608 |
| AAEL020502 | 214.3529 | 0.791625 | 0.210813 | 3.755111 | 0.000173 | 0.007686 |

| | | | | | | |
|---|---|---|---|---|---|---|
| AAEL014391 | 111.498 | 0.851273 | 0.227605 | 3.740132 | 0.000184 | 0.008079 |
| AAEL012853 | 159.8212 | 0.851246 | 0.227676 | 3.738844 | 0.000185 | 0.008079 |
| AAEL022775 | 49.32167 | 1.080662 | 0.29047 | 3.720391 | 0.000199 | 0.008628 |
| AAEL002385 | 297.9651 | 0.681258 | 0.183269 | 3.717264 | 0.000201 | 0.008672 |
| AAEL020392 | 13.62418 | 1.128363 | 0.303514 | 3.717661 | 0.000201 | 0.008672 |
| AAEL002893 | 24.91057 | 1.043772 | 0.28233 | 3.696988 | 0.000218 | 0.009225 |
| AAEL002390 | 743.3171 | 0.706725 | 0.191132 | 3.697575 | 0.000218 | 0.009225 |
| AAEL027984 | 23.0876 | 0.974827 | 0.264064 | 3.691631 | 0.000223 | 0.009369 |
| AAEL010596 | 104.2237 | 0.869562 | 0.235581 | 3.691133 | 0.000223 | 0.009369 |
| AAEL017513 | 366.3585 | 0.98735 | 0.267601 | 3.689633 | 0.000225 | 0.009369 |
| AAEL013732 | 72.55543 | 0.959131 | 0.260256 | 3.685339 | 0.000228 | 0.009412 |
| AAEL027545 | 21.0711 | 0.871389 | 0.236605 | 3.682891 | 0.000231 | 0.009412 |
| AAEL006805 | 1232.235 | 0.846908 | 0.229933 | 3.683282 | 0.00023 | 0.009412 |
| AAEL003737 | 145.1071 | 0.698272 | 0.190192 | 3.671412 | 0.000241 | 0.009767 |
| AAEL012702 | 2185.461 | 0.949385 | 0.259582 | 3.657367 | 0.000255 | 0.010223 |
| AAEL005731 | 1630.859 | 0.759913 | 0.207874 | 3.655648 | 0.000257 | 0.010257 |
| AAEL011365 | 87.11608 | 0.63695 | 0.174387 | 3.652504 | 0.00026 | 0.010348 |
| AAEL025921 | 1053.944 | 0.622837 | 0.170665 | 3.649471 | 0.000263 | 0.010436 |
| AAEL013853 | 17.96277 | 0.924485 | 0.253415 | 3.648101 | 0.000264 | 0.010456 |
| AAEL002412 | 789.0255 | 0.700523 | 0.192256 | 3.6437 | 0.000269 | 0.010587 |
| AAEL010269 | 161.0076 | 0.789138 | 0.216695 | 3.641693 | 0.000271 | 0.010613 |
| AAEL010180 | 593.7807 | 1.097152 | 0.301525 | 3.638676 | 0.000274 | 0.010667 |
| AAEL009629 | 2065.598 | 0.657224 | 0.180667 | 3.63777 | 0.000275 | 0.010669 |
| AAEL000834 | 472.9553 | 0.906143 | 0.249399 | 3.633301 | 0.00028 | 0.01082 |
| AAEL029061 | 63.28002 | 0.897938 | 0.247607 | 3.62647 | 0.000287 | 0.011037 |
| AAEL006563 | 32.15283 | 0.977453 | 0.270784 | 3.609721 | 0.000307 | 0.011586 |
| AAEL009955 | 15274.56 | 0.903363 | 0.250374 | 3.608057 | 0.000308 | 0.011623 |
| AAEL010773 | 257.6855 | 0.90279 | 0.250571 | 3.602929 | 0.000315 | 0.011817 |
| AAEL012395 | 319.2499 | 0.933424 | 0.25932 | 3.599504 | 0.000319 | 0.011919 |
| AAEL005738 | 133.9378 | 0.856895 | 0.238204 | 3.597308 | 0.000322 | 0.011919 |
| AAEL007342 | 1814.734 | 0.735001 | 0.204246 | 3.59861 | 0.00032 | 0.011919 |
| AAEL005921 | 200.5262 | 0.632953 | 0.175988 | 3.596577 | 0.000322 | 0.011919 |
| AAEL007808 | 124.4748 | 0.747978 | 0.207877 | 3.598185 | 0.00032 | 0.011919 |
| AAEL013351 | 301.1988 | 1.029996 | 0.28754 | 3.582098 | 0.000341 | 0.012482 |

| | | | | | | |
|---|---|---|---|---|---|---|
| AAEL006179 | 57.31212 | 1.082713 | 0.302352 | 3.580964 | 0.000342 | 0.012497 |
| AAEL012513 | 352.2005 | 0.593043 | 0.165671 | 3.57964 | 0.000344 | 0.012522 |
| AAEL021257 | 60.41726 | 1.000669 | 0.279999 | 3.57383 | 0.000352 | 0.012646 |
| AAEL023294 | 4593.122 | 0.663776 | 0.186367 | 3.561659 | 0.000369 | 0.013207 |
| AAEL001189 | 8.51749 | 1.045852 | 0.294116 | 3.555914 | 0.000377 | 0.013336 |
| AAEL002254 | 513.1662 | 0.670733 | 0.188927 | 3.550224 | 0.000385 | 0.013529 |
| AAEL005386 | 42.92092 | 0.658306 | 0.185548 | 3.547905 | 0.000388 | 0.013545 |
| AAEL007878 | 65.59102 | 0.89719 | 0.252866 | 3.548086 | 0.000388 | 0.013545 |
| AAEL020936 | 343.6 | 0.582133 | 0.164269 | 3.543785 | 0.000394 | 0.013717 |
| AAEL014354 | 131.8967 | 0.861564 | 0.243493 | 3.538354 | 0.000403 | 0.013961 |
| AAEL023125 | 173.2256 | 0.847134 | 0.239761 | 3.533246 | 0.00041 | 0.014192 |
| AAEL015430 | 98.33715 | 0.812388 | 0.230083 | 3.530845 | 0.000414 | 0.014269 |
| AAEL001582 | 257.3216 | 0.721753 | 0.204867 | 3.523028 | 0.000427 | 0.014495 |
| AAEL002130 | 2551.637 | 0.667593 | 0.189437 | 3.524081 | 0.000425 | 0.014495 |
| AAEL007856 | 23.18957 | 1.069977 | 0.303915 | 3.520648 | 0.00043 | 0.014583 |
| AAEL008828 | 126.6542 | 0.869882 | 0.248012 | 3.507423 | 0.000452 | 0.015107 |
| AAEL003316 | 175.307 | 0.660549 | 0.18852 | 3.503866 | 0.000459 | 0.015213 |
| AAEL013525 | 405.2231 | 0.63866 | 0.182583 | 3.497919 | 0.000469 | 0.01544 |
| AAEL021073 | 843.091 | 0.77509 | 0.221849 | 3.493771 | 0.000476 | 0.015594 |
| AAEL026446 | 374.8875 | 0.842943 | 0.241556 | 3.489634 | 0.000484 | 0.01575 |
| AAEL006434 | 208.574 | 0.625826 | 0.179421 | 3.48803 | 0.000487 | 0.015801 |
| AAEL002624 | 261.4895 | 0.954202 | 0.273828 | 3.484679 | 0.000493 | 0.015956 |
| AAEL007624 | 598.2947 | 0.721101 | 0.207027 | 3.483124 | 0.000496 | 0.016005 |
| AAEL008007 | 415.0432 | 1.054705 | 0.303474 | 3.475443 | 0.00051 | 0.016258 |
| AAEL005428 | 3100.073 | 0.935934 | 0.269743 | 3.469732 | 0.000521 | 0.016503 |
| AAEL021795 | 2528.497 | 0.933923 | 0.269263 | 3.468443 | 0.000523 | 0.016503 |
| AAEL004833 | 57.25577 | 0.988494 | 0.285181 | 3.466203 | 0.000528 | 0.016503 |
| AAEL007296 | 445.058 | 0.930626 | 0.26835 | 3.467957 | 0.000524 | 0.016503 |
| AAEL024406 | 1464.12 | 0.839349 | 0.242338 | 3.463546 | 0.000533 | 0.016577 |
| AAEL006323 | 218.9261 | 1.029202 | 0.297273 | 3.462147 | 0.000536 | 0.016577 |
| AAEL013027 | 5.590501 | 0.974992 | 0.282532 | 3.450913 | 0.000559 | 0.017063 |
| AAEL024540 | 136.8602 | 0.788553 | 0.228667 | 3.448484 | 0.000564 | 0.017077 |
| AAEL004213 | 29.94395 | 0.939 | 0.272346 | 3.447821 | 0.000565 | 0.017077 |
| AAEL022982 | 513.8749 | 0.704866 | 0.204489 | 3.446954 | 0.000567 | 0.017088 |

| | | | | | | |
|---|---|---|---|---|---|---|
| AAEL005959 | 26.68813 | 1.008493 | 0.29272 | 3.445255 | 0.000571 | 0.017108 |
| AAEL022829 | 5.715702 | 1.042455 | 0.302657 | 3.444348 | 0.000572 | 0.017122 |
| AAEL019487 | 1582.578 | 0.625554 | 0.181687 | 3.443021 | 0.000575 | 0.017163 |
| AAEL011203 | 61.36906 | 0.941392 | 0.273753 | 3.438838 | 0.000584 | 0.017383 |
| AAEL014255 | 83.20852 | 0.892642 | 0.259624 | 3.438207 | 0.000586 | 0.017383 |
| AAEL024269 | 23.30053 | 0.959329 | 0.279297 | 3.434806 | 0.000593 | 0.017558 |
| AAEL023847 | 24.28288 | 0.975997 | 0.284341 | 3.432486 | 0.000598 | 0.017664 |
| AAEL006723 | 228.1399 | 0.752672 | 0.219584 | 3.427714 | 0.000609 | 0.017901 |
| AAEL013532 | 227.3889 | 0.594484 | 0.174487 | 3.407036 | 0.000657 | 0.018968 |
| AAEL006795 | 153.574 | 0.867536 | 0.255318 | 3.397866 | 0.000679 | 0.01933 |
| AAEL019954 | 52.71657 | 1.023466 | 0.301517 | 3.39439 | 0.000688 | 0.01953 |
| AAEL015644 | 201.0627 | 0.77547 | 0.22864 | 3.391657 | 0.000695 | 0.019678 |
| AAEL000243 | 37.70593 | 1.020635 | 0.301085 | 3.389863 | 0.000699 | 0.01976 |
| AAEL021016 | 159.3543 | 0.62619 | 0.18479 | 3.388663 | 0.000702 | 0.019799 |
| AAEL003645 | 1160.819 | 0.82356 | 0.243852 | 3.377298 | 0.000732 | 0.020247 |
| AAEL007090 | 140.8638 | 0.932203 | 0.275979 | 3.377807 | 0.000731 | 0.020247 |
| AAEL013225 | 1609.375 | 0.658488 | 0.194959 | 3.377564 | 0.000731 | 0.020247 |
| AAEL017403 | 12.0762 | 0.899032 | 0.266321 | 3.375748 | 0.000736 | 0.020266 |
| AAEL000811 | 265.837 | 0.869935 | 0.257853 | 3.373758 | 0.000741 | 0.020365 |
| AAEL008961 | 149.261 | 0.626215 | 0.186067 | 3.365535 | 0.000764 | 0.020884 |
| AAEL024753 | 88.15822 | 0.931237 | 0.276856 | 3.363614 | 0.000769 | 0.020933 |
| AAEL013613 | 2085.21 | 0.701672 | 0.208757 | 3.361196 | 0.000776 | 0.021068 |
| AAEL023100 | 28.37116 | 0.794484 | 0.236456 | 3.359968 | 0.00078 | 0.021113 |
| AAEL014350 | 60.52949 | 0.992138 | 0.295426 | 3.358326 | 0.000784 | 0.021118 |
| AAEL013257 | 40.17639 | 0.589155 | 0.175389 | 3.35913 | 0.000782 | 0.021118 |
| AAEL007926 | 321.0101 | 0.935373 | 0.279236 | 3.349759 | 0.000809 | 0.021658 |
| AAEL001586 | 273.0323 | 0.640416 | 0.191583 | 3.342756 | 0.00083 | 0.022066 |
| AAEL012726 | 109.2739 | 0.638451 | 0.191096 | 3.340986 | 0.000835 | 0.02212 |
| AAEL007794 | 40.43803 | 0.905844 | 0.27115 | 3.340753 | 0.000836 | 0.02212 |
| AAEL014045 | 182.8779 | 0.798563 | 0.239355 | 3.336312 | 0.000849 | 0.022426 |
| AAEL009936 | 29.19303 | 0.850153 | 0.255156 | 3.331887 | 0.000863 | 0.022583 |
| AAEL002765 | 305.1896 | 0.610712 | 0.183396 | 3.330018 | 0.000868 | 0.022634 |
| AAEL013434 | 240.1297 | 0.588193 | 0.178012 | 3.304225 | 0.000952 | 0.024338 |
| AAEL006585 | 279.2277 | 0.834265 | 0.252665 | 3.301864 | 0.00096 | 0.024358 |

| | | | | | | |
|---|---|---|---|---|---|---|
| AAEL013341 | 791.4733 | 0.745544 | 0.225753 | 3.302475 | 0.000958 | 0.024358 |
| AAEL009695 | 105.5119 | 0.754852 | 0.228634 | 3.301569 | 0.000961 | 0.024358 |
| AAEL009952 | 99.87689 | 0.798505 | 0.242279 | 3.295807 | 0.000981 | 0.024809 |
| AAEL007907 | 116.1941 | 0.804487 | 0.244428 | 3.291313 | 0.000997 | 0.025047 |
| AAEL004701 | 126.522 | 0.955274 | 0.290841 | 3.284526 | 0.001022 | 0.025573 |
| AAEL019847 | 5.289382 | 0.994898 | 0.302984 | 3.283663 | 0.001025 | 0.025573 |
| AAEL011177 | 45.18812 | 0.799456 | 0.243741 | 3.279943 | 0.001038 | 0.025858 |
| AAEL010671 | 24.07153 | 0.894153 | 0.27301 | 3.275168 | 0.001056 | 0.026133 |
| AAEL005749 | 251.0069 | 0.686759 | 0.210396 | 3.26412 | 0.001098 | 0.027003 |
| AAEL010076 | 104.2662 | 0.757702 | 0.232393 | 3.260436 | 0.001112 | 0.027242 |
| AAEL004673 | 175.841 | 0.747257 | 0.229475 | 3.256369 | 0.001128 | 0.027521 |
| AAEL006361 | 16.80501 | 0.972718 | 0.298805 | 3.255364 | 0.001132 | 0.027562 |
| AAEL025296 | 65.94521 | 0.933498 | 0.286871 | 3.254068 | 0.001138 | 0.02763 |
| AAEL003738 | 143.0541 | 0.947385 | 0.29123 | 3.25305 | 0.001142 | 0.027672 |
| AAEL000663 | 686.7743 | 0.701535 | 0.215908 | 3.249228 | 0.001157 | 0.027989 |
| AAEL001184 | 113.6894 | 0.715922 | 0.220444 | 3.247643 | 0.001164 | 0.028057 |
| AAEL004344 | 61.92219 | 0.946255 | 0.291754 | 3.243337 | 0.001181 | 0.028399 |
| AAEL021583 | 361.7207 | 0.644956 | 0.199062 | 3.239973 | 0.001195 | 0.028678 |
| AAEL003067 | 199.4711 | 0.735418 | 0.227297 | 3.235489 | 0.001214 | 0.029013 |
| AAEL005753 | 253.4142 | 0.903745 | 0.279277 | 3.236021 | 0.001212 | 0.029013 |
| AAEL009232 | 719.261 | 0.59774 | 0.184799 | 3.234544 | 0.001218 | 0.029051 |
| AAEL029031 | 210.3387 | 0.899357 | 0.278608 | 3.228032 | 0.001246 | 0.02965 |
| AAEL014662 | 100.3897 | 0.820568 | 0.254284 | 3.226967 | 0.001251 | 0.02965 |
| AAEL002853 | 353.8561 | 0.783151 | 0.242955 | 3.223439 | 0.001267 | 0.029958 |
| AAEL010125 | 452.0773 | 0.827231 | 0.257058 | 3.218076 | 0.001291 | 0.03028 |
| AAEL022387 | 1555.035 | 0.839831 | 0.261106 | 3.216431 | 0.001298 | 0.030333 |
| AAEL003516 | 35.32607 | 0.818741 | 0.254859 | 3.21252 | 0.001316 | 0.030688 |
| AAEL010451 | 55.59873 | 0.883062 | 0.275001 | 3.211126 | 0.001322 | 0.030776 |
| AAEL007806 | 95.1836 | 0.824052 | 0.256984 | 3.20662 | 0.001343 | 0.031017 |
| AAEL008404 | 15.78988 | 0.927311 | 0.290059 | 3.196973 | 0.001389 | 0.031823 |
| AAEL003611 | 503.1464 | 0.749426 | 0.235086 | 3.187881 | 0.001433 | 0.032587 |
| AAEL019955 | 56.06252 | 0.818249 | 0.257095 | 3.182672 | 0.001459 | 0.03305 |
| AAEL008936 | 505.5039 | 0.897201 | 0.282588 | 3.174942 | 0.001499 | 0.033684 |
| AAEL001402 | 1624.63 | 0.803909 | 0.253299 | 3.173761 | 0.001505 | 0.033757 |

| | | | | | | |
|---|---|---|---|---|---|---|
| AAEL002394 | 68.52411 | 0.964397 | 0.303931 | 3.173083 | 0.001508 | 0.033771 |
| AAEL020430 | 53.68024 | 0.869754 | 0.274401 | 3.169644 | 0.001526 | 0.03398 |
| AAEL018150 | 256.8273 | 0.637242 | 0.201189 | 3.167374 | 0.001538 | 0.034052 |
| AAEL028228 | 71.79854 | 0.60176 | 0.190749 | 3.154719 | 0.001607 | 0.035037 |
| AAEL008628 | 11.60906 | 0.936794 | 0.297523 | 3.148643 | 0.00164 | 0.035707 |
| AAEL019527 | 98.32211 | 0.936215 | 0.297447 | 3.147499 | 0.001647 | 0.035715 |
| AAEL010772 | 12.71458 | 0.935409 | 0.297307 | 3.146271 | 0.001654 | 0.035733 |
| AAEL004513 | 501.493 | 0.672957 | 0.214319 | 3.139977 | 0.00169 | 0.036243 |
| AAEL004581 | 115.7052 | 0.747296 | 0.238105 | 3.138521 | 0.001698 | 0.036292 |
| AAEL006754 | 188.9597 | 0.729288 | 0.233323 | 3.125656 | 0.001774 | 0.037643 |
| AAEL007603 | 91.07154 | 0.790994 | 0.253281 | 3.122995 | 0.00179 | 0.037917 |
| AAEL013352 | 165.8765 | 0.899788 | 0.28834 | 3.120586 | 0.001805 | 0.038091 |
| AAEL001519 | 89.37989 | 0.754631 | 0.241797 | 3.120924 | 0.001803 | 0.038091 |
| AAEL010480 | 171.9286 | 0.712333 | 0.228505 | 3.117365 | 0.001825 | 0.038304 |
| AAEL007807 | 291.0773 | 0.709335 | 0.227497 | 3.118003 | 0.001821 | 0.038304 |
| AAEL002964 | 372.1749 | 0.708892 | 0.227773 | 3.112275 | 0.001857 | 0.038639 |
| AAEL003978 | 86.0666 | 0.734852 | 0.23628 | 3.11009 | 0.00187 | 0.038639 |
| AAEL012099 | 1071.938 | 0.644528 | 0.207692 | 3.103283 | 0.001914 | 0.039126 |
| AAEL006027 | 369.2281 | 0.712718 | 0.229633 | 3.103733 | 0.001911 | 0.039126 |
| AAEL003954 | 244.363 | 0.899993 | 0.290701 | 3.095942 | 0.001962 | 0.0399 |
| AAEL025177 | 7.287182 | 0.937595 | 0.30297 | 3.094679 | 0.00197 | 0.040001 |
| AAEL002031 | 364.7844 | 0.662513 | 0.214363 | 3.090608 | 0.001997 | 0.040414 |
| AAEL019752 | 96.5247 | 0.935317 | 0.302789 | 3.089012 | 0.002008 | 0.040456 |
| AAEL024216 | 36.26313 | 0.814318 | 0.263575 | 3.089518 | 0.002005 | 0.040456 |
| AAEL007131 | 41.35659 | 0.902229 | 0.2921 | 3.088769 | 0.00201 | 0.040456 |
| AAEL007830 | 93.73634 | 0.803854 | 0.260699 | 3.083455 | 0.002046 | 0.040975 |
| AAEL013367 | 27.47528 | 0.830021 | 0.26916 | 3.08375 | 0.002044 | 0.040975 |
| AAEL002353 | 44.52078 | 0.883784 | 0.287199 | 3.077257 | 0.002089 | 0.041765 |
| AAEL002397 | 270.8589 | 0.672917 | 0.218925 | 3.073726 | 0.002114 | 0.042119 |
| AAEL013111 | 417.0919 | 0.606105 | 0.197237 | 3.072975 | 0.002119 | 0.042154 |
| AAEL000786 | 663.9844 | 0.613819 | 0.199955 | 3.069792 | 0.002142 | 0.04253 |
| AAEL026214 | 54.25816 | 0.931681 | 0.303943 | 3.065316 | 0.002174 | 0.042958 |
| AAEL012110 | 1114.376 | 0.680283 | 0.222055 | 3.063574 | 0.002187 | 0.043064 |
| AAEL009112 | 150.0326 | 0.75735 | 0.247619 | 3.058532 | 0.002224 | 0.043563 |

| | | | | | | |
|---|---|---|---|---|---|---|
| AAEL002288 | 177.7313 | 0.701476 | 0.229506 | 3.056461 | 0.00224 | 0.043733 |
| AAEL010164 | 136.0554 | 0.926719 | 0.303441 | 3.054033 | 0.002258 | 0.044015 |
| AAEL022578 | 1111.711 | 0.651825 | 0.213518 | 3.052779 | 0.002267 | 0.044109 |
| AAEL026481 | 16.45832 | 0.923289 | 0.302529 | 3.051906 | 0.002274 | 0.044109 |
| AAEL014426 | 613.9428 | 0.78692 | 0.258058 | 3.049398 | 0.002293 | 0.044333 |
| AAEL011130 | 220.0641 | 0.84931 | 0.278737 | 3.046994 | 0.002311 | 0.044542 |
| AAEL010960 | 497.0209 | 0.590991 | 0.194244 | 3.042514 | 0.002346 | 0.044989 |
| AAEL006682 | 268.142 | 0.600672 | 0.19767 | 3.038753 | 0.002376 | 0.045455 |
| AAEL024717 | 43.29953 | 0.7425 | 0.245675 | 3.022284 | 0.002509 | 0.04726 |
| AAEL022593 | 36.32292 | 0.748915 | 0.247928 | 3.020691 | 0.002522 | 0.047355 |
| AAEL013349 | 138.7392 | 0.916768 | 0.303543 | 3.020225 | 0.002526 | 0.047355 |
| AAEL001503 | 250.6286 | 0.616026 | 0.203919 | 3.02094 | 0.00252 | 0.047355 |
| AAEL007792 | 188.4432 | 0.663433 | 0.219998 | 3.015624 | 0.002565 | 0.047905 |
| AAEL014349 | 162.4543 | 0.819806 | 0.271922 | 3.014853 | 0.002571 | 0.047905 |
| AAEL020706 | 171.17 | 0.799672 | 0.265249 | 3.014796 | 0.002572 | 0.047905 |
| AAEL019917 | 182.4769 | 0.863065 | 0.287364 | 3.003385 | 0.00267 | 0.049348 |
| AAEL027068 | 16.06901 | 0.910432 | 0.303506 | 2.999714 | 0.002702 | 0.049542 |
| AAEL009483 | 396.3347 | 0.624344 | 0.208195 | 2.998836 | 0.00271 | 0.049542 |
| AAEL002632 | 20.76006 | 0.853735 | 0.28468 | 2.998925 | 0.002709 | 0.049542 |
| AAEL005142 | 139.5816 | 0.697069 | 0.232773 | 2.994635 | 0.002748 | 0.049768 |
| AAEL000650 | 518.4899 | 0.829376 | 0.277 | 2.994139 | 0.002752 | 0.049773 |

FC: fold change; P-adj: adjusted *p* value

**7 dpi 28 ºC downregulated**

| **GeneID** | **Base mean** | **log2(FC)** | **StdErr** | **Wald-Stats** | **P-value** | **P-adj** |
|---|---|---|---|---|---|---|
| AAEL006539 | 796.8858 | -0.73678 | 0.152472 | -4.83225 | 1.35E-06 | 0.000174 |
| AAEL006548 | 103.4199 | -0.9033 | 0.196717 | -4.59187 | 4.39E-06 | 0.000461 |
| AAEL001985 | 31.22873 | -0.89225 | 0.206364 | -4.32368 | 1.53E-05 | 0.001288 |
| AAEL019885 | 4768.937 | -0.67283 | 0.157481 | -4.27249 | 1.93E-05 | 0.001487 |
| AAEL021308 | 699.5122 | -0.83538 | 0.197223 | -4.2357 | 2.28E-05 | 0.001653 |
| AAEL012725 | 4190.36 | -0.63749 | 0.152979 | -4.16718 | 3.08E-05 | 0.002072 |
| AAEL024284 | 92.68213 | -1.24023 | 0.303653 | -4.08437 | 4.42E-05 | 0.00275 |
| AAEL003945 | 1538.941 | -0.59791 | 0.14691 | -4.06988 | 4.70E-05 | 0.002895 |
| AAEL007472 | 152.4488 | -0.75298 | 0.185976 | -4.04879 | 5.15E-05 | 0.003103 |

| | | | | | | |
|---|---|---|---|---|---|---|
| AAEL015566 | 13.39188 | -1.16857 | 0.302098 | -3.86818 | 0.00011 | 0.005621 |
| AAEL019889 | 315.843 | -0.78786 | 0.205865 | -3.82706 | 0.00013 | 0.00622 |
| AAEL018234 | 311.1289 | -0.76938 | 0.201293 | -3.82217 | 0.000132 | 0.00627 |
| AAEL012762 | 38.69028 | -0.82461 | 0.21902 | -3.76501 | 0.000167 | 0.007493 |
| AAEL021449 | 19.00106 | -1.12455 | 0.303932 | -3.7 | 0.000216 | 0.009183 |
| AAEL024757 | 12.57881 | -1.117 | 0.302825 | -3.68858 | 0.000226 | 0.009369 |
| AAEL006538 | 1261.145 | -0.72467 | 0.199573 | -3.63111 | 0.000282 | 0.010876 |
| AAEL005403 | 195.1592 | -0.72627 | 0.204596 | -3.54977 | 0.000386 | 0.013529 |
| AAEL002173 | 46.9797 | -1.07221 | 0.303871 | -3.52849 | 0.000418 | 0.014323 |
| AAEL011023 | 5220.941 | -0.61058 | 0.175352 | -3.48202 | 0.000498 | 0.016027 |
| AAEL012377 | 182.5946 | -0.87641 | 0.252186 | -3.47526 | 0.00051 | 0.016258 |
| AAEL006556 | 1382.892 | -0.7928 | 0.232545 | -3.40923 | 0.000651 | 0.018955 |
| AAEL026107 | 37.50244 | -0.95965 | 0.283578 | -3.38407 | 0.000714 | 0.019989 |
| AAEL022124 | 30.82317 | -1.01051 | 0.300338 | -3.36458 | 0.000767 | 0.020908 |
| AAEL001997 | 605.8789 | -0.68818 | 0.206507 | -3.33248 | 0.000861 | 0.022583 |
| AAEL008454 | 117.2711 | -0.96631 | 0.292072 | -3.30848 | 0.000938 | 0.024129 |
| AAEL020075 | 275.3158 | -0.93843 | 0.286306 | -3.27772 | 0.001046 | 0.026007 |
| AAEL010362 | 138.3527 | -0.74182 | 0.230461 | -3.21886 | 0.001287 | 0.030257 |
| AAEL002753 | 972.7655 | -0.88924 | 0.278464 | -3.19339 | 0.001406 | 0.032095 |
| AAEL019629 | 348.0743 | -0.96619 | 0.303852 | -3.17982 | 0.001474 | 0.03325 |
| AAEL009777 | 23.26452 | -0.93079 | 0.292797 | -3.17897 | 0.001478 | 0.033283 |
| AAEL011202 | 4899.99 | -0.64138 | 0.202483 | -3.16755 | 0.001537 | 0.034052 |
| AAEL004576 | 55.46567 | -0.94456 | 0.303401 | -3.11325 | 0.00185 | 0.038634 |
| AAEL011416 | 20.50216 | -0.91963 | 0.296211 | -3.10465 | 0.001905 | 0.039098 |
| AAEL000305 | 69.42238 | -0.82838 | 0.269374 | -3.0752 | 0.002104 | 0.041984 |
| AAEL011499 | 19.02131 | -0.90922 | 0.298348 | -3.04753 | 0.002307 | 0.044536 |
| AAEL024830 | 179.1432 | -0.71887 | 0.237273 | -3.02971 | 0.002448 | 0.046411 |
| AAEL021576 | 21.2682 | -0.76916 | 0.255667 | -3.00844 | 0.002626 | 0.048769 |
| AAEL008183 | 1134.361 | -0.60889 | 0.20306 | -2.99857 | 0.002712 | 0.049542 |

FC: fold change; P-adj: adjusted *p* value

**7 dpi 32 ºC upregulated**

| GeneID | Base mean | log2(FC) | StdErr | Wald-Stats | P-value | P-adj |
|---|---|---|---|---|---|---|
| AAEL017976 | 446.7218 | 2.919349 | 0.359109 | 8.129415 | 4.31E-16 | 5.31E-12 |

| | | | | | | |
|---|---|---|---|---|---|---|
| AAEL027610 | 342.6254 | 2.880902 | 0.363423 | 7.927137 | 2.24E-15 | 1.38E-11 |
| AAEL006323 | 204.7075 | 1.811199 | 0.261353 | 6.930083 | 4.21E-12 | 1.72E-08 |
| AAEL017975 | 659.6938 | 2.424532 | 0.362353 | 6.691067 | 2.22E-11 | 6.81E-08 |
| AAEL006990 | 710.9443 | 1.738503 | 0.265983 | 6.536144 | 6.31E-11 | 1.55E-07 |
| AAEL020330 | 80.5189 | 2.272107 | 0.36694 | 6.192037 | 5.94E-10 | 1.04E-06 |
| AAEL023591 | 3057.906 | 1.555017 | 0.250685 | 6.20308 | 5.54E-10 | 1.04E-06 |
| AAEL001098 | 139.8757 | 1.82806 | 0.299515 | 6.103404 | 1.04E-09 | 1.42E-06 |
| AAEL026008 | 3314.774 | 1.576086 | 0.257949 | 6.110066 | 9.96E-10 | 1.42E-06 |
| AAEL027829 | 142.1775 | 2.10786 | 0.354259 | 5.95006 | 2.68E-09 | 3.30E-06 |
| AAEL014348 | 162.4059 | 1.266885 | 0.217238 | 5.831786 | 5.48E-09 | 5.88E-06 |
| AAEL001646 | 97.74182 | 1.763908 | 0.302848 | 5.824402 | 5.73E-09 | 5.88E-06 |
| AAEL001077 | 180.9688 | 1.285756 | 0.224488 | 5.727507 | 1.02E-08 | 9.64E-06 |
| AAEL014020 | 160.3409 | 2.096797 | 0.366975 | 5.713736 | 1.11E-08 | 9.71E-06 |
| AAEL013350 | 177.675 | 2.077968 | 0.367169 | 5.659434 | 1.52E-08 | 1.25E-05 |
| AAEL003345 | 9199.334 | 1.887363 | 0.336661 | 5.606128 | 2.07E-08 | 1.59E-05 |
| AAEL005985 | 4343.619 | 0.649724 | 0.116935 | 5.556259 | 2.76E-08 | 1.78E-05 |
| AAEL009127 | 823.6629 | 1.67233 | 0.300912 | 5.557547 | 2.74E-08 | 1.78E-05 |
| AAEL000037 | 232.6124 | 1.394313 | 0.250482 | 5.566527 | 2.60E-08 | 1.78E-05 |
| AAEL003728 | 266.4348 | 1.480512 | 0.269575 | 5.492032 | 3.97E-08 | 2.44E-05 |
| AAEL011203 | 68.67572 | 1.344507 | 0.247184 | 5.439291 | 5.35E-08 | 3.13E-05 |
| AAEL023844 | 316.0924 | 1.360717 | 0.250813 | 5.425223 | 5.79E-08 | 3.24E-05 |
| AAEL003505 | 1961.892 | 1.117491 | 0.209226 | 5.341064 | 9.24E-08 | 4.94E-05 |
| AAEL013770 | 116.996 | 1.559145 | 0.294503 | 5.294154 | 1.20E-07 | 6.13E-05 |
| AAEL008473 | 3062.51 | 1.718822 | 0.32574 | 5.276673 | 1.32E-07 | 6.47E-05 |
| AAEL022208 | 91.06876 | 1.446752 | 0.274668 | 5.267276 | 1.38E-07 | 6.55E-05 |
| AAEL019751 | 776.5939 | 1.287111 | 0.250391 | 5.140404 | 2.74E-07 | 0.00012 |
| AAEL022059 | 87.8943 | 1.848594 | 0.367529 | 5.029791 | 4.91E-07 | 0.000208 |
| AAEL023999 | 19.01198 | 1.421984 | 0.285488 | 4.980893 | 6.33E-07 | 0.00026 |
| AAEL001087 | 547.9228 | 1.656446 | 0.333591 | 4.965506 | 6.85E-07 | 0.000272 |
| AAEL006883 | 538.1236 | 1.584296 | 0.320162 | 4.948414 | 7.48E-07 | 0.000288 |
| AAEL002610 | 1194.176 | 1.528829 | 0.309799 | 4.934908 | 8.02E-07 | 0.000299 |
| AAEL014045 | 379.1003 | 1.57201 | 0.320175 | 4.909841 | 9.12E-07 | 0.00033 |
| AAEL022253 | 340.4038 | 1.800957 | 0.367743 | 4.89732 | 9.72E-07 | 0.000341 |
| AAEL022334 | 1089.324 | 1.280693 | 0.262337 | 4.881862 | 1.05E-06 | 0.000359 |

| | | | | | | |
|---|---|---|---|---|---|---|
| AAEL000825 | 20.46806 | 1.741355 | 0.35981 | 4.839651 | 1.30E-06 | 0.000421 |
| AAEL012764 | 1407.784 | 1.42573 | 0.29495 | 4.833795 | 1.34E-06 | 0.000423 |
| AAEL022079 | 96.07221 | 1.36087 | 0.284021 | 4.791448 | 1.66E-06 | 0.000497 |
| AAEL013284 | 280.9838 | 1.748619 | 0.367739 | 4.755048 | 1.98E-06 | 0.000581 |
| AAEL001594 | 91.54731 | 1.528298 | 0.32188 | 4.74804 | 2.05E-06 | 0.000588 |
| AAEL009850 | 110.7414 | 1.001692 | 0.212355 | 4.717067 | 2.39E-06 | 0.000654 |
| AAEL009198 | 828.4534 | 0.74214 | 0.15723 | 4.720089 | 2.36E-06 | 0.000654 |
| AAEL008274 | 1180.8 | 0.943283 | 0.200634 | 4.701506 | 2.58E-06 | 0.000691 |
| AAEL011038 | 590.2102 | 0.995029 | 0.211846 | 4.696952 | 2.64E-06 | 0.000691 |
| AAEL001084 | 129.3413 | 1.432871 | 0.308204 | 4.649092 | 3.33E-06 | 0.000837 |
| AAEL010434 | 58.39876 | 1.395248 | 0.300437 | 4.644068 | 3.42E-06 | 0.00084 |
| AAEL003626 | 316.58 | 1.332189 | 0.287698 | 4.630514 | 3.65E-06 | 0.00088 |
| AAEL019504 | 1039.776 | 1.410592 | 0.304987 | 4.625083 | 3.74E-06 | 0.000886 |
| AAEL017334 | 3153.811 | 1.621586 | 0.352744 | 4.597059 | 4.28E-06 | 0.000995 |
| AAEL001813 | 234.0416 | 1.543163 | 0.336516 | 4.585708 | 4.52E-06 | 0.001012 |
| AAEL021302 | 754.1271 | 1.259606 | 0.274511 | 4.58855 | 4.46E-06 | 0.001012 |
| AAEL025126 | 82.35019 | 1.683724 | 0.36779 | 4.577951 | 4.70E-06 | 0.001031 |
| AAEL014510 | 176.4301 | 0.875211 | 0.19136 | 4.573642 | 4.79E-06 | 0.001034 |
| AAEL007765 | 3748.655 | 0.617202 | 0.136073 | 4.535817 | 5.74E-06 | 0.001217 |
| AAEL001949 | 67.29029 | 0.96705 | 0.214315 | 4.51229 | 6.41E-06 | 0.001315 |
| AAEL013812 | 464.8684 | 1.220807 | 0.27322 | 4.468219 | 7.89E-06 | 0.001591 |
| AAEL005045 | 230.6594 | 1.621136 | 0.363703 | 4.457303 | 8.30E-06 | 0.001621 |
| AAEL000044 | 884.8495 | 1.444323 | 0.323811 | 4.460381 | 8.18E-06 | 0.001621 |
| AAEL001241 | 44.46046 | 1.263618 | 0.284338 | 4.444064 | 8.83E-06 | 0.001688 |
| AAEL012712 | 243.566 | 1.454281 | 0.330154 | 4.404857 | 1.06E-05 | 0.001943 |
| AAEL006581 | 168.3474 | 1.086651 | 0.246615 | 4.406262 | 1.05E-05 | 0.001943 |
| AAEL007218 | 307.9408 | 0.998626 | 0.227978 | 4.380358 | 1.18E-05 | 0.002143 |
| AAEL024179 | 3549.131 | 0.830567 | 0.190646 | 4.356583 | 1.32E-05 | 0.002355 |
| AAEL019590 | 200.524 | 1.329421 | 0.306152 | 4.342351 | 1.41E-05 | 0.002453 |
| AAEL010393 | 19.65437 | 1.556713 | 0.358575 | 4.341384 | 1.42E-05 | 0.002453 |
| AAEL010076 | 208.0212 | 1.270966 | 0.293128 | 4.335867 | 1.45E-05 | 0.00248 |
| AAEL013345 | 35.93927 | 1.583963 | 0.367557 | 4.309437 | 1.64E-05 | 0.002732 |
| AAEL010379 | 577.0775 | 1.267505 | 0.295651 | 4.287161 | 1.81E-05 | 0.002948 |
| AAEL006805 | 1722.145 | 1.145897 | 0.267372 | 4.28577 | 1.82E-05 | 0.002948 |

| | | | | | | |
|---|---|---|---|---|---|---|
| AAEL007090 | 226.3581 | 1.407091 | 0.329313 | 4.272814 | 1.93E-05 | 0.003084 |
| AAEL007902 | 293.3958 | 1.083611 | 0.253791 | 4.269701 | 1.96E-05 | 0.003087 |
| AAEL001293 | 335.6991 | 1.421436 | 0.334346 | 4.251392 | 2.12E-05 | 0.003308 |
| AAEL013346 | 44.75176 | 1.523031 | 0.360531 | 4.224415 | 2.40E-05 | 0.003638 |
| AAEL008622 | 121.9277 | 1.508754 | 0.357862 | 4.216025 | 2.49E-05 | 0.00373 |
| AAEL017345 | 1297.581 | 1.064248 | 0.252898 | 4.208207 | 2.57E-05 | 0.003815 |
| AAEL003954 | 557.3156 | 1.458347 | 0.347417 | 4.197683 | 2.70E-05 | 0.003949 |
| AAEL007387 | 919.4093 | 0.662786 | 0.158004 | 4.194744 | 2.73E-05 | 0.003953 |
| AAEL007126 | 180.1804 | 1.368633 | 0.326626 | 4.190219 | 2.79E-05 | 0.003986 |
| AAEL027654 | 931.9174 | 1.162701 | 0.278422 | 4.17604 | 2.97E-05 | 0.004194 |
| AAEL026537 | 18.50074 | 1.429866 | 0.342787 | 4.171298 | 3.03E-05 | 0.004234 |
| AAEL003708 | 104.9343 | 1.061444 | 0.25543 | 4.155524 | 3.25E-05 | 0.004436 |
| AAEL002917 | 824.9129 | 1.358663 | 0.328008 | 4.142165 | 3.44E-05 | 0.004541 |
| AAEL015465 | 195.1952 | 1.123156 | 0.271441 | 4.137748 | 3.51E-05 | 0.004541 |
| AAEL014556 | 983.993 | 1.218744 | 0.294542 | 4.137754 | 3.51E-05 | 0.004541 |
| AAEL017380 | 38.84343 | 1.523205 | 0.367482 | 4.144982 | 3.40E-05 | 0.004541 |
| AAEL006904 | 1600.12 | 0.837621 | 0.203669 | 4.112657 | 3.91E-05 | 0.004904 |
| AAEL014539 | 51.32586 | 1.348928 | 0.32816 | 4.11058 | 3.95E-05 | 0.004904 |
| AAEL005533 | 90.23157 | 1.49545 | 0.364803 | 4.099338 | 4.14E-05 | 0.005054 |
| AAEL002658 | 301.2503 | 0.950003 | 0.231764 | 4.099005 | 4.15E-05 | 0.005054 |
| AAEL010337 | 337.9218 | 1.240084 | 0.303146 | 4.090715 | 4.30E-05 | 0.005136 |
| AAEL008635 | 1277.263 | 1.398547 | 0.343442 | 4.072155 | 4.66E-05 | 0.005457 |
| AAEL008532 | 72.24792 | 1.497715 | 0.367772 | 4.072407 | 4.65E-05 | 0.005457 |
| AAEL010206 | 397.5056 | 1.162759 | 0.285702 | 4.069832 | 4.70E-05 | 0.00546 |
| AAEL014246 | 3290.577 | 1.202206 | 0.29701 | 4.047694 | 5.17E-05 | 0.005891 |
| AAEL010769 | 538.2967 | 1.162549 | 0.287645 | 4.041606 | 5.31E-05 | 0.005991 |
| AAEL007989 | 7.168395 | 1.440137 | 0.359007 | 4.011445 | 6.03E-05 | 0.006724 |
| AAEL000811 | 360.2371 | 1.207735 | 0.301169 | 4.01016 | 6.07E-05 | 0.006724 |
| AAEL021929 | 175.7643 | 1.46841 | 0.3671 | 4.000025 | 6.33E-05 | 0.006956 |
| AAEL014604 | 162.4537 | 0.975968 | 0.245487 | 3.975643 | 7.02E-05 | 0.007574 |
| AAEL003656 | 271.4358 | 0.949151 | 0.239128 | 3.96922 | 7.21E-05 | 0.007713 |
| AAEL004118 | 662.4558 | 1.139949 | 0.289203 | 3.94169 | 8.09E-05 | 0.00858 |
| AAEL012457 | 618.3535 | 1.328268 | 0.338319 | 3.926079 | 8.63E-05 | 0.008925 |
| AAEL022982 | 610.0555 | 0.997812 | 0.254062 | 3.927434 | 8.59E-05 | 0.008925 |

| | | | | | | |
|---|---|---|---|---|---|---|
| AAEL014335 | 516.5232 | 0.944089 | 0.24138 | 3.911209 | 9.18E-05 | 0.009414 |
| AAEL012856 | 783.5518 | 1.389763 | 0.356661 | 3.896596 | 9.76E-05 | 0.009917 |
| AAEL026161 | 396.1913 | 1.190966 | 0.306088 | 3.89092 | 9.99E-05 | 0.010004 |
| AAEL013163 | 994.9419 | 0.912059 | 0.236521 | 3.856147 | 0.000115 | 0.011245 |
| AAEL014419 | 138.4878 | 1.189442 | 0.310149 | 3.83507 | 0.000126 | 0.012063 |
| AAEL002301 | 595.0697 | 1.08825 | 0.283741 | 3.835368 | 0.000125 | 0.012063 |
| AAEL027362 | 23.11579 | 1.342468 | 0.350727 | 3.827672 | 0.000129 | 0.012335 |
| AAEL008401 | 27.55802 | 1.396174 | 0.365882 | 3.81591 | 0.000136 | 0.012839 |
| AAEL006321 | 2850.859 | 0.650008 | 0.17071 | 3.807685 | 0.00014 | 0.013172 |
| AAEL001667 | 530.0897 | 1.233206 | 0.325737 | 3.785899 | 0.000153 | 0.014059 |
| AAEL013257 | 44.33485 | 0.882617 | 0.234425 | 3.765024 | 0.000167 | 0.015063 |
| AAEL014541 | 277.04 | 1.183489 | 0.314746 | 3.760144 | 0.00017 | 0.015247 |
| AAEL003632 | 22.32452 | 1.236254 | 0.328957 | 3.758101 | 0.000171 | 0.015261 |
| AAEL012640 | 86.26759 | 1.362207 | 0.364137 | 3.740917 | 0.000183 | 0.016226 |
| AAEL000915 | 292.4677 | 1.284868 | 0.344692 | 3.727585 | 0.000193 | 0.016747 |
| AAEL023509 | 149.1324 | 1.001252 | 0.270994 | 3.694743 | 0.00022 | 0.018803 |
| AAEL001403 | 60.93914 | 0.922478 | 0.249908 | 3.691266 | 0.000223 | 0.018869 |
| AAEL011350 | 1004.095 | 0.927922 | 0.251447 | 3.690335 | 0.000224 | 0.018869 |
| AAEL024560 | 145.6707 | 0.83561 | 0.227987 | 3.665167 | 0.000247 | 0.020544 |
| AAEL010917 | 44.85952 | 1.332682 | 0.364534 | 3.655852 | 0.000256 | 0.021021 |
| AAEL002124 | 763.5451 | 0.95074 | 0.259965 | 3.657187 | 0.000255 | 0.021021 |
| AAEL007453 | 25.62713 | 1.318314 | 0.362529 | 3.636434 | 0.000276 | 0.022424 |
| AAEL007784 | 158.5976 | 1.330548 | 0.367271 | 3.622796 | 0.000291 | 0.023129 |
| AAEL004844 | 11.5029 | 1.234893 | 0.341244 | 3.618792 | 0.000296 | 0.023339 |
| AAEL002075 | 38.14323 | 1.281871 | 0.355178 | 3.609097 | 0.000307 | 0.024074 |
| AAEL010480 | 226.9322 | 1.035451 | 0.287839 | 3.597329 | 0.000322 | 0.02503 |
| AAEL010068 | 1383.658 | 1.106547 | 0.307802 | 3.59499 | 0.000324 | 0.025097 |
| AAEL013525 | 617.8744 | 0.749726 | 0.208717 | 3.592064 | 0.000328 | 0.025222 |
| AAEL027157 | 337.3348 | 0.878586 | 0.244974 | 3.586445 | 0.000335 | 0.025612 |
| AAEL013159 | 28.46296 | 1.166178 | 0.325735 | 3.58014 | 0.000343 | 0.026076 |
| AAEL020502 | 325.3333 | 0.91947 | 0.257584 | 3.569592 | 0.000358 | 0.026817 |
| AAEL025574 | 808.3258 | 0.985478 | 0.276021 | 3.570302 | 0.000357 | 0.026817 |
| AAEL007789 | 132.4106 | 0.786266 | 0.220558 | 3.564895 | 0.000364 | 0.027137 |
| AAEL026819 | 109.2034 | 0.922467 | 0.258952 | 3.562312 | 0.000368 | 0.0272 |

| | | | | | | |
|---|---|---|---|---|---|---|
| AAEL009629 | 3513.993 | 1.096565 | 0.307927 | 3.561119 | 0.000369 | 0.0272 |
| AAEL001420 | 7877.547 | 0.939145 | 0.264278 | 3.553619 | 0.00038 | 0.027821 |
| AAEL012701 | 2446.635 | 0.589274 | 0.165915 | 3.551665 | 0.000383 | 0.027863 |
| AAEL019641 | 592.2827 | 0.932353 | 0.263518 | 3.538094 | 0.000403 | 0.029163 |
| AAEL003123 | 1780.523 | 1.220832 | 0.345505 | 3.53347 | 0.00041 | 0.029504 |
| AAEL003713 | 245.679 | 0.861885 | 0.245095 | 3.516527 | 0.000437 | 0.03127 |
| AAEL009640 | 143.485 | 0.798098 | 0.227962 | 3.501016 | 0.000463 | 0.032766 |
| AAEL013885 | 6407.882 | 1.186245 | 0.338801 | 3.5013 | 0.000463 | 0.032766 |
| AAEL017071 | 967.1173 | 0.94955 | 0.273901 | 3.46676 | 0.000527 | 0.036609 |
| AAEL022589 | 70.16731 | 1.267022 | 0.367725 | 3.44557 | 0.00057 | 0.037802 |
| AAEL013109 | 209.294 | 0.958501 | 0.27825 | 3.444748 | 0.000572 | 0.037802 |
| AAEL003076 | 953.9536 | 1.051074 | 0.304708 | 3.449449 | 0.000562 | 0.037802 |
| AAEL007773 | 438.2615 | 1.034985 | 0.300092 | 3.448887 | 0.000563 | 0.037802 |
| AAEL019602 | 251.8779 | 0.835476 | 0.242534 | 3.444778 | 0.000572 | 0.037802 |
| AAEL001794 | 1266.131 | 1.029609 | 0.298217 | 3.452549 | 0.000555 | 0.037802 |
| AAEL002555 | 176.1723 | 1.030978 | 0.299896 | 3.437786 | 0.000586 | 0.038375 |
| AAEL018103 | 407.3152 | 1.138742 | 0.331503 | 3.435089 | 0.000592 | 0.038554 |
| AAEL008628 | 10.12739 | 1.241043 | 0.362826 | 3.42049 | 0.000625 | 0.040469 |
| AAEL009645 | 6653.894 | 0.856867 | 0.251118 | 3.412206 | 0.000644 | 0.040896 |
| AAEL002347 | 111.4024 | 1.241881 | 0.363978 | 3.411964 | 0.000645 | 0.040896 |
| AAEL026496 | 5.767867 | 1.193677 | 0.349698 | 3.413448 | 0.000641 | 0.040896 |
| AAEL013347 | 949.1318 | 1.039507 | 0.305001 | 3.408204 | 0.000654 | 0.04122 |
| AAEL005221 | 3658.496 | 0.913722 | 0.268189 | 3.407011 | 0.000657 | 0.04122 |
| AAEL014891 | 165.9633 | 1.144306 | 0.337079 | 3.394769 | 0.000687 | 0.042672 |
| AAEL002432 | 12.90657 | 1.221794 | 0.361287 | 3.381785 | 0.00072 | 0.04409 |
| AAEL010881 | 80.62625 | 0.828413 | 0.244971 | 3.381683 | 0.00072 | 0.04409 |
| AAEL004003 | 23.51713 | 1.11224 | 0.328874 | 3.38196 | 0.00072 | 0.04409 |
| AAEL005769 | 364.4213 | 1.109089 | 0.328161 | 3.379705 | 0.000726 | 0.044188 |
| AAEL028247 | 23.29229 | 1.227407 | 0.36364 | 3.375335 | 0.000737 | 0.044675 |
| AAEL019564 | 620.3279 | 1.028056 | 0.306753 | 3.351409 | 0.000804 | 0.047779 |
| AAEL026390 | 1302.221 | 0.608807 | 0.182065 | 3.343893 | 0.000826 | 0.048856 |
| AAEL013341 | 940.2407 | 0.80422 | 0.241003 | 3.336967 | 0.000847 | 0.049613 |
| AAEL018159 | 583.4205 | 0.966312 | 0.28956 | 3.337169 | 0.000846 | 0.049613 |

FC: fold change; P-adj: adjusted *p* value

**7 dpi 32 ºC downregulated**

| GeneID | Base mean | log2(FC) | StdErr | Wald-Stats | P-value | P-adj |
|---|---|---|---|---|---|---|
| AAEL024449 | 7.900429 | -1.89165 | 0.366118 | -5.16676 | 2.38E-07 | 0.000109 |
| AAEL006938 | 25.42905 | -1.65673 | 0.341677 | -4.84882 | 1.24E-06 | 0.000413 |
| AAEL020913 | 93.79685 | -0.84025 | 0.174434 | -4.81699 | 1.46E-06 | 0.000448 |
| AAEL018041 | 131.1753 | -1.70489 | 0.363607 | -4.68883 | 2.75E-06 | 0.000704 |
| AAEL021011 | 89.28689 | -1.15135 | 0.267228 | -4.30848 | 1.64E-05 | 0.002732 |
| AAEL020306 | 4.225345 | -1.54521 | 0.365169 | -4.23148 | 2.32E-05 | 0.00357 |
| AAEL000322 | 434.8694 | -0.81085 | 0.194666 | -4.16534 | 3.11E-05 | 0.004297 |
| AAEL000035 | 22.7378 | -1.51775 | 0.365904 | -4.14795 | 3.35E-05 | 0.004535 |
| AAEL024183 | 54.88617 | -1.51972 | 0.367732 | -4.13267 | 3.59E-05 | 0.004547 |
| AAEL006871 | 390.1212 | -0.66076 | 0.159871 | -4.13311 | 3.58E-05 | 0.004547 |
| AAEL013003 | 347.7431 | -1.09056 | 0.266392 | -4.09382 | 4.24E-05 | 0.005117 |
| AAEL005621 | 34.90218 | -1.49456 | 0.367646 | -4.0652 | 4.80E-05 | 0.005517 |
| AAEL004843 | 822.7724 | -0.70063 | 0.175736 | -3.98683 | 6.70E-05 | 0.007289 |
| AAEL005070 | 840.8001 | -0.81728 | 0.21007 | -3.89051 | 0.0001 | 0.010004 |
| AAEL023039 | 27.23422 | -1.28358 | 0.332377 | -3.86183 | 0.000113 | 0.011164 |
| AAEL005068 | 942.282 | -1.11388 | 0.293884 | -3.79021 | 0.000151 | 0.014027 |
| AAEL026029 | 11.85499 | -1.27482 | 0.346728 | -3.67673 | 0.000236 | 0.019769 |
| AAEL022225 | 19.50603 | -1.26525 | 0.347996 | -3.63582 | 0.000277 | 0.022424 |
| AAEL008435 | 746.1219 | -0.9798 | 0.269878 | -3.63054 | 0.000283 | 0.022739 |
| AAEL019689 | 202.5493 | -1.1764 | 0.337385 | -3.48682 | 0.000489 | 0.034359 |
| AAEL025193 | 13.09782 | -1.23745 | 0.35752 | -3.46122 | 0.000538 | 0.037161 |
| AAEL025401 | 534.8338 | -0.78726 | 0.228847 | -3.44011 | 0.000581 | 0.038251 |
| AAEL026468 | 471.9231 | -0.68725 | 0.204249 | -3.36477 | 0.000766 | 0.046194 |
| AAEL018184 | 46.58388 | -1.22761 | 0.365171 | -3.36172 | 0.000775 | 0.046478 |

FC: fold change; P-adj: adjusted *p* value

Supplementary Table 3. Number of DEGs.

| | | **Upregulated genes P-adj<0.05, FC>+1.5** | | **Downregulated genes P-adj<0.05, FC>-1.5** | |
|---|---|---|---|---|---|
| | | Total | Immune | Total | Immune |
| | 18ºC | 141 | 12 | 134 | 4 |
| **3 dpi** | 28ºC | 374 | 39 | 36 | 1 |
| | 32ºC | 26 | 2 | 4 | 4 |
| | 18ºC | 100 | 3 | 13 | 3 |
| **7 dpi** | 28ºC | 381 | 41 | 38 | 0 |
| | 32ºC | 170 | 18 | 24 | 0 |
| **Total** | | 1192 | 115 | 249 | 12 |

Adj: adjusted; FC: fold change

Supplementary Table 4. Classical and non-classical immune gene families and number of genes identified.

| Category | Gene family | Number of genes identified |
| --- | --- | --- |
| **Classical immune families** | Anti-microbial peptides | 12 |
| | Apoptosis | 29 |
| | Autophagy | 21 |
| | Catalase | 1 |
| | CLIP | 92 |
| | C-type lectins (CTL) | 48 |
| | Fibrinogen-related protein (FREP) | 31 |
| | Galectin | 10 |
| | Gram-negative binding proteins (GNBP) | 7 |
| | IMD pathway | 21 |
| | JAK-STAT pathway | 6 |
| | Leucine-rich repeat-containing proteins/ Leucine-rich repeat immune proteins (LRR/ LRIMs) | 56 |
| | Lysozymes | 5 |
| | Myeloid differentiation 2-related lipid recognition protein (ML) | 22 |
| | Peroxidase | 21 |
| | Peptidoglycan recognition proteins (PGRP) | 9 |
| | Prophenoloxidase (PPO) | 10 |
| | Relish | 3 |
| | Scavenger receptors (SCR) | 20 |
| | Serpins | 25 |
| | RNA inhibition pathway | 30 |
| | Superoxide dismutase (SOD) | 8 |
| | Spatzle | 7 |
| | Thioester proteins (TEP) | 11 |
| | Toll pathway | 16 |
| **Non-classical immune families** | Trypsin | 59 |
| | Serine proteases | 119 |

| | |
|---|---|
| Tubulin | 21 |
| Actin | 31 |
| Myosin | 23 |
| Lachesin | 6 |
| Heat shock protein | 12 |
| Cytochrome P450 | 150 |
| Lethal (2) essential for life protein, l2efl | 10 |
| Vacuolar ATPase | 14 |
| Sidestep proteins | 6 |
| Salivary proteins | 26 |
| **Total** | **998** |

Supplementary Table 5. Classical immune genes.

| | Gene ID | Name/Description |
|---|---|---|
| **Classical** | | |
| **AMP** | AAEL003389 | ATT |
| | AAEL029038 | CECA |
| | AAEL029046 | CECD |
| | AAEL029044 | CECE |
| | AAEL029041 | CECD |
| | AAEL029047 | CECN |
| | AAEL027792 | DEFA |
| | AAEL003832 | DEFC |
| | AAEL003857 | DEFD |
| | AAEL004833 | DPT1 |
| | AAEL004522 | GAM1 |
| | AAEL017536 | Holotricin glycine rich repeat protein (GRRP) anti-microbial peptide |
| **Autophagy** | AAEL009089 | APG12 |
| | AAEL019779 | APG16L |
| | AAEL013063 | APG18A |
| | AAEL013995 | APG18B |
| | AAEL000955 | APG3 |
| | AAEL010516 | APG4A |
| | AAEL007228 | APG4B |
| | AAEL002286 | APG5 |
| | AAEL010427 | APG6 |
| | AAEL010641 | APG7A |
| | AAEL012306 | APG7B |
| | AAEL007162 | APG8 |
| | AAEL009105 | APG9 |
| | AAEL001515 | DEBCL |
| | AAEL003777 | APG2 |
| | AAEL020638 | TOR |
| | AAEL009814 | Autophagy related gene |
| | AAEL010791 | Autophagy-specific protein, putative |
| | AAEL021581 | BUFFY |
| | AAEL019922 | APG1 |
| | AAEL021061 | APG10 |
| **GNBP** | AAEL000652 | GNBPA2 |
| | AAEL003889 | GNBPB1 |
| | AAEL003894 | GNBPB5 |
| | AAEL007064 | GNBPB6 |
| | AAEL007626 | GNBPA1 |
| | AAEL009176 | GNBPB3 |

| | | |
|---|---|---|
| | AAEL009178 | GNBPB4 |
| **Caspase and apoptosis** | AAEL014148 | Dredd |
| | AAEL026744 | CASPS9 |
| | AAEL011562 | Dronc |
| | AAEL005956 | CASPS16 |
| | AAEL005955 | CASPS17 |
| | AAEL003439 | CASPS18 |
| | AAEL003444 | CASPS19 |
| | AAEL017498 | CASPS21 |
| | AAEL012143 | CASPS7 |
| | AAEL014348 | CASPS8 |
| | AAEL000874 | ARK |
| | AAEL004392 | IMP |
| | AAEL014196 | Michelob-x |
| | AAEL011277 | Apoptosis stimulating of p53 |
| | AAEL009642 | Cathepsin B |
| | AAEL009637 | Cathepsin B |
| | AAEL000420 | Cathepsin O |
| | AAEL002833 | Cathepsin L |
| | AAEL011167 | Cathepsin L |
| | AAEL006389 | Cathepsin L |
| | AAEL006633 | IAP2 |
| | AAEL009074 | IAP1 |
| | AAEL007713 | Viral IAP-associated factor, putative |
| | AAEL010486 | Viral IAP-associated factor, putative |
| | AAEL011096 | Viral IAP-associated factor, putative |
| | AAEL012446 | IAP6 |
| | AAEL025438 | Wengen |
| | AAEL008634 | JNK |
| | AAEL014251 | IAP5 |
| **CLIP** | AAEL002601 | CLIPA1 |
| | AAEL019853 | CLIP |
| | AAEL019781 | CLIP |
| | AAEL015430 | CLIP |
| | AAEL026876 | CLIP |
| | AAEL019590 | CLIP |
| | AAEL000224 | CLIP |
| | AAEL026937 | CLIP |
| | AAEL005718 | CLIPA3 |
| | AAEL002288 | CLIPA4 |
| | AAEL002629 | CLIPA6 |
| | AAEL001675 | CLIPA10 |
| | AAEL002585 | CLIPA11 |
| | AAEL002590 | CLIPA12 |

| | |
|---|---|
| AAEL002595 | CLIPA14 |
| AAEL002126 | CLIPA15 |
| AAEL008404 | CLIPA16 |
| AAEL008668 | CLIP |
| AAEL000074 | CLIPB1 |
| AAEL005064 | CLIPB5 |
| AAEL003243 | CLIPB13A |
| AAEL003253 | CLIPB13B |
| AAEL014349 | CLIPB15 |
| AAEL005648 | CLIPB16 |
| AAEL007006 | CLIPA17 |
| AAEL001084 | CLIPB21 |
| AAEL014140 | CLIPB24 |
| AAEL014137 | CLIPB25 |
| AAEL007993 | CLIPB27 |
| AAEL013245 | CLIPB28 |
| AAEL006674 | CLIPB29 |
| AAEL000760 | CLIPB30 |
| AAEL006161 | CLIPB31 |
| AAEL000099 | CLIPB33 |
| AAEL000028 | CLIPB34 |
| AAEL000037 | CLIPB35 |
| AAEL000038 | CLIPB6-B36 |
| AAEL005431 | CLIPB37 |
| AAEL003628 | CLIPB38 |
| AAEL003632 | CLIPB39 |
| AAEL003631 | CLIPB41 |
| AAEL006168 | CLIPB42 |
| AAEL014354 | CLIPB43 |
| AAEL005060 | CLIPB44 |
| AAEL001077 | CLIPB45 |
| AAEL005093 | CLIPB46 |
| AAEL027429 | CLIPB76 |
| AAEL014139 | CLIPB79 |
| AAEL017003 | Clip-domain serine protease, family B |
| AAEL011991 | CLIPC1 |
| AAEL007593 | CLIPC2 |
| AAEL007597 | CLIPC3 |
| AAEL004524 | CLIPC5B |
| AAEL011593 | CLIPC11 |
| AAEL012711 | CLIPC12 |
| AAEL012712 | CLIPC13 |
| AAEL004948 | CLIPC14 |
| AAEL010270 | CLIPC15 |

| | | |
|---|---|---|
| | AAEL012713 | CLIPC16 |
| | AAEL007796 | CLIPD1 |
| | AAEL004979 | CLIPD2 |
| | AAEL002997 | CLIPD3 |
| | AAEL002124 | CLIPD6 |
| | AAEL005906 | CLIPD8 |
| | AAEL000238 | CLIPD9 |
| | AAEL015109 | CLIPD10 |
| | AAEL011375 | CLIPD11 |
| | AAEL005792 | CLIPE8 |
| | AAEL001233 | CLIPE9 |
| | AAEL010773 | CLIPE10 |
| | AAEL018347 | CLIPE12 |
| | AAEL019767 | SP-CLIP-SP |
| | AAEL006689 | CUBSP1 |
| | AAEL006696 | CUBSP2 |
| | AAEL006700 | CUBSP3 |
| | AAEL006703 | CUBSP4 |
| | AAEL016975 | ZFSP |
| | AAEL023229 | HP14 |
| | AAEL005748 | CBSP |
| | AAEL027371 | SPH145 |
| | AAEL013413 | IgSP2 |
| | AAEL014367 | SEASP |
| | AAEL011349 | SP55/SP218 |
| | AAEL001098 | CLIP-domain serine protease, putative |
| | AAEL003279 | CLIP-domain serine protease, putative |
| | AAEL006576 | CLIP-domain serine protease, putative |
| | AAEL009722 | CLIP-domain serine protease, putative |
| | AAEL014386 | CLIP-domain serine protease, putative |
| | AAEL015465 | CLIP-domain serine protease, putative |
| | AAEL007587 | CLIP-domain serine protease |
| | AAEL022578 | CLIP-domain serine protease |
| | AAEL015637 | Aaeg:CLIP23 |
| **CTL** | AAEL000283 | CTLMA16 |
| | AAEL029028 | CTL |
| | AAEL029020 | CTL |
| | AAEL029053 | CTL |
| | AAEL029039 | CTL |
| | AAEL009338 | CTL10 |
| | AAEL000533 | CTL16 |
| | AAEL000543 | CTLMA11 |
| | AAEL000556 | CTL25 |
| | AAEL000563 | CTLMA15 |

| | | |
|---|---|---|
| | AAEL002524 | CTL24 |
| | AAEL022136 | CTL6 |
| | AAEL005482 | CTL18 |
| | AAEL005641 | CTLGA5 |
| | AAEL008299 | CTL11 |
| | AAEL008681 | CTL12 |
| | AAEL021200 | CTLSE1 |
| | AAEL009209 | CTLGA6 |
| | AAEL018207 | CTL8 |
| | AAEL011070 | CTLGA3 |
| | AAEL011078 | CTLGA1 |
| | AAEL011079 | CTLMA10 |
| | AAEL029068 | CTL5 |
| | AAEL011404 | CTL19 |
| | AAEL011407 | CTL20 |
| | AAEL011408 | CTL21 |
| | AAEL011453 | CTL14 |
| | AAEL011455 | CTLMA12 |
| | AAEL011612 | CTLMA6 |
| | AAEL011621 | CTLMA13 |
| | AAEL012353 | CTL15 |
| | AAEL018265 | CTL9 |
| | AAEL013853 | CTLGA2 |
| | AAEL014382 | CTLMA14 |
| | AAEL019633 | CTLGA9 |
| | AAEL025802 | CTLD-S |
| | AAEL025598 | CTLD-S |
| | AAEL022823 | mosGCTL-11 |
| | AAEL026955 | CTLD-S |
| | AAEL023353 | CTLD-S |
| | AAEL014384 | CTLD-S |
| | AAEL027215 | CTLD-S |
| | AAEL027443 | CTLD-S |
| | AAEL011622 | CLSP1(mosGCTL-31) |
| | AAEL006825 | CTLD-E |
| | AAEL014357 | CTLD-X |
| | AAEL006958 | CTLD-X |
| | AAEL001935 | CTL-like protein 1 |
| **FREP** | AAEL000508 | FREP15 |
| | AAEL000726 | FREP20 |
| | AAEL000749 | FREP22 |
| | AAEL001713 | FREP2 |
| | AAEL002713 | FREP31 |
| | AAEL003156 | FREP28 |

| | | |
|---|---|---|
| | AAEL003294 | FREP3 |
| | AAEL006691 | FREP |
| | AAEL006699 | FREP34 |
| | AAEL006702 | FREP33 |
| | AAEL006704 | FREP18 |
| | AAEL007942 | FREP14 |
| | AAEL008104 | FREB23 |
| | AAEL009384 | FREP5 |
| | AAEL009723 | FREP11 |
| | AAEL010117 | FREP35 |
| | AAEL011009 | FREP13 |
| | AAEL011633 | FREP16 |
| | AAEL011634 | FREP12 |
| | AAEL013417 | FREP24 |
| | AAEL013506 | FREP29 |
| | AAEL014432 | FREP25 |
| | AAEL021180 | FREP26 |
| | AAEL025451 | FREP27 |
| | AAEL014773 | FREP |
| | AAEL019868 | FREP |
| | AAEL020192 | FREP |
| | AAEL023956 | FREP |
| | AAEL025744 | FREP |
| | AAEL028175 | FREP |
| | AAEL011007 | FREP |
| **Galectin** | AAEL003541 | GALE1 |
| | AAEL003840 | GALE11 |
| | AAEL003844 | GALE5 |
| | AAEL004196 | GALE3 |
| | AAEL005293 | GALE8A |
| | AAEL009842 | GALE12 |
| | AAEL009850 | GALE14 |
| | AAEL012003 | GALE6B |
| | AAEL012135 | GALE2 |
| | AAEL026564 | Galectin |
| **IMD pathway** | AAEL010083 | Imd |
| | AAEL027860 | Caspar |
| | AAEL001932 | FADD |
| | AAEL012510 | IKK2 |
| | AAEL018130 | TAK1 |
| | AAEL003245 | IKK1 |
| | AAEL026170 | TAB2 |
| | AAEL003371 | Beta-TrCP [KO:K03362] |

| | | |
|---|---|---|
| | AAEL003103 | Ubiquitin-conjugating enzyme E2-17 kDa [KO:K06689] [EC:2.3.2.23] |
| | AAEL011873 | Ubiquitin-conjugating enzyme E2 variant 1 [KO:K10704] |
| | AAEL002118 | Ubiquitin-conjugating enzyme E2 N [KO:K10580] [EC:2.3.2.23] |
| | AAEL028161 | Mitogen-activated protein kinase kinase kinase 4 isoform X1 [KO:K04428] [EC:2.7.11.25] |
| | AAEL023782 | Mitogen-activated protein kinase kinase kinase 4 isoform X1 [KO:K04428] [EC:2.7.11.25] |
| | AAEL001622 | Dual specificity mitogen-activated protein kinase kinase 3 isoform X1 [KO:K04432] [EC:2.7.12.2] |
| | AAEL008379 | Mitogen-activated protein kinase 14B isoform X2 [KO:K04441] [EC:2.7.11.24] |
| | AAEL013261 | Cyclic AMP-dependent transcription factor ATF-2 [KO:K04450] |
| | AAEL021333 | Dual specificity mitogen-activated protein kinase kinase hemipterous isoform X1 [KO:K04431] [EC:2.7.12.2] |
| | AAEL003505 | Transcription factor AP-1 isoform X2 [KO:K04448] |
| | AAEL008953 | Transcription factor kayak isoform X3 [KO:K09031] |
| | AAEL011563 | Ankyrin-3 isoform X1 [KO:K10380] |
| | AAEL013466 | Ankyrin-3 isoform X1 [KO:K10380] |
| **Jak-stat** | AAEL012471 | DOME |
| | AAEL012553 | HOP |
| | AAEL020559 | STAT |
| | AAEL019728 | SOCS |
| | AAEL009822 | GPRMGL5 |
| | AAEL009645 | Hypotheical protein |
| **Lysozymes** | AAEL003712 | LYSC10 |
| | AAEL003723 | LYSC11 |
| | AAEL019435 | LYSC6 |
| | AAEL015404 | LYSC7B |
| | AAEL017132 | LYSC4 |
| **LRR/LRIM** | AAEL010125 | LRIM17 |
| | AAEL010132 | LRIM3 |
| | AAEL012255 | LRIM13 |
| | AAEL001420 | LRIM8 |
| | AAEL001401 | LRIM10A |
| | AAEL012092 | Leucine-rich repeat |
| | AAEL010772 | Leucine-rich repeat-containing protein |
| | AAEL012093 | Leucine-rich transmembrane protein |
| | AAEL005734 | Leucine-rich transmembrane protein |
| | AAEL002295 | Leucine-rich transmembrane protein |
| | AAEL003597 | Leucine-rich transmembrane protein |
| | AAEL000243 | Leucine-rich transmembrane protein |
| | AAEL006026 | Leucine rich protein, putative |
| | AAEL007565 | Leucine rich protein, putative |
| | AAEL003554 | Leucine rich repeat protein |

| | | |
|---|---|---|
| | AAEL004711 | Testis specific leucine rich repeat protein |
| | AAEL004466 | Leucine-rich immune protein (Coil-less) |
| | AAEL002615 | Leucine-rich transmembrane protein |
| | AAEL006377 | LRIM31 |
| | AAEL007785 | Leucine-rich transmembrane protein |
| | AAEL006920 | LRIM20 |
| | AAEL012763 | LRIM24 |
| | AAEL000762 | LRIM19 |
| | AAEL007224 | LRIM22 |
| | AAEL001402 | LRIM10B |
| | AAEL001417 | LRIM7 |
| | AAEL009792 | LRIM25 |
| | AAEL007103 | LRIM15 |
| | AAEL010128 | LRIM4 |
| | AAEL007778 | Leucine-rich transmembrane protein |
| | AAEL012086 | LRIM1 |
| | AAEL012538 | LRIM6 |
| | AAEL008658 | LRIM16 |
| | AAEL010656 | LRIM12 |
| | AAEL001649 | Leucine aminopeptidase |
| | AAEL007363 | Leucine-rich transmembrane protein |
| | AAEL000108 | Leucine aminopeptidase |
| | AAEL006975 | Leucine aminopeptidase |
| | AAEL000424 | Leucine aminopeptidase |
| | AAEL012767 | LRIM5 |
| | AAEL003262 | Leucine-rich transmembrane protein |
| | AAEL004773 | Leucine carboxyl methyltransferase |
| | AAEL006797 | F-box/Leucine rich repeat protein |
| | AAEL001414 | LRIM9 |
| | AAEL009894 | LRIM21 |
| | AAEL011387 | Leucine-rich repeat |
| | AAEL002307 | Leucine-rich transmembrane protein |
| | AAEL007442 | F-box/leucine rich repeat protein |
| | AAEL010793 | F-box/leucine rich repeat protein |
| | AAEL005762 | Leucine-rich transmembrane proteins |
| | AAEL001766 | Leucine-rich transmembrane proteins |
| | AAEL010286 | Leucine-rich transmembrane protein |
| | AAEL003713 | Leucine-rich transmembrane protein |
| | AAEL003408 | Leucine-rich transmembrane protein |
| | AAEL012911 | LRIM18 |
| | AAEL000925 | Leucine-zipper-like transcriptional regulator 1 (LZTR-1) |
| **ML** | AAEL009531 | Niemann-Pick |
| | AAEL004120 | ML1 |
| | AAEL006854 | ML13 |

| | | |
|---|---|---|
| | AAEL009553 | ML9B |
| | AAEL009555 | ML15A |
| | AAEL009556 | ML15B |
| | AAEL015140 | ML16 |
| | AAEL009760 | ML21 |
| | AAEL026174 | ML9B |
| | AAEL012064 | ML2 |
| | AAEL007592 | ML20B |
| | AAEL007591 | ML26A |
| | AAEL001654 | ML30 |
| | AAEL019611 | ML31 |
| | AAEL001634 | ML32 |
| | AAEL001650 | ML33 |
| | AAEL015136 | ML6 |
| | AAEL015137 | ML20 |
| | AAEL015139 | ML22A |
| | AAEL008492 | DVRF1 |
| | AAEL019883 | ML1 |
| | AAEL009956 | Aaeg:ML18 |
| **PGRP** | AAEL017056 | PGRPS4 |
| | AAEL019745 | PGPPLD putative |
| | AAEL009474 | PGRPS1 |
| | AAEL010171 | PGRPLB |
| | AAEL012380 | PGRPLA |
| | AAEL027982 | PGRPLE |
| | AAEL014640 | PGRPLC |
| | AAEL007039 | Peptidoglycan recognition protein (short) |
| | AAEL021026 | PGRP |
| **ROS** | AAEL000342 | PRDX |
| | AAEL020747 | PRDX |
| | AAEL026038 | PRDX |
| | AAEL004386 | pxt |
| | AAEL000495 | GPXH3 |
| | AAEL002309 | TPX4 |
| | AAEL002354 | HPX5 |
| | AAEL019408 | TPX2 |
| | AAEL004388 | HPX8A |
| | AAEL004390 | HPX8B |
| | AAEL006014 | HPX1 |
| | AAEL007563 | DUOX |
| | AAEL008397 | GPXH2 |
| | AAEL009051 | TPX5 |
| | AAEL012069 | GPXH1 |
| | AAEL013171 | HPX2 |

| | | |
|---|---|---|
| | AAEL013528 | TPX1 |
| | AAEL025567 | TPX3 |
| | AAEL019639 | HPX3 |
| | AAEL011941 | Oxidase/peroxidase |
| | AAEL000507 | Peroxidase |
| **PPO** | AAEL011763 | PPO3 |
| | AAEL011764 | PPO10 |
| | AAEL013492 | PPO5 |
| | AAEL013493 | PPO7 |
| | AAEL013496 | PPO8 |
| | AAEL014544 | PPO1 |
| | AAEL013501 | PPO4 |
| | AAEL015113 | PPO2 |
| | AAEL015116 | PPO1 |
| | AAEL020579 | PPO9 |
| **Relish** | AAEL007696 | REL1A |
| | AAEL006930 | REL1B |
| | AAEL007624 | REL2 |
| **SCR** | AAEL000227 | SCRB8 |
| | AAEL000234 | SCRB7 |
| | AAEL000256 | SCRB9 |
| | AAEL001914 | SCRAC1 |
| | AAEL002741 | SCRB6 |
| | AAEL005374 | SCRB1 |
| | AAEL005979 | SCRB3 |
| | AAEL006355 | SCRC1 |
| | AAEL006361 | SCRC2 |
| | AAEL008370 | SCRB17 |
| | AAEL009192 | SCRASP1 |
| | AAEL009420 | SCRBQ1 |
| | AAEL009423 | SCRBQ2 |
| | AAEL009432 | SCRBQ3 |
| | AAEL010655 | SCRBSP2 |
| | AAEL011222 | SCRB5 |
| | AAEL019436 | SCRB16 |
| | AAEL022263 | SCRAL1 |
| | AAEL027694 | SCRASP3 |
| | AAEL027927 | SCR |
| **SOD** | AAEL004823 | MNSOD1 |
| | AAEL005108 | MNSOD2 |
| | AAEL006271 | CUSOD2 |
| | AAEL019759 | CUSOD1 |
| | AAEL019761 | CUSOD1 |
| | AAEL019937 | SOD-Cu-Zn |

| | | |
|---|---|---|
| | AAEL019938 | CUSOD4 |
| | AAEL025388 | SOD7 |
| **SPZ** | AAEL001435 | SPZ2 |
| | AAEL001929 | SPZ5 |
| | AAEL007897 | SPZ4 |
| | AAEL008596 | SPZ3A |
| | AAEL012164 | SPZ6 |
| | AAEL013433 | SPZ1C |
| | AAEL013434 | SPZ1B |
| **Serpins** | AAEL002699 | SRPN7 |
| | AAEL002704 | SRPN |
| | AAEL002720 | SRPN20 |
| | AAEL002730 | SRPN21 |
| | AAEL002731 | SRPN14 |
| | AAEL003182 | SRPN26 |
| | AAEL003653 | SRPN12 |
| | AAEL003697 | SRPN17 |
| | AAEL005665 | SRPN3 |
| | AAEL006137 | SRPN19 |
| | AAEL007420 | SRPN25 |
| | AAEL007765 | SRPN10 |
| | AAEL008364 | SRPN9 |
| | AAEL010769 | SRPN6 |
| | AAEL011777 | SRPN8 |
| | AAEL012378 | SRPN |
| | AAEL013936 | SRPN4 |
| | AAEL014078 | SRPN2 |
| | AAEL014079 | SRPN1 |
| | AAEL014138 | SRPN16 |
| | AAEL014141 | SRPN5 |
| | AAEL017249 | SRPN24 |
| | AAEL020823 | SRPN11 |
| | AAEL024006 | SRPN |
| | AAEL028034 | SRPN |
| **siRNA** | AAEL000293 | TSN |
| | AAEL001317 | RM62A |
| | AAEL001612 | DCR1 |
| | AAEL001769 | RM62B |
| | AAEL002083 | RM62C |
| | AAEL002351 | RM62D |
| | AAEL004978 | RM62E |
| | AAEL006287 | PIWI7 |
| | AAEL006794 | DCR2 |
| | AAEL007698 | PIWI4 |

| | | |
|---|---|---|
| | AAEL007823 | PIWI |
| | AAEL008073 | VIG |
| | AAEL008098 | PIWI2 |
| | AAEL008592 | DROSHA |
| | AAEL008687 | LOQS |
| | AAEL008738 | RM62F |
| | AAEL009326 | FMR1 |
| | AAEL010402 | RM62G |
| | AAEL010787 | DEAD box ATP-dependent RNA helicase |
| | AAEL011663 | Aa-ago2 |
| | AAEL011753 | R2D2 |
| | AAEL012410 | AGO1B |
| | AAEL013233 | PIWI5 |
| | AAEL013277 | PIWI6 |
| | AAEL013692 | PIWI3 |
| | AAEL013985 | RM62I |
| | AAEL023716 | SPNE |
| | AAEL017251 | AGO2 |
| | AAEL019460 | Combined Dicer-1 and ARM |
| | AAEL021519 | PASHA |
| **TEP** | AAEL000087 | TEP3 |
| | AAEL001794 | TEP5 |
| | AAEL004725 | TEP |
| | AAEL005432 | TEP |
| | AAEL008607 | TEP |
| | AAEL009266 | c4b-binding protein beta chain |
| | AAEL012267 | TEP1 |
| | AAEL021904 | TEP |
| | AAEL001163 | TEP |
| | AAEL017023 | TEP |
| | AAEL025334 | TEP |
| **Toll** | AAEL000057 | TOLL5B |
| | AAEL000633 | TOLL8 |
| | AAEL000671 | TOLL6 |
| | AAEL000709 | CACT |
| | AAEL002583 | TOLL7 |
| | AAEL004000 | TOLL10 |
| | AAEL005075 | Ecsit |
| | AAEL006571 | PELLE |
| | AAEL007619 | TOLL5A |
| | AAEL007642 | TUBE |
| | AAEL007768 | MYD |
| | AAEL009551 | TOLL11 |
| | AAEL013441 | TOLL9A |

| | |
|---|---|
| AAEL015018 | Toll |
| AAEL026297 | TOLL1A |
| AAEL028236 | TRAF6 |

Supplementary Table 6. Non-classical immune genes.

| Gene family | Gene ID | Name/Description |
|---|---|---|
| **Trypsin** | AAEL009680 | Chymotrypsin |
| | AAEL001693 | Female-specific chymotrypsin |
| | AAEL006414 | Trypsin |
| | AAEL006430 | Trypsin |
| | AAEL006429 | Trypsin |
| | AAEL010202 | Trypsin |
| | AAEL013703 | Trypsin |
| | AAEL006425 | Trypsin |
| | AAEL013715 | Trypsin |
| | AAEL007601 | Trypsin |
| | AAEL010196 | Trypsin |
| | AAEL013623 | Trypsin |
| | AAEL007992 | Trypsin |
| | AAEL007818 | Trypsin 3A1 Precursor |
| | AAEL013712 | Trypsin 5G1 Precursor |
| | AAEL008079 | Trypsin-alpha |
| | AAEL006403 | Trypsin-beta |
| | AAEL005596 | Trypsin-epsilon |
| | AAEL008080 | Trypsin-eta |
| | AAEL013628 | Trypsin-eta |
| | AAEL005604 | Trypsin-epsilon, putative |
| | AAEL006365 | Trypsin-alpha, putative |
| | AAEL006368 | Trypsin-beta, putative |
| | AAEL006382 | Trypsin-eta, putative |
| | AAEL008097 | Trypsin-eta, putative |
| | AAEL011230 | Chymotrypsin, putative |
| | AAEL011882 | Trypsin-zeta, putative |
| | AAEL006418 | Trypsin |
| | AAEL006903 | Trypsin |
| | AAEL015638 | Trypsin |
| | AAEL000203 | Trypsin |
| | AAEL011553 | Trypsin |
| | AAEL012852 | Trypsin |
| | AAEL004996 | Trypsin |
| | AAEL005609 | Trypsin |
| | AAEL005607 | Trypsin |
| | AAEL005616 | Trypsin |
| | AAEL005611 | Trypsin |
| | AAEL013713 | Trypsin |
| | AAEL006421 | Trypsin |
| | AAEL006378 | Trypsin |
| | AAEL006376 | Trypsin |

| | | |
|---|---|---|
| | AAEL004543 | Trypsin |
| | AAEL002273 | Trypsin |
| | AAEL014579 | Trypsin |
| | AAEL007600 | Trypsin |
| | AAEL008214 | Trypsin |
| | AAEL006123 | Trypsin |
| | AAEL015432 | Trypsin |
| | AAEL006384 | Trypsin |
| | AAEL007102 | Trypsin |
| | AAEL007602 | Trypsin |
| | AAEL003308 | Trypsin |
| | AAEL006121 | Trypsin |
| | AAEL009853 | Trypsin |
| | AAEL017403 | TMOF |
| | AAEL013284 | LT1 |
| **Tubulin** | AAEL004172 | Tubulin alpha chain |
| | AAEL006642 | Tubulin alpha chain |
| | AAEL002848 | Tubulin beta chain |
| | AAEL013229 | Tubulin alpha chain |
| | AAEL019894 | Tubulin beta chain |
| | AAEL006179 | Tubulin alpha chain |
| | AAEL012101 | Tubulin alpha chain |
| | AAEL004939 | Tubulin beta chain |
| | AAEL012843 | Tubulin gamma chain |
| | AAEL005084 | Tubulin beta chain |
| | AAEL002851 | Tubulin beta chain |
| | AAEL005052 | Tubulin beta chain |
| | AAEL012424 | Tubulin alpha chain |
| | AAEL002135 | Tubulin-specific chaperone b (tubulin folding cofactor b) |
| | AAEL003546 | Gamma-tubulin complex component 4 (gcp-4) |
| | AAEL008465 | Gamma-tubulin complex component 3 (gcp-3) |
| | AAEL011067 | Tubulin-specific chaperone, putative |
| | AAEL005303 | Beta-tubulin cofactor d |
| | AAEL013903 | Gamma-tubulin complex component 2 (gcp-2) |
| | AAEL004440 | Tubulin-specific chaperone e |
| | AAEL008679 | Alpha-tubulin N-acetyltransferase |
| **Actin** | AAEL004646 | Actin |
| | AAEL005964 | Actin |
| | AAEL011317 | Actin |
| | AAEL004631 | Actin |
| | AAEL004616 | Actin |
| | AAEL009451 | Actin |
| | AAEL001673 | Actin |

| | | |
|---|---|---|
| | AAEL005961 | Actin |
| | AAEL003383 | Actin |
| | AAEL012310 | Actin |
| | AAEL004843 | Actin |
| | AAEL011197 | Actin |
| | AAEL011750 | Actin |
| | AAEL001951 | Act4 |
| | AAEL003754 | Actin binding |
| | AAEL010664 | Actin binding protein, putative |
| | AAEL013661 | Actin binding protein, putative |
| | AAEL002953 | Actin 3 isoform, putative |
| | AAEL011972 | Actin binding protein, putative |
| | AAEL007660 | Suppressor of actin (sac) |
| | AAEL012519 | Actin binding protein, putative |
| | AAEL008102 | Actin binding protein, putative |
| | AAEL004371 | SWI/SNF related matrix associated actin dependent regulator of chromatin subfamily B member 1 |
| | AAEL002822 | Arp5 |
| | AAEL002184 | F-actin capping protein beta subunit |
| | AAEL007546 | Actin-related protein 2/3 complex subunit 1A |
| | AAEL001928 | Actin-1 |
| | AAEL013778 | F-actin capping protein alpha |
| | AAEL027716 | Actin-related protein 2/3 complex subunit 3 |
| | AAEL003701 | Actin-binding protein ipp |
| | AAEL010762 | Arp8 |
| **Myosin** | AAEL000596 | Myosin |
| | AAEL012449 | Myosin x |
| | AAEL011905 | Myosin i |
| | AAEL008610 | Myosin vii |
| | AAEL004227 | Myosin VI |
| | AAEL011436 | Myosin xv |
| | AAEL009991 | Myosin iii |
| | AAEL012543 | Myosin motor, putative |
| | AAEL000382 | Myosin motor, putative |
| | AAEL021838 | Myosin heavy chain |
| | AAEL003676 | Myosin I homologue, putative |
| | AAEL007439 | Myosin light chain 1, |
| | AAEL012207 | Myosin light chain 1, |
| | AAEL007632 | Myosin light chain kinase |
| | AAEL012926 | Unconventional Myosin 95e isoform |
| | AAEL001068 | Myosin light chain 2V, putative |
| | AAEL004750 | Nonmuscle Myosin heavy chain-A, putative |
| | AAEL011428 | Fast myosin heavy chain HCIII, putative |
| | AAEL002572 | Myosin regulatory light chain 2 (mlc-2) |

| | | |
|---|---|---|
| | AAEL008921 | Myosin regulatory light chain 2 smooth muscle |
| | AAEL001411 | Myosin heavy chain, nonmuscle or smooth muscle |
| | AAEL005656 | Myosin heavy chain, nonmuscle or smooth muscle |
| | AAEL001220 | CK |
| **Heat shock proteins** | AAEL011708 | Heat shock protein |
| | AAEL000301 | Heat shock protein |
| | AAEL014843 | Heat shock protein |
| | AAEL013161 | Heat shock protein, putative |
| | AAEL001052 | Heat shock protein, putative |
| | AAEL017976 | Heat shock protein HSP70 |
| | AAEL019403 | Heat shock cognate 70 |
| | AAEL017975 | Heat shock protein HSP70 |
| | AAEL013350 | Heat shock protein 26kD, putative |
| | AAEL010546 | Heat shock factor binding protein, putative |
| | AAEL004148 | Heat shock protein 70 (hsp70)-interacting protein |
| | AAEL001952 | 28 kDa heat- and acid-stable phosphoprotein (PDGF-associated protein), putative |
| **Cell proliferation** | AAEL008069 | NOTCH |
| | AAEL011396 | DELTA |
| **Cytochrome P450** | AAEL009132 | CYP6Y3 |
| | AAEL014615 | CYP9J23 |
| | AAEL001292 | CYP9M7 |
| | AAEL014603 | CYP9J30 |
| | AAEL007816 | CYP4D23 |
| | AAEL002633 | CYP9J31 |
| | AAEL014613 | CYP9J24 |
| | AAEL009133 | CYP6N14 |
| | AAEL005700 | CYP325X4 |
| | AAEL003890 | CYP |
| | AAEL014619 | CYP9J22 |
| | AAEL011463 | CYP |
| | AAEL009591 | CYP9M8 |
| | AAEL006984 | CYP6AG5 |
| | AAEL008638 | CYP49A1 |
| | AAEL010151 | CYP6N16 |
| | AAEL007808 | CYP4D39 |
| | AAEL004941 | CYP6AK1 |
| | AAEL010158 | CYP6N17 |
| | AAEL002043 | CYP305A5 |
| | AAEL014610 | CYP9J29 |
| | AAEL001960 | CYP |
| | AAEL005771 | CYP325K2 |

| | |
|---|---|
| AAEL017539 | CYP6BY1 |
| AAEL005788 | CYP325K3 |
| AAEL006992 | CYP6AG6 |
| AAEL003763 | CYP329B1 |
| AAEL004054 | CYP4G36 |
| AAEL014605 | CYP9J9 |
| AAEL003748 | CYP9AE1 |
| AAEL019911 | CYP325S2 |
| AAEL011850 | CYP315A1 |
| AAEL014893 | CYP6BB2 |
| AAEL009125 | CYP6M10 |
| AAEL009128 | CYP6M6 |
| AAEL017297 | CYP6M9 |
| AAEL009762 | CYP307A1 |
| AAEL009130 | CYP6Z7 |
| AAEL014891 | CYP6P12 |
| AAEL007807 | CYP4D38 |
| AAEL012770 | CYP325N1 |
| AAEL014594 | CYP301A1 |
| AAEL006257 | CYP325Y1 |
| AAEL007798 | CYP4K3 |
| AAEL013554 | CYP4J14 |
| AAEL002638 | CYP9J6 |
| AAEL000357 | CYP325S3 |
| AAEL009138 | CYP6N11 |
| AAEL011761 | CYP325M5 |
| AAEL004870 | CYP18A1 |
| AAEL014411 | CYP304B3 |
| AAEL009117 | CYP6M5 |
| AAEL002005 | CYP12F6 |
| AAEL003399 | CYP4H30 |
| AAEL002031 | CYP12F7 |
| AAEL012761 | CYP325T2 |
| AAEL000320 | CYP325T1 |
| AAEL009127 | CYP6M11 |
| AAEL000340 | CYP |
| AAEL009129 | CYP6Z9 |
| AAEL007812 | CYP4H32 |
| AAEL005695 | CYP325X1 |
| AAEL010154 | CYP4AR2 |
| AAEL009124 | CYP6N12 |
| AAEL007473 | CYP6AH1 |
| AAEL007795 | CYP4D37 |
| AAEL009123 | CYP6Z6 |
| AAEL012765 | CYP325M3 |
| AAEL013556 | CYP4J15 |

| | |
|---|---|
| AAEL001312 | CYP9M6 |
| AAEL006784 | CYP9J17 |
| AAEL011770 | CYP325L1 |
| AAEL006827 | CYP12F8 |
| AAEL009120 | CYP6S3 |
| AAEL003380 | CYP4H28 |
| AAEL009018 | CYP |
| AAEL005696 | CYP325X2 |
| AAEL005775 | CYP325R1 |
| AAEL018028 | CYP325Y3 |
| AAEL014413 | CYP304C1 |
| AAEL014614 | CYP |
| AAEL017136 | CYP325V1 |
| AAEL014609 | CYP9J26 |
| AAEL012766 | CYP325G2 |
| AAEL017215 | CYP325U1 |
| AAEL005006 | CYP6CD1 |
| AAEL012266 | CYP4C38 |
| AAEL009131 | CYP6Z8 |
| AAEL012769 | CYP325M2 |
| AAEL006824 | CYP |
| AAEL012772 | CYP325G3 |
| AAEL014019 | CYP4J16 |
| AAEL006815 | CYP9J16 |
| AAEL007830 | CYP4H29 |
| AAEL006795 | CYP9J15 |
| AAEL009121 | CYP6N9 |
| AAEL002085 | CYP4H31 |
| AAEL012762 | CYP325N2 |
| AAEL007010 | CYP6AG4 |
| AAEL006989 | CYP6AG7 |
| AAEL014618 | CYP |
| AAEL014830 | CYP |
| AAEL014678 | CYP6F2 |
| AAEL010946 | CYP314A1 |
| AAEL008889 | CYP6AL1 |
| AAEL006805 | CYP9J2 |
| AAEL009126 | CYP6N6 |
| AAEL008018 | CYP4C51 |
| AAEL014604 | CYP |
| AAEL013798 | CYP4H33 |
| AAEL009122 | CYP |
| AAEL000338 | CYP325E3 |
| AAEL007815 | CYP4D24 |
| AAEL000326 | CYP325S1 |
| AAEL014684 | CYP6F3 |

| | | |
|---|---|---|
| | AAEL009656 | CYP6AL3 |
| | AAEL001807 | CYP9M9 |
| | AAEL014412 | CYP304B2 |
| | AAEL006058 | CYP325Q2 |
| | AAEL001320 | CYP9M4 |
| | AAEL014924 | CYP |
| | AAEL012144 | CYP303A1 |
| | AAEL019910 | CYP325S2 |
| | AAEL007024 | CYP6AG3 |
| | AAEL009137 | CYP6N13 |
| | AAEL014890 | CYP6CC1 |
| | AAEL008017 | CYP4C50 |
| | AAEL006044 | CYP325Q1 |
| | AAEL014208 | CYP |
| | AAEL014617 | CYP9J28 |
| **Small Heat shock proteins** | AAEL013348 | Lethal (2)essential for life protein, l2efl |
| | AAEL013344 | Lethal (2)essential for life protein, l2efl |
| | AAEL013351 | Lethal (2)essential for life protein, l2efl |
| | AAEL013338 | Lethal (2)essential for life protein, l2efl |
| | AAEL013349 | Lethal (2)essential for life protein, l2efl |
| | AAEL013352 | Lethal (2)essential for life protein, l2efl |
| | AAEL013341 | Lethal (2)essential for life protein, l2efl |
| | AAEL013346 | Lethal (2)essential for life protein, l2efl |
| | AAEL013347 | Lethal (2)essential for life protein, l2efl |
| | AAEL010659 | Lethal (2)essential for life protein, l2efl |
| **Salivary proteins** | AAEL004407 | Allergen, putative |
| | AAEL004199 | Allergen |
| | AAEL003057 | Allergen |
| | AAEL010235 | Allergen |
| | AAEL000793 | Venom allergen |
| | AAEL002693 | Venom allergen |
| | AAEL005531 | Venom allergen |
| | AAEL013406 | Venom allergen |
| | AAEL011798 | Allergen, putative |
| | AAEL005997 | Allergen, putative |
| | AAEL006524 | Venom allergen |
| | AAEL011797 | Venom allergen |
| | AAEL002476 | Venom allergen |
| | AAEL003053 | Allergen, putative |
| | AAEL026620 | Allergen, putative |
| | AAEL002682 | Venom allergen |
| | AAEL009239 | Venom allergen |
| | AAEL027045 | Allergen, putative |
| | AAEL010269 | Venom allergen |
| | AAEL026087 | D7 |

| | | |
|---|---|---|
| | AAEL006347 | APY |
| | AAEL002726 | D7 protein, putative |
| | AAEL008620 | D7 protein, putative |
| | AAEL006424 | D7 |
| | AAEL006417 | D7 protein |
| **Lachesin** | AAEL009295 | Lachesin |
| | AAEL004992 | Lachesin, putative |
| | AAEL014334 | Lachesin, putative |
| | AAEL006478 | Lachesin, putative |
| | AAEL003966 | Lachesin |
| | AAEL000576 | Lachesin |
| **Sidestep proteins** | AAEL001227 | Sidestep protein |
| | AAEL008010 | Sidestep protein |
| | AAEL011989 | Sidestep protein |
| | AAEL019917 | Sidestep protein |
| | AAEL022485 | Sidestep protein |
| | AAEL023141 | Sidestep protein |
| **Vacuolar ATPase** | AAEL000291 | Vacuolar ATPases |
| | AAEL008787 | Vacuolar ATPases |
| | AAEL012113 | Vacuolar ATPases |
| | AAEL011025 | Vacuolar ATPases |
| | AAEL005798 | Vacuolar ATPases |
| | AAEL015594 | Vacuolar ATPases |
| | AAEL012035 | Vacuolar ATPases |
| | AAEL013302 | Vacuolar ATPases |
| | AAEL007184 | Vacuolar ATPases |
| | AAEL012819 | Vacuolar ATPases |
| | AAEL006516 | Vacuolar ATPases |
| | AAEL010819 | Vacuolar ATPases |
| | AAEL014053 | Vacuolar ATPases |
| | AAEL003743 | Vacuolar ATPases |

Supplementary Table 7. Genes unmapped to DAVID cloud map.

| **3 dpi 18 ºC upregulated** | **3 dpi 18 ºC downregulated** | **3 dpi 28 ºC upregulated** | **3 dpi 28 ºC upregulated** | **3 dpi 28 ºC upregulated** |
|---|---|---|---|---|
| AAEL020330 | AAEL024221 | AAEL025079 | AAEL028635 | AAEL027610 |
| AAEL019751 | AAEL021086 | AAEL020330 | AAEL027655 | AAEL022876 |
| AAEL027238 | AAEL019650 | AAEL019578 | AAEL021257 | AAEL027362 |
| AAEL023745 | AAEL021576 | AAEL029047 | AAEL027493 | AAEL020033 |
| AAEL018301 | AAEL024146 | AAEL026440 | AAEL026008 | AAEL026843 |
| AAEL018304 | AAEL023395 | AAEL025125 | AAEL024175 | AAEL027514 |
| AAEL026878 | AAEL019698 | AAEL024669 | AAEL026300 | |
| AAEL023999 | AAEL021513 | AAEL022059 | AAEL026031 | |
| AAEL022900 | AAEL024475 | AAEL027093 | AAEL024540 | |
| AAEL026300 | AAEL023431 | AAEL020078 | AAEL022600 | |
| AAEL025532 | AAEL019834 | AAEL025894 | AAEL024112 | |
| AAEL026868 | AAEL024926 | AAEL022674 | AAEL028247 | |
| AAEL021302 | AAEL021138 | AAEL027106 | AAEL026819 | |
| AAEL023746 | AAEL023844 | AAEL021929 | AAEL022427 | |
| AAEL022253 | AAEL018216 | AAEL027700 | AAEL026447 | |
| AAEL024512 | AAEL024064 | AAEL019773 | AAEL026833 | |
| AAEL022363 | AAEL023158 | AAEL024838 | AAEL026537 | |
| AAEL024540 | AAEL023753 | AAEL026751 | AAEL026041 | |
| AAEL025894 | AAEL020477 | AAEL025126 | AAEL029046 | |
| AAEL019935 | AAEL024122 | AAEL026175 | AAEL025750 | |
| AAEL021595 | AAEL028236 | AAEL018351 | AAEL019849 | |
| AAEL023591 | AAEL027166 | AAEL025392 | AAEL018241 | |
| AAEL027610 | AAEL018125 | AAEL022387 | AAEL019438 | |
| AAEL018241 | AAEL024149 | AAEL027699 | AAEL027829 | |
| AAEL026519 | AAEL025667 | AAEL023395 | AAEL026603 | |
| AAEL025531 | AAEL026023 | AAEL019537 | AAEL023729 | |
| AAEL017975 | AAEL019588 | AAEL019528 | AAEL025332 | |
| AAEL021614 | AAEL025199 | AAEL021072 | AAEL022079 | |
| AAEL026194 | AAEL023874 | AAEL023882 | AAEL023999 | |
| AAEL020777 | AAEL024161 | AAEL021302 | AAEL021795 | |
| AAEL026751 | AAEL021180 | AAEL027019 | AAEL025839 | |
| AAEL017976 | AAEL028002 | AAEL020603 | AAEL023560 | |
| AAEL020575 | | AAEL022829 | AAEL027270 | |
| AAEL025126 | | AAEL025552 | AAEL029031 | |
| AAEL019637 | | AAEL019868 | AAEL026215 | |
| AAEL019995 | | AAEL019844 | AAEL019751 | |
| AAEL028247 | | AAEL017975 | AAEL028221 | |
| AAEL020957 | | AAEL020340 | AAEL023348 | |
| AAEL026833 | | AAEL021016 | AAEL027188 | |
| AAEL023321 | | AAEL019893 | AAEL021583 | |
| AAEL022079 | | AAEL024560 | AAEL027632 | |
| AAEL018189 | | AAEL022334 | AAEL019463 | |
| AAEL020236 | | AAEL019564 | AAEL017976 | |
| AAEL026215 | | AAEL024038 | AAEL024512 | |
| AAEL022059 | | AAEL025531 | AAEL023321 | |
| AAEL024560 | | AAEL021982 | AAEL026531 | |
| AAEL019902 | | AAEL019767 | AAEL023591 | |
| AAEL027096 | | AAEL022253 | AAEL019684 | |
| AAEL026008 | | AAEL024468 | AAEL023243 | |

| 3 dpi 28 ºC downregulated | 3 dpi 32 ºC upregulated | 3 dpi 32 ºC downregulated |
|---|---|---|
| AAEL019677 | AAEL020330 | AAEL024207 |
| AAEL024298 | AAEL021012 | |
| AAEL022506 | AAEL024560 | |
| AAEL026775 | AAEL023321 | |
| AAEL028022 | AAEL017976 | |
| AAEL026260 | | |
| AAEL018189 | | |
| AAEL025401 | | |
| AAEL024022 | | |
| AAEL020175 | | |
| AAEL026241 | | |
| AAEL021303 | | |
| AAEL022382 | | |
| AAEL025549 | | |
| AAEL022900 | | |

| 7 dpi 18 ºC upregulated | 7 dpi 18 ºC downregulated | 7 dpi 28 ºC upregulated | 7 dpi 28 ºC upregulated | 7 dpi 28 ºC downregulated |
|---|---|---|---|---|
| AAEL026008 | AAEL025334 | AAEL019955 | AAEL024337 | AAEL019889 |
| AAEL022059 | AAEL018189 | AAEL027610 | AAEL028635 | AAEL019885 |
| AAEL022253 | AAEL023617 | AAEL020330 | AAEL021929 | AAEL024757 |
| AAEL025332 | AAEL020035 | AAEL021257 | AAEL026214 | AAEL018234 |
| AAEL024597 | AAEL029039 | AAEL019658 | AAEL025531 | AAEL024284 |
| AAEL024257 | | AAEL023524 | AAEL026603 | AAEL020075 |
| AAEL026833 | | AAEL024512 | AAEL019463 | AAEL026107 |
| AAEL021072 | | AAEL020706 | AAEL026833 | AAEL021576 |
| AAEL020330 | | AAEL026481 | AAEL018334 | AAEL021308 |
| AAEL019704 | | AAEL022059 | AAEL020936 | AAEL019629 |
| AAEL020804 | | AAEL024269 | AAEL027545 | AAEL022124 |
| AAEL023321 | | AAEL025921 | AAEL025126 | AAEL021449 |
| AAEL019751 | | AAEL020502 | AAEL022775 | AAEL024830 |
| AAEL019899 | | AAEL029061 | AAEL026300 | |
| AAEL028247 | | AAEL027700 | AAEL024717 | |
| AAEL017976 | | AAEL026175 | AAEL021555 | |
| AAEL019684 | | AAEL019537 | AAEL019751 | |
| AAEL027802 | | AAEL024406 | AAEL018340 | |
| AAEL017975 | | AAEL019527 | AAEL022253 | |
| AAEL022079 | | AAEL027068 | AAEL022932 | |
| AAEL023002 | | AAEL021073 | AAEL024753 | |
| AAEL023591 | | AAEL024838 | AAEL023125 | |
| AAEL024512 | | AAEL025296 | AAEL018150 | |
| AAEL024038 | | AAEL019487 | AAEL021375 | |
| AAEL019688 | | AAEL021016 | AAEL022593 | |
| AAEL020028 | | AAEL025177 | AAEL017976 | |
| AAEL022829 | | AAEL026446 | AAEL018343 | |
| AAEL023395 | | AAEL029031 | AAEL022982 | |
| AAEL026848 | | AAEL022387 | AAEL020392 | |
| AAEL025731 | | AAEL021795 | AAEL027984 | |
| AAEL024887 | | AAEL028228 | AAEL019752 | |
| AAEL020092 | | AAEL022578 | AAEL021302 | |
| AAEL025894 | | AAEL023100 | AAEL023294 | |
| AAEL023555 | | AAEL028247 | AAEL025903 | |
| AAEL026031 | | AAEL019564 | AAEL023844 | |
| AAEL029030 | | AAEL019847 | AAEL022829 | |
| AAEL027610 | | AAEL021583 | AAEL023847 | |
| | | AAEL026843 | AAEL019917 | |
| | | AAEL024291 | AAEL026751 | |
| | | AAEL023591 | AAEL024583 | |
| | | AAEL023321 | AAEL022079 | |
| | | AAEL024540 | AAEL017975 | |
| | | AAEL024216 | | |
| | | AAEL019954 | | |
| | | AAEL026008 | | |
| | | AAEL022674 | | |
| | | AAEL021099 | | |
| | | AAEL020430 | | |
| | | AAEL027188 | | |
| | | AAEL021263 | | |

| 7 dpi 32ºC upregulated | 7 dpi 32ºC downregulated |
|---|---|
| AAEL020330 | AAEL025193 |
| AAEL027157 | AAEL023039 |
| AAEL027610 | AAEL022225 |
| AAEL023509 | AAEL025401 |
| AAEL022253 | AAEL018041 |
| AAEL019504 | AAEL020306 |
| AAEL028247 | AAEL021011 |
| AAEL019590 | AAEL024449 |
| AAEL023844 | AAEL026029 |
| AAEL026819 | AAEL020913 |
| AAEL024560 | AAEL019689 |
| AAEL023591 | AAEL026468 |
| AAEL018159 | AAEL024183 |
| AAEL027654 | AAEL018184 |
| AAEL025574 | |
| AAEL020502 | |
| AAEL022334 | |
| AAEL018103 | |
| AAEL022208 | |
| AAEL019641 | |
| AAEL019564 | |
| AAEL026161 | |
| AAEL022059 | |
| AAEL026537 | |
| AAEL017975 | |
| AAEL017976 | |
| AAEL027362 | |
| AAEL022589 | |
| AAEL019751 | |
| AAEL023999 | |
| AAEL021929 | |
| AAEL021302 | |
| AAEL022079 | |
| AAEL027829 | |
| AAEL022982 | |
| AAEL024179 | |
| AAEL026008 | |
| AAEL019602 | |
| AAEL026390 | |
| AAEL025126 | |
| AAEL026496 | |

Supplementary Table 8. LncRNA

| 3 dpi 18 ºC upregulated | 3 dpi 18 ºC downregulated | 3 dpi 28 ºC upregulated | 3 dpi 28 ºC downregulated |
|---|---|---|---|
| AAEL019751 | AAEL021086 | AAEL025079 | AAEL026775 |
| AAEL027238 | AAEL021576 | AAEL019578 | AAEL026260 |
| AAEL023999 | AAEL024146 | AAEL027093 | AAEL018189 |
| AAEL022900 | AAEL023395 | AAEL025894 | AAEL020175 |
| AAEL025532 | AAEL019698 | AAEL027106 | AAEL022382 |
| AAEL026868 | AAEL021513 | AAEL021929 | AAEL025549 |
| AAEL021302 | AAEL023431 | AAEL027700 | AAEL022900 |
| AAEL024540 | AAEL024926 | AAEL019773 | |
| AAEL025894 | AAEL021138 | AAEL025126 | |
| AAEL026519 | AAEL023158 | AAEL023395 | |
| AAEL021614 | AAEL024122 | AAEL019528 | |
| AAEL020777 | AAEL024149 | AAEL021072 | |
| AAEL020575 | AAEL024161 | AAEL023882 | |
| AAEL025126 | | AAEL021302 | |
| AAEL022079 | | AAEL022829 | |
| AAEL018189 | | AAEL025552 | |
| | | AAEL019844 | |
| | | AAEL020340 | |
| | | AAEL022334 | |
| | | AAEL024038 | |
| | | AAEL021982 | |
| | | AAEL027655 | |
| | | AAEL027493 | |
| | | AAEL024540 | |
| | | AAEL024112 | |
| | | AAEL026447 | |
| | | AAEL026537 | |
| | | AAEL026603 | |
| | | AAEL025332 | |
| | | AAEL022079 | |
| | | AAEL023999 | |
| | | AAEL027270 | |
| | | AAEL019751 | |
| | | AAEL021583 | |
| | | AAEL022876 | |
| | | AAEL027362 | |

| 3 dpi 32ºC upregulated, 3 dpi 32ºC downregulated |
|---|
| None |

| 7 dpi 18 ºC upregulated | 7 dpi 18 ºC downregulated | 7 dpi 28 ºC upregulated | 7 dpi 28 ºC downregulated | 7 dpi 32 ºC upregulated | 7 dpi 32 ºC downregulated |
|---|---|---|---|---|---|
| AAEL025332 | AAEL018189 | AAEL026481 | AAEL026107 | AAEL022334 | AAEL025193 |
| AAEL024257 | | AAEL024269 | AAEL021576 | AAEL022208 | AAEL023039 |
| AAEL021072 | | AAEL027700 | AAEL022124 | AAEL026161 | AAEL020306 |
| AAEL019704 | | AAEL025296 | | AAEL026537 | AAEL026029 |
| AAEL019751 | | AAEL023100 | | AAEL027362 | AAEL024183 |
| AAEL022079 | | AAEL021583 | | AAEL022589 | |
| AAEL024038 | | AAEL024540 | | AAEL019751 | |
| AAEL020028 | | AAEL019954 | | AAEL023999 | |
| AAEL022829 | | AAEL021929 | | AAEL021929 | |
| AAEL023395 | | AAEL026603 | | AAEL021302 | |
| AAEL024887 | | AAEL018334 | | AAEL022079 | |
| AAEL025894 | | AAEL025126 | | AAEL026390 | |
| | | AAEL019751 | | AAEL025126 | |
| | | AAEL022932 | | | |
| | | AAEL019752 | | | |
| | | AAEL021302 | | | |
| | | AAEL025903 | | | |
| | | AAEL022829 | | | |
| | | AAEL023847 | | | |
| | | AAEL019917 | | | |
| | | AAEL022079 | | | |
